# The two-component regulator WalKR provides an essential link between cell wall homeostasis with DNA replication in *Staphylococcus aureus*

**DOI:** 10.1101/2023.02.27.530350

**Authors:** Liam K. R. Sharkey, Romain Guerillot, Calum Walsh, Adrianna M. Turner, Jean Y. H. Lee, Stephanie L. Neville, Stephan Klatt, Sarah L. Baines, Sacha Pidot, Fernando J. Rossello, Torsten Seemann, Hamish McWilliam, Ellie Cho, Glen P. Carter, Benjamin P. Howden, Christopher A. McDevitt, Abderrahman Hachani, Timothy P. Stinear, Ian R. Monk

## Abstract

Among the 16 two-component systems (TCSs) in the opportunistic human pathogen *Staphylococcus aureus*, only WalKR is essential. Like orthologous systems in other Bacillota, *S. aureus* WalKR controls autolysins involved in peptidoglycan remodelling and is therefore intimately involved in cell division. However, despite the importance of WalKR in *S. aureus*, the basis for its essentiality is not understood and the regulon poorly defined. Here, we defined a consensus WalR DNA-binding motif and the direct WalKR regulon by using functional genomics, including ChIP-seq, with a panel of isogenic *walKR* mutants that had a spectrum of altered activities. Consistent with prior findings, the direct regulon includes multiple autolysin genes. However, this work also revealed that WalR directly regulates at least five essential genes involved in lipoteichoic acid synthesis (*ltaS*); translation *(rplK*); DNA compaction (*hup*); initiation of DNA replication (*dnaA, hup*); and purine nucleotide metabolism (*prs*). Thus, WalKR in *S. aureus* serves as a polyfunctional regulator that contributes to fundamental control over critical cell processes by co-ordinately linking cell wall homeostasis with purine biosynthesis, protein biosynthesis, and DNA replication. Collectively, our findings address the essentiality of this locus and highlight the importance of WalKR as a *bona fide* target for novel anti-staphylococcal therapeutics.

## Introduction

*Staphylococcus aureus* is an opportunistic pathogen that causes a wide range of hospital and community acquired infections. Antibiotic-resistant strains, notably methicillin resistant (MRSA) and vancomycin-intermediate (VISA) strains, are persistent problems, with last-line agents, such as vancomycin, linezolid, and daptomycin, commonly associated with treatment failure ^1, 2^. MRSA is a World Health Organization “priority antibiotic-resistant pathogen” for the research and development of new antibiotics. Mortality from serious *S. aureus* infection is high (20 – 50% of bacteraemias) ^3^, and the socioeconomic burden of *S. aureus* disease is substantial ^4^.

*S. aureus* encodes 16 core genome two-component systems (TCSs) that allow the bacterium to sense and respond to a range of stimuli, providing regulatory flexibility in changing environments. Of these 16 TCSs, only WalKR is essential for cell viability under laboratory conditions ^5, 6, 7, 8^. WalKR is a canonical TCS that is conserved amongst low-GC Gram-positive bacteria, comprised of a multi-domain transmembrane sensor histidine kinase (WalK) and DNA binding response regulator (WalR) of the OmpR family ^9, 10^. Upon activation by signal(s) WalK auto-phosphorylates a conserved histidine residue (H385) and subsequently transfers the phosphoryl group to a conserved aspartate residue (D53) in WalR. Phosphorylated WalR binds to promoter regions of genes within the WalKR regulon, operating as either a transcriptional activator or repressor.

WalKR is a master regulator of cell wall homeostasis through the control of a suite of autolysins ^7, 11, 12, 13^. Although the locus is highly conserved, several points of difference between bacterial genera suggest variation in the precise cellular function of the system and the associated mechanism(s) of the essentiality. WalKR is located within an operon of three to six genes that also includes a varying number of accessory factors. Two accessory genes, *yycH* and *yycI* encode membrane associated proteins that differ in their function across genera. In *S. aureus*, these proteins are activators of WalKR activity, while conversely in *Bacillus subtilis* they are repressors ^14, 15^. A second key difference between the systems is how WalKR interacts with the division septum. In *B. subtilis*, WalK controls the expression of FtsZ ^16^, it is localised to the division septum in an FtsZ dependent manner and interacts with proteins of the divisome. Consequently, WalKR essentiality in *B. subtilis* arises from the co-ordination of cell wall remodelling with cell division, in response to signalling via an extracellular Per Arnt Sim (PAS) domain ^17, 18, 19, 20^. In *S. aureus*, WalK is also reported to localise to the division septum in growing cells ^21^. Despite this, there remains no evidence of interaction with proteins of the divisome and FtsZ has not been mapped to the staphylococcal WalKR regulon. WalKR depleted *S. aureus* can be complemented with constitutively overexpressed autolysins *lytM* or *ssaA* to restore bacterial viability. However, the resultant cells have morphological defects and neither of these genes are themselves essential ^6^. The *B. subtilis* extracellular PAS domain of WalK senses peptidoglycan cleavage products generated by WalKR regulated autolysins, leading to homeostatic control of cell wall remodelling ^22^. The signal sensed by the extracellular PAS domain in *S. aureus* is not known. However, WalK activity in staphylococci (and predicted in enterococci), but not in other bacillota, is modulated through co-ordination of a divalent metal ion by an intracellular PAS domain ^23^, raising the possibility of differing roles in regulation beyond peptidoglycan biosynthesis in these genera.

The WalKR regulon in *S. aureus* has been determined by comparative transcriptomics by depleting WalKR ^7, 12^ or by the expression a constitutively active WalR phosphomemetic amino acid substitution (D53E) ^24, 25^. These studies, coupled with motif searching using a WalR DNA binding motif defined in *B. subtilis* ^11^ have built a partial map of the WalKR regulon that includes genes involved in cell wall homeostasis ^7, 12^ and virulence ^24^. Here, we applied a customised implementation of chromatin immunoprecipitation sequencing (ChIP-seq), to define a 17 bp *S. aureus* WalR consensus-binding motif and identify regulation of a number of essential genes involved in lipoteichoic acid polymerisation, ribosome biogenesis, purine nucleotide salvage/de novo synthesis and DNA replication. These data connect the regulation of cell division with chromosomal replication in *S. aureus* for the first time and identify pathways outside the regulation of autolysins to explain the essentiality of WalKR.

## Results

### Specific mutations in the *walKR* locus increase and decrease WalKR activity

We initially assembled a panel of isogenic mutants with altered WalK or WalR activity in the native context to understand regulation without under-or over-expression. These included two previously described ‘down’ mutants with decreased WalKR activity; *S. aureus* NRS384 Δ*yycHI*, with the deletion of both WalKR auxiliary proteins (*yycH* and *yycI*) ^14^, and NRS384 WalR_T101A_, in which second site PknB phosphorylation at residue T101 was abolished ^26^. We also selected two ‘up’ mutants that have increased WalKR activity; NRS384 WalK_Y32C_ with a mutation in the first transmembrane domain (identified from a sectored Δ*yycHI* colony ^23^), and NRS384 WalK_T389A_ that is predicted to prevent the dephosphorylation of WalR ^27^.

We then confirmed that each mutant exhibited the expected phenotypes with ‘up’ mutations resulting in an increased susceptibility to vancomycin and lysostaphin, increased haemolysis, reduced growth rate, and increased autolysis with the opposite true for ‘down’ mutants [Figure 1A-C] ^12, 24, 28^. Of the two ‘up’ mutants, WalK_T389A_ showed the most prominent differences, suggesting the higher level of activation [Figure 1A-C]. Mutational activation of WalKR caused a striking increase in susceptibility to oxacillin and tunicamycin, eliciting >16-fold changes in susceptibility to these cell wall targeting agents. Conversely, mutational dampening of WalKR activity caused a small but reproducible (2-fold) decrease in erythromycin susceptibility, but we did not observe erythromycin or lincomycin hypersensitivity upon WalKR modulation as has previously been reported ^5, 29^. Susceptibility to compounds targeting other cellular pathways remained unchanged [Figure 1D].

**Figure 1.**
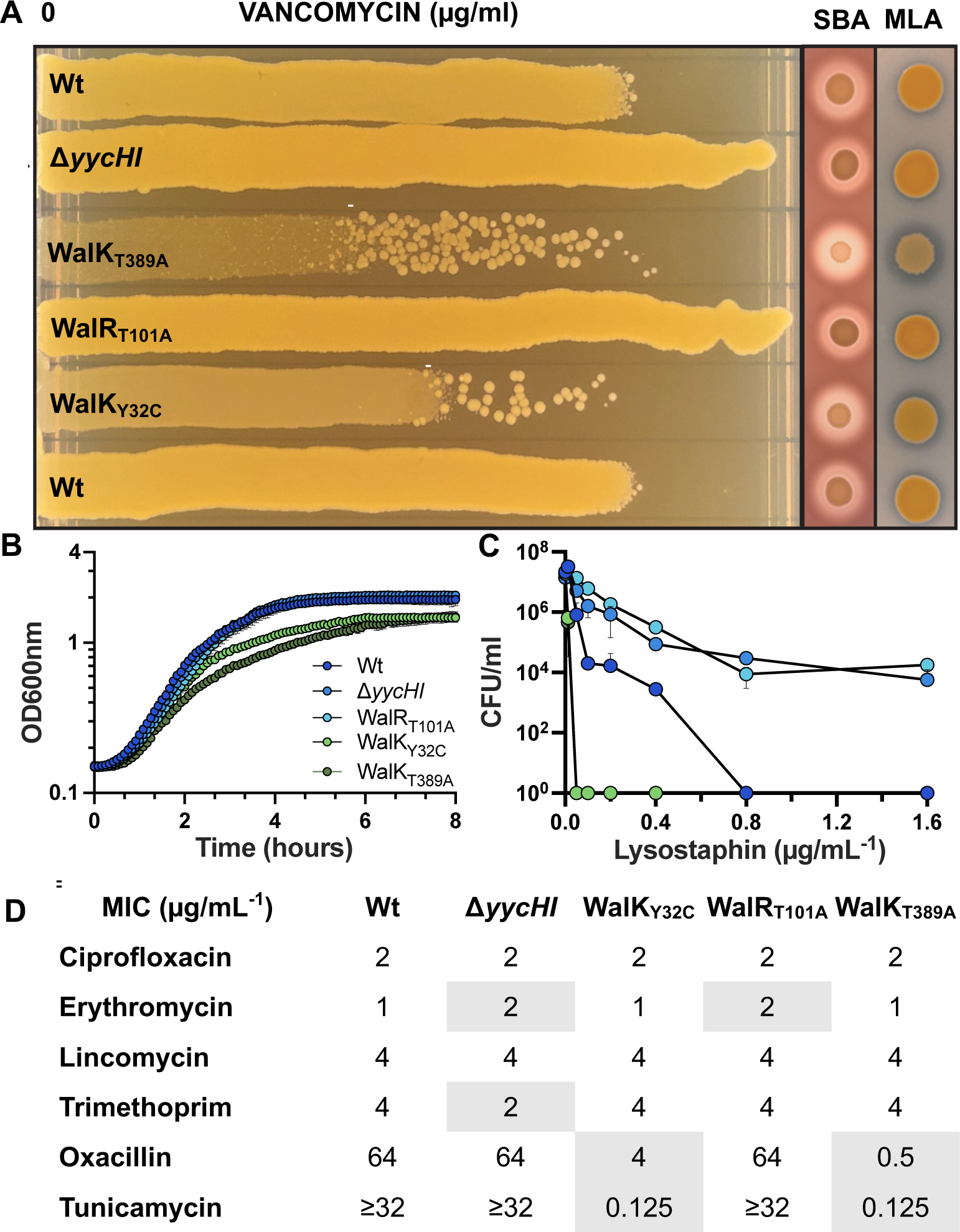
Phenotypic characterisation of WalKR mutant strains. A) Vancomycin gradient assay (VGA), sheep blood agar (SBA) Micrococcus luteus agar (MLA) for the comparison of vancomycin susceptibility, haemolysis and Atl mediated autolytic activity, respectively. White line on VGA plates highlights suppressor mutants that arise from exposure to vancomycin in the ‘up’ mutants. B) Growth curves, C) Susceptibility to lysostaphin after a 90 min exposure and D) Minimal inhibitory concentration of antibiotics against the different WalKR mutant strains, with differences to NRS384 Wt highlighted.

TCS phosphoryl transfer from the sensor histidine kinase to the cognate DNA-binding response regulator requires direct interaction ^30^. To assess the impact of the ‘up’ and ‘down’ mutations on interaction dynamics between WalK and WalR across growth, we implemented a split luciferase system ^31^. Proteins were C-terminally tagged (separated by a glycine serine linker) with either the small bit (SmBIT-11 amino acids) or large bit (LgBIT – 17.6 kDa) to reconstitute a functional luciferase that emits light in the presence of the furimazine substrate. Our modifications allowed the kinetics of protein-protein interaction to be non-invasively measured throughout *S. aureus* growth. Chromosomal C-terminal tagging of WalR-SmBIT and WalK-LgBIT at the native locus showed that the proteins tolerated the presence of either tag, with no growth defect detected [Figure S1].

We then constructed WalR-SmBIT and WalK-LgBIT fusions in the plasmid system with native and mutant proteins, following cell density (OD600nm) and light emission (RLU) throughout *S. aureus* growth. We observed immediate interaction of WalK with WalR upon dilution in fresh LB with the peak interaction in the mid-exponential phase of growth, and subsequent rapid decline of the interaction to undetectable levels (5 h) in stationary phase. This pattern of interaction was enhanced in the ‘up’ mutant strains throughout growth (including lag phase) for both WalK_Y32C_, WalK_T389A_, and the previously described WalR_D53E_ mutant ^24, 25^ compared to the native WalK/WalR interaction [Figure 2]. In contrast, the ‘down’ mutants strains, which included WalR_D53A_ (cannot be phosphorylated by WalK), WalR_T101A_, WalK_D53A/T101A_, and WalK_G223D_ (reduced autophosphorylation and transfer to WalR ^32^), exhibited a consistent profile of reduced initial WalK/WalR interaction during lag phase in comparison to the native WalK/WalR alleles [Figure 2]. The kinetic profile of the WalR_T101A_ mutant mimicked the interaction profile of the WalK_G223D_ mutant with a 1.5-fold increase in the maximal level of interaction (compared to native WalK/WalR). This increase correlated with the same mid-log phase time point as seen in the native interaction [Figure 2]. ‘Down’ mutants also yielded detectable interaction into stationary phase. These interaction profiles combined with phenotypic profiling validate our panel of ‘up’ and ‘down’ mutants.

**Figure 2.**
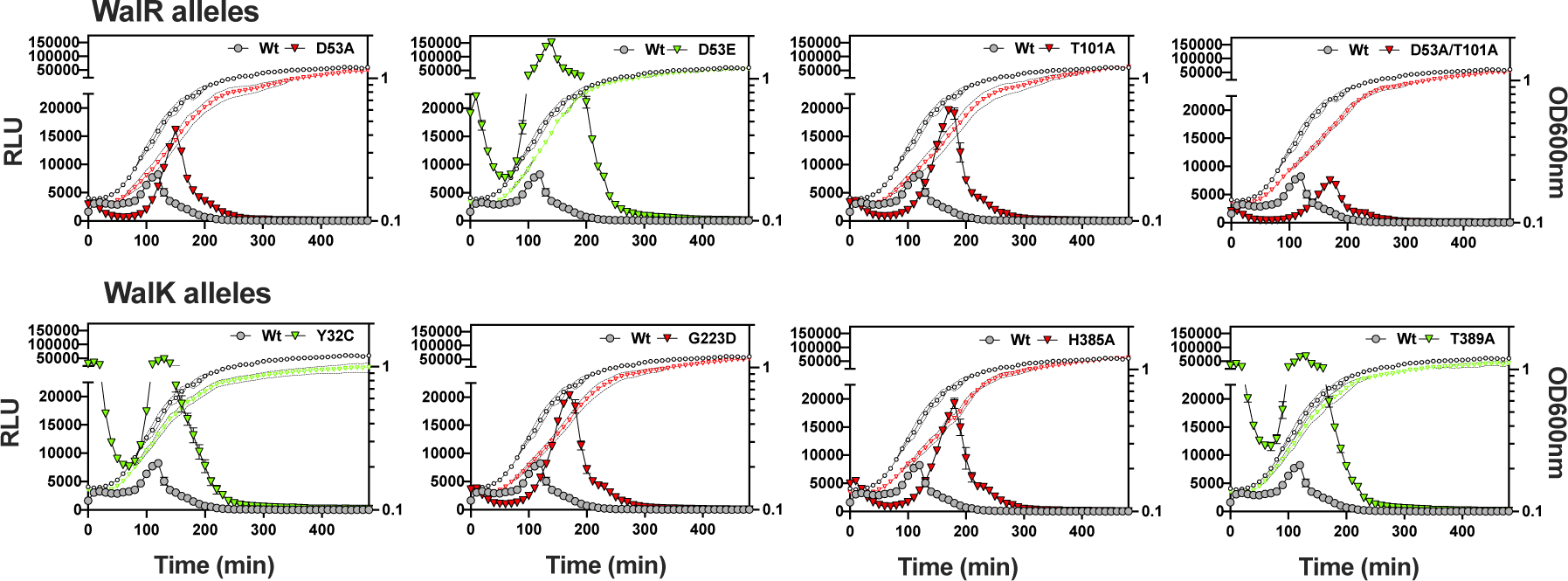
Kinetic interaction of WalR with WalK in S. aureus. Analysis of the interaction of WalR-SmBIT with WalK-LgBIT and WalK/WalR mutant proteins throughout growth in NRS384 Wt. OD600nm: open black circle (WalR/WalK Wt) and open red triangle (‘down’ mutant) or open green triangle (‘up’ mutant). RLU: grey filled circle (WalR/WalK Wt) and either red (‘down’ mutation) or green (‘up’ mutation) filled triangle. Results are the mean from three independent determinations and the error bars show the standard deviation.

### WalKR activation causes a global change in gene expression

To ascertain the impact of WalKR activation on *S. aureus* gene expression we compared the transcriptome of ‘up’ mutant WalK_T389A_ to the NRS384 strain. Activation of WalKR was associated with a global gene expression change, wherein ∼55% of genes (1,117) were significant (False Discovery Rate (FDR) ≤0.05, log_2_FC ≥0.585) with approximately half (551) up-regulated and half (564) down-regulated [Figure 3C]. Genes with increased expression (≥1.5 log_2_FC, FDR ≤0.05) included those encoding autolysins, virulence factors, membrane transporters, and proteins of the amino acid, purine, and fatty acid biosynthesis pathways [Figure 3A, Table S1]. Genes with decreased expression (log_2_FC ≤-1.5, FDR ≤0.05) were primarily involved in oxidative stress response, metal ion homeostasis, and protein fate [Figure 3A, Table S1]. Whereas the expression of genes encoding ribosomal proteins, transcription factors, and proteins involved in carbohydrate metabolism, DNA maintenance, and cell wall organisation were both up- and down-regulated (FDR ≤0.05, log_2_FC ≥1.5 or ≤-1.5) [Figure 3A, Table S1].

**Figure 3.**
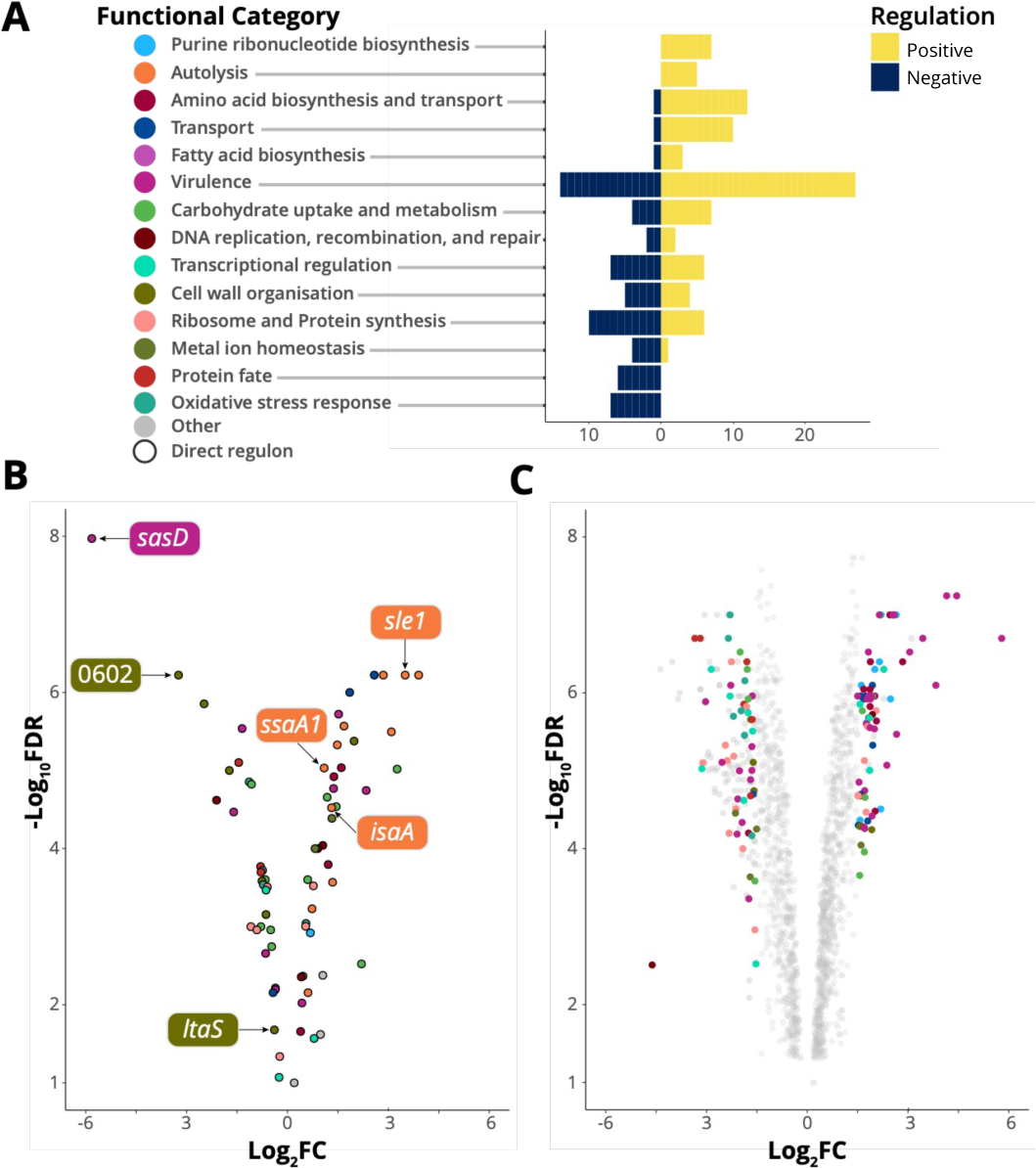
The direct and indirect WalKR regulon. A) The number of genes in each category that is positively or negatively regulated upon WalK activation (FDR ≤0.05, log_2_FC ≥1.5 or ≤-1.5). For clarity, categories where n ≤ 2 are not shown B) Gene expression changes of the predicted direct WalKR regulon upon WalK activation (Wt vs WalK_T389A_). Genes with WalR binding sites confirmed by ChIP-seq are highlighted (see Figure 4D and Table S1). C) Global change in transcriptome upon mutational activation of WalK (Wt vs WalKT389A), members of gene sets undergoing a log_2_FC ≥1.5 or ≤-1.5 are highlighted. Y-axis labels in panel A serve as the legend for panels B and C.

Explanations for two of our observed WalKR phenotypes can be inferred in the WalK_T389A_ RNA-seq data. Firstly, increased haemoylsis [Figure 1A] in WalK_T389A_ mutants is explained by an increase in *hla* expression (+1.7-log_2_FC), encoding alpha-toxin, upon WalK activation. Secondly, enhanced zones of clearing surrounding WalK_T389A_ on heat killed *Micrococcus luteus* agar plates [Figure 1A] arise due to increased peptidoglycan degradation of secreted processed Atl by the dual activity autolysin Atl ^33^ (+2.85 −log_2_ FC).

### Defining an *in vivo* WalR regulon using ChIP-seq

The direct regulon of WalR was then investigated using ChIP-seq. To permit immunoprecipitation, a 1xFLAG-tag was incorporated onto the C-terminus of WalR using a modified anhydrotetracycline (aTc) inducible expression plasmid ^34^. To validate functionally, the transcriptome of a strain *S. aureus* NRS384 with chromosomally C-terminally FLAG tagged WalR ^23^ was compared to the wild type (Wt) strain. This strain had a gene expression profile that was like NRS384 Wt during mid-log phase, as determined by RNA-seq (no significant changes in gene expression (0.585 Log_2_FC, 0.05 false discovery rate [FDR]). Subsequently, FLAG-tagged expression constructs were also made in pRAB11 for three other response regulators with known DNA-binding motifs, HptR, and SaeR, and VraR [Figure 4A] ^35, 36, 37^. The four plasmids and an empty vector control were each transformed into the CA-MRSA USA300 strain NRS384 with *spa*, which encodes Protein A, deleted to reduce non-specific IgG-binding. Dose-dependent aTc induction was observed (induced at an optical density of 600 nm (OD600nm) = 0.6 for 1 h) for *walR* [Figure 4B]. We examined the impact of increasing aTc concentrations on tag-counts (mapped sequence reads for ChIP purified DNA) and selected an aTc concentration of 100 ng•mL^-1^ for subsequent ChIP-seq experiments [Figure 4C]. ChIP-seq was then conducted with each of the four constructs and the empty vector. Initial analysis of the resulting sequence reads revealed a high background of reads mapping across the entire *S. aureus* chromosome. To improve the signal-to-noise ratio for each of the four ChIP-seq experiments, we performed *in silico* subtraction of the read sets for the three non-target response-regulators and the empty vector from the read set for the target response-regulator [Figure S2, Table S2]. This revealed peaks anticipated for HptR and SaeR, upstream of *hpt* and *hla*, respectively [Figure 4D, Table S1], and identified WalR binding sites upstream of six genes that included autolysins SAUSA300_2249 (*ssaA*), SAUSA300_2506 (*isaA*), SAUSA300_0438 (*sle1*), cell wall cross-linked SAUSA300_0136 (*sasD*), hypothetical secreted SAUSA300_0602, and SAUSA300_0703 (*ltaS*) which encodes the essential lipoteichoic acid (LTA) synthase responsible for polymerising glycerol-6-phosphate into LTA chains ^38^ [Figure 4D, Table S1]. All these genes had previously been identified as belonging to the WalKR regulon ^7, 24^. For VraR, we observed a ChIP-seq binding site upstream of *ltaS*, that was adjacent to the putative WalR-binding site [Figure 4D].

**Figure 4.**
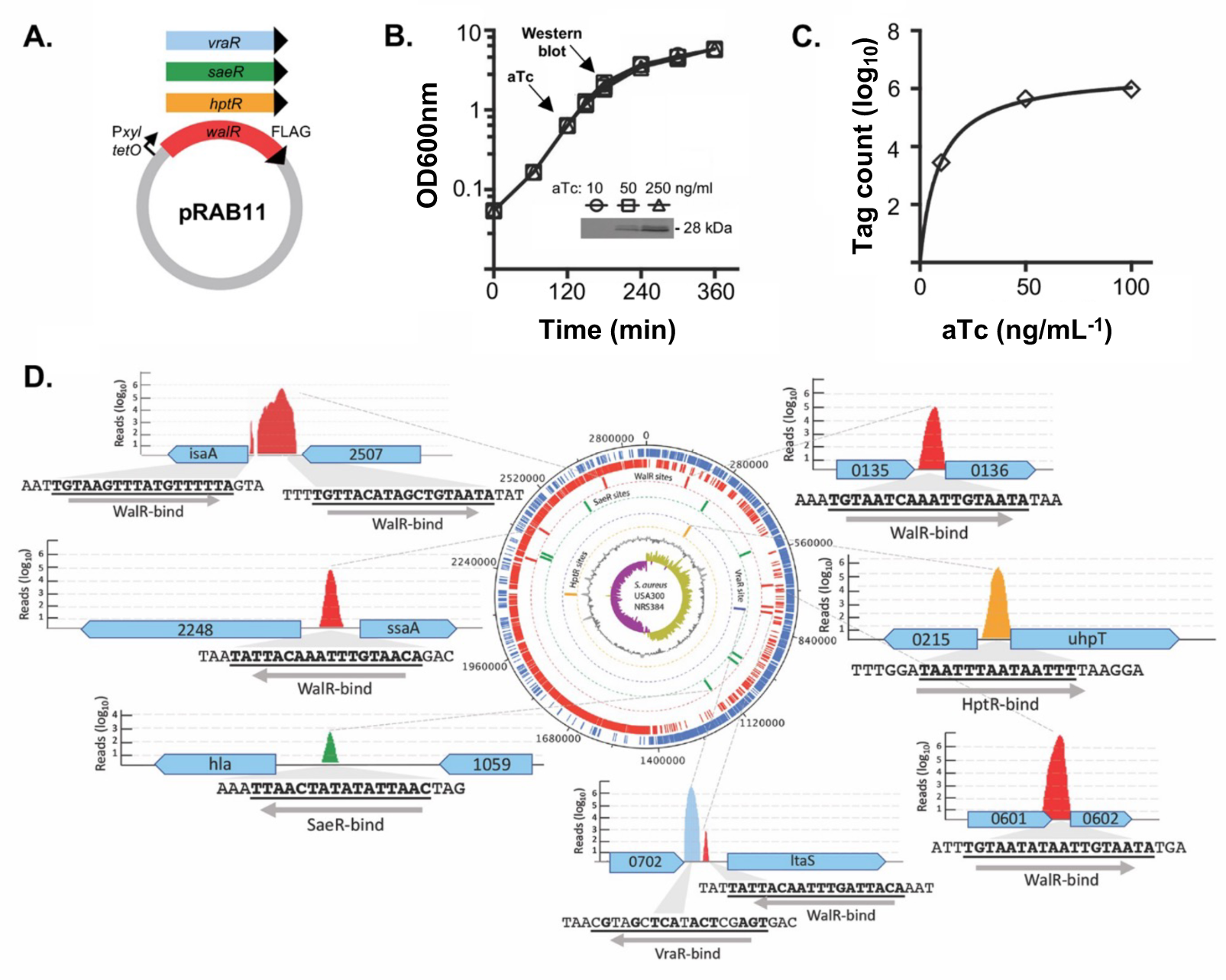
ChIP-seq analysis of S. aureus WalR chromosome binding sites. A) Four response regulators with a Cterminal FLAG-tag for immunoprecipitation after induction with anhydrotetracycline (aTc) in NRS384Δspa. B) Dose dependent production of WalR-FLAG (inset Western blot) with increasing concentrations of aTc. C) Plot showing an increasing number of WalR ChIP-seq mapped reads (Tag count) as the aTc concentration increases. D) Summary of key ChIP-seq binding sites across the S. aureus chromosome, showing binding motif and direction, including five WalR peaks (red), and single peaks for SaeR (green), VraR (blue) and HptR (yellow).

Including the above six genes (seven peaks), a total of 22 WalR ChIP-seq peaks upstream of 21 genes were identified, with 12 set aside due to small peak size (7-20 nucleotides) and poor peak prediction scores. Two peaks were identified in 16S rRNA genes, but the use of ribosomal RNA depletion precluded interpretation of RNA-seq data for these loci. The peak upstream of SAUSA300_0681 was not pursued, as no candidate WalR binding sites were identified (Table S1].

The high stringency required for the *in-silico* subtraction approach could eliminate peaks corresponding to lower affinity DNA binding regions and thereby obscure the identification across the breadth of the direct WalR regulon. Consequently, we used ChIP-seq defined WalR binding regions in conjunction with previously validated WalR binding sites ^12,24^ to generate a 17 bp consensus *S. aureus* WalR-binding motif (5’-TGTHH[N]_6_WGTNDD-3’) [Figure S3 AB]. This motif was the used to conduct an *in-silico* search of the genome for potential WalR binding sites. In total, 118 putative intergenic WalR binding sites were identified within the NRS384 chromosome [Figure S2B], of which 109 were within 500 bp of a predicted CDS transcriptional start site [Table S3].

### Positioning of WalR binding site, not sequence or orientation, dictates of mode of regulation

To investigate whether the sequence of WalR motifs could determine the mode, or degree, of change in gene expression upon activation, WalR motif diversity was visualised as a maximum-likelihood phylogenetic tree and the tips labelled with gene expression changes upon WalK activation [Figure S4]. Functional groups of regulated genes were also mapped to examine whether specific motif signatures were linked to gene sets [Figure S5]. We did not observe clustering of positively or negatively regulated genes, nor did branches of the tree correspond to specific gene functions [Figure S4]. Taken together, these analyses indicate that the sequence of WalR motifs does not dictate the mode of regulation, nor is it linked to the functional class of the gene it controls.

Here, building on framework used to analyse WalR in *B. subtilis* ^13^, we mapped the positions of *S. aureus* WalR binding sites of the direct regulon in relation to predicted transcriptional start sites and promoters ^39^ [Table S4]. The orientation of WalR binding site in relation to the downstream gene was not significantly associated with the magnitude (Student’s t test, p=0.52) or mode (p=0.81) of expression change. However, the position of the WalR binding site did dictate the mode of regulation; WalR binding upstream of the −35 element was significantly associated with positive regulation (p < 0.001), whereas WalR binding between, or downstream of, −35 and −10 elements was associated with negative regulation (p < 0.001) [Table S4]. Thus, whether WalR activates or represses the expression of a gene is determined primarily by its position relative to the promoter rather than motif sequence.

### Functional validation of WalR directly regulated genes

WalR binding to promoters of the six loci identified by ChIP-seq was assessed *in vitro* using electrophoretic mobility shift assays (EMSAs) with recombinant WalR [Figure 5A, Figure S5]. We observed WalR binding to all six promoters identified by ChIP-seq, with specificity confirmed through competition experiments with excess labelled or unlabelled DNA duplexes [Figure S6]. We also corroborated VraR binding to the *ltaS* promoter at the consensus VraR binding motif (5ʹ-TGA[N_1-3_]TCA-3ʹ)^35, 40^ [Figure S7A], while WalR had no affinity for this duplex [Figure S7B]. VraR did exhibit affinity for the WalR binding site duplex, however, this was shown to be non-specific by competition assay [Figure S7C]. Therefore, WalR and VraR bound to the *ltaS* promoter at discrete sites dictated by their respective recognition motifs.

**Figure 5.**
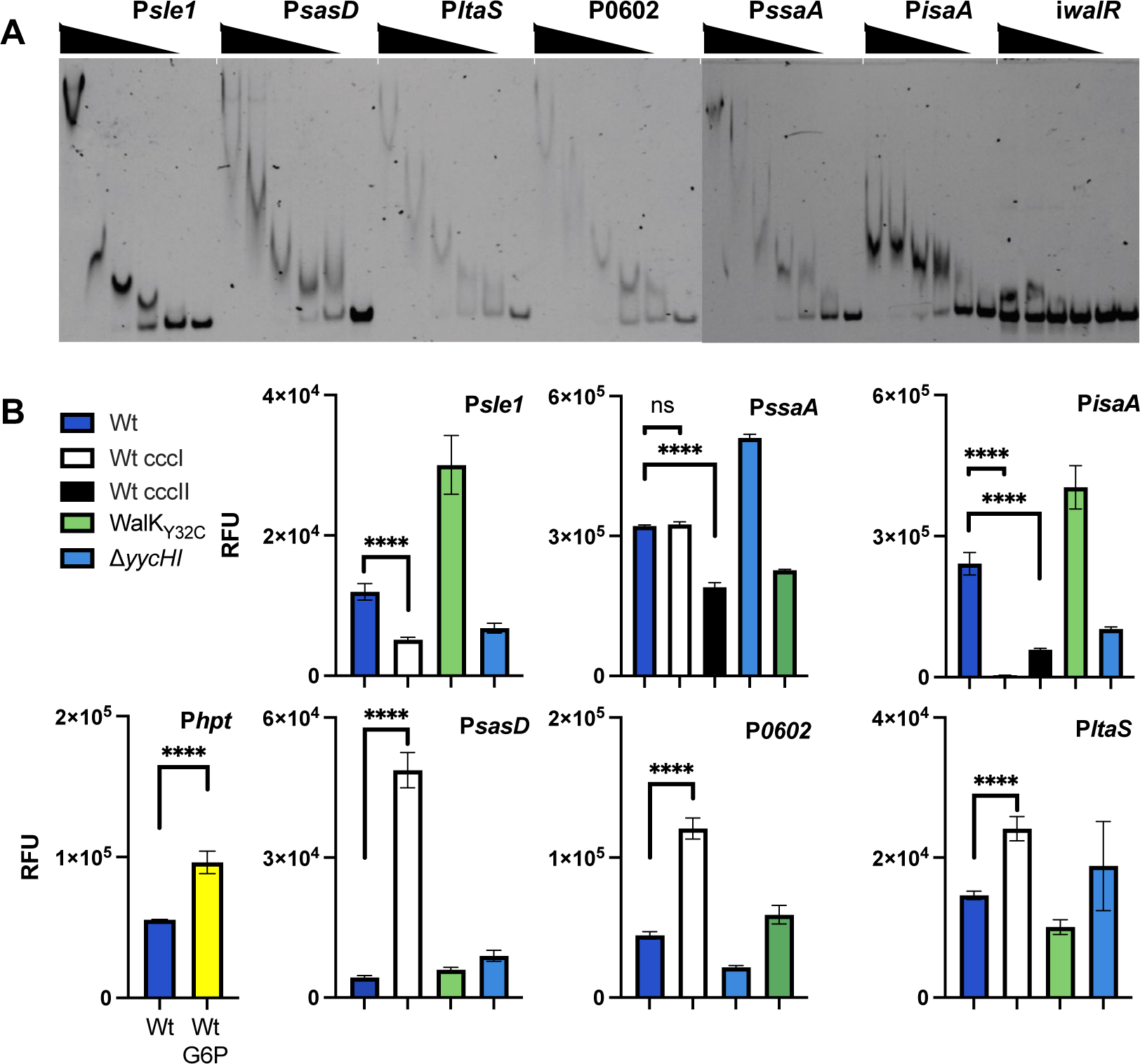
In vitro and in cellulo confirmation of WalR regulation. A) Dose dependent band shifts caused through specific WalR binding to the ChIP-seq identified promoter regions. An intragenic region of the walR gene was used as a non-specific control (iwalR). A total of 5 doubling dilutions (plus a no protein control – right lane) of WalR starting at 20 µM, for all except PssaA where 40 µM was used. B) Cumulative YFP fluorescence of S. aureus reporter strains from stationary phase cultures. The glucose-6-phosphate (G6P) inducible hpt gene was used as a positive assay control through addition of 25 µM of G6P. Promoter activity was assessed in NRS384, with or without an abrogating (ccc) mutation in the WalR binding site. The native promoter construct was also analysed in WalKY32C (‘up’ mutant) or ΔyycHI (‘down’ mutant) strains. Results are means from three independent experiments with the error bars showing the standard deviation of the mean. The significance of the ccc mutation was determined using an unpaired Student’s t test; ns: not significant, <0.0001: **** versus the Wt promoter in NRS384.

Following the *in vitro* confirmation of WalR binding to sites identified by ChIP-seq, we sought to assess the impact of WalR binding on promoter activity in *S. aureus* [Figure 5B]. The promoter regions encompassing the WalR binding motif were transcriptionally fused with yellow fluorescent protein (YFP). To assess the impact of WalR activity on gene expression, each construct and a paired WalR binding motif mutant (first TGT in WalR motif mutated to CCC) was transformed into NRS384. Additionally, the native promoter construct was transformed into NRS384 ‘down’ mutant (Δ*yycHI*) and ‘up’ mutant (WalK_Y32C_). The WalK_Y32C_ strain was chosen rather than WalK_T389A_ as suppressor mutants arose in this background through genetic instability, which was not observed in WalK_Y32C_. As a positive control, we included the promoter for the *hpt gene* encoding the glucose-6-phosphate transporter which is responsive to the presence of glucose-6-phosphate ^37^ [Figure 5B]. Strains were grown to stationary phase with fluorescence and colony forming units determined. All of the RNA-seq down regulated genes (*sasD*, *ltaS,* and SAUSA300_0602), exhibited increased fluorescence upon abrogation of the WalR binding motif, showed increased fluorescence in the ‘down’ mutant (Δ*yycHI*), and decreased fluorescence in the ‘up’ mutant (WalK_Y32C_), indicative of negative regulation by WalR [Figure 5B]. For upregulated genes, abrogation of the WalR binding motif caused a reduction in fluorescence, with all showing increased activity in the WalK_Y32C_ background and decreased activity in the Δ*yycHI* mutant, characteristic of positive WalR regulation [Figure 5B]. As the *isaA* and *ssaA* promoter regions contain two WalR binding motifs, both were individually mutated. For *isaA*, the WalR binding site closest to the TSS (CCCI, 37 bp to TSS) caused the greatest decrease in promoter activity ^28^, whereas for *ssaA*, only the more distal binding site (CCCII, 167 bp from the TSS) reduced expression, which corresponded to the single ChIP-seq peak for *ssaA* [Figure 4D].

### Control over additional essential genes by WalKR

To investigate WalKR essentiality in *S. aureus* and triage genes for further analysis based on their likely contribution to the essentiality phenotype, we analysed intersecting data sets where genes fulfilled the following criteria: (i) contain a predicted upstream WalR-binding site; (ii) belong to the core *S. aureus* genome; (iii) essential for growth in rich media ^41, 42, 43^; and (iv) their expression is significantly changed upon WalK activation as defined by RNA-seq [Figure 3A]. We found that within the predicted direct WalR regulon, seven essential genes undergo a significant change in gene expression upon activation by WalKR (FDR ≤ 0.05, log_2_ FC ≥0.585) [Figure 3AB]. These genes were *ltaS* (see above); *dnaA*, which encodes chromosomal replication initiator protein; *hup,* the sole DNA-binding protein HU; *prs*, which encodes a ribose-phosphate pyrophosphokinase involved in purine salvage and de novo synthesis; *rplK,* ribosomal protein L11 (50S subunit component); and *tagG* and *tarF*, which encode teichoic acid biosynthetic proteins [Figure 6AB]. Additionally, three essential genes had a predicted upstream WalR binding site but did not undergo a significant change in expression: *dnaD*, *rnz*, and *sufB*, encoding putative replication restart protein DnaD, ribonuclease Z, and FeS assembly protein SufB. Of these three essential genes, a ChIP-seq peak was identified upstream of *dnaD* [Figure 6A].

**Figure 6.**
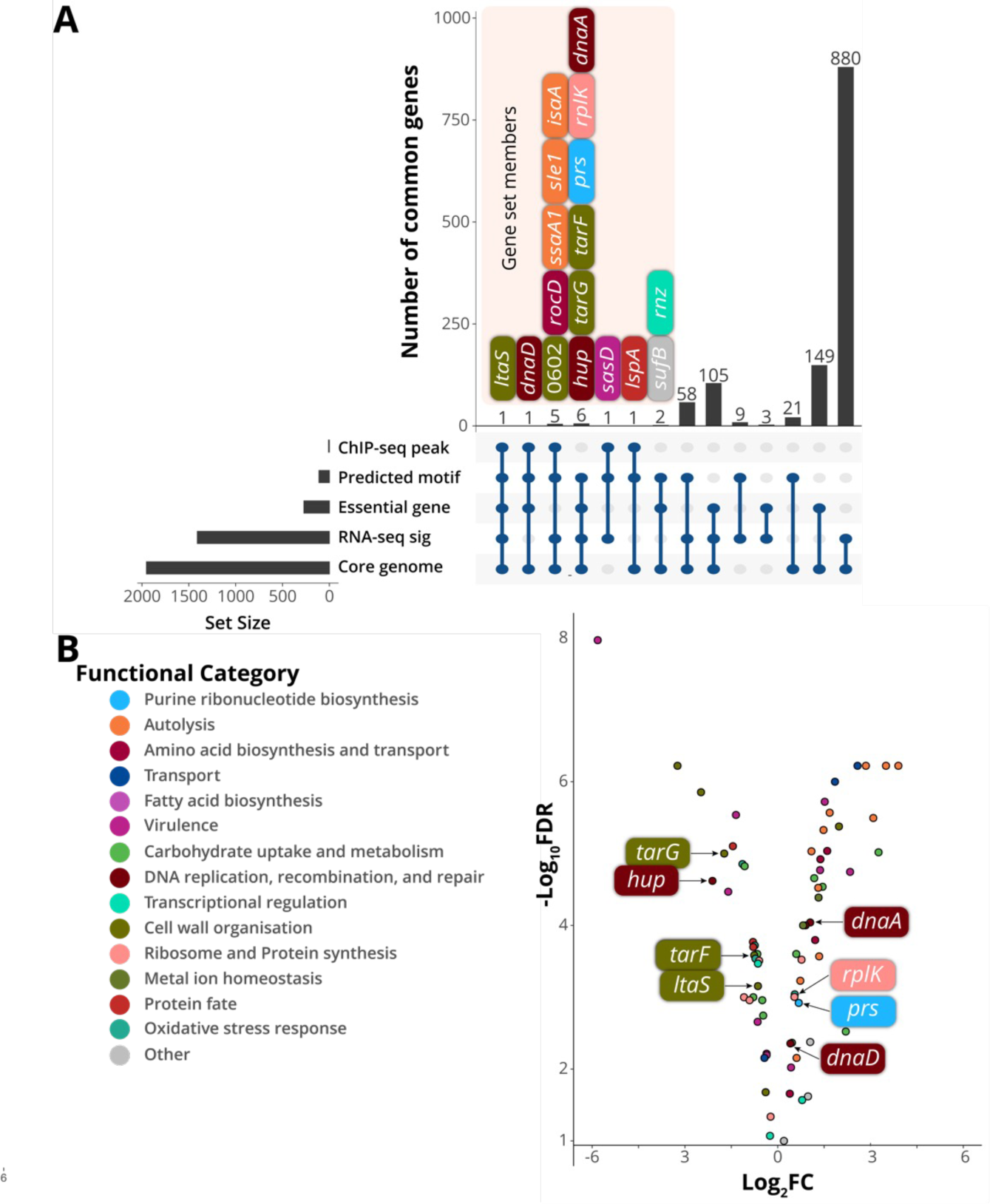
Analysis of putative WalR controlled essential genes. A) Intersections between essentiality analysis and -omics data displayed as an UpSet plot. Members of gene sets where n<8 are shown. RNA-seq significance ((RNA-seq sig) FDR £0.05, log2FC ≥0.585). B) Gene expression changes of the predicted direct WalKR regulon upon WalK activation (Wt vs WalKT389A). Essential genes with predicted WalR binding sites are highlighted.

To further characterise the seven essential genes within the predicted direct WalR regulon and *dnaD* ^44, 45^ [Table S3], we built a time-resolved picture of their expression. The promoter regions for these genes, with or without an abrogating mutation (TGT to CCC) in the WalR-binding site, were introduced into a luciferase reporter plasmid and transformed into *S. aureus* NRS384 [Figure 7A-C]. The resulting *S. aureus* strains were monitored for growth and light emission every 10 min over an 8 h period. For comparison, promoters from the previously characterised WalR regulated genes and the *walR* gene itself were included [Figure 7AB]. Expression from the *walR* promoter rapidly peaked in mid-exponential phase and then tapered off, as has been observed previously [Figure 7B] ^46^. The previously ascribed WalR regulation of the six ChIP-seq hits were corroborated by the luciferase reporter assays; mutation of the WalR binding site yielded a significant reduction in the level of expression throughout growth for the positively regulated genes (*sle1, ssaA* and *isaA*)[Figure 7A]. While an increased level of expression was observed in the negatively regulated genes upon mutation of the WalR binding site mutation (*sasD*, SAUSA300_0602 and *ltaS*)[Figure 7B]. Loss of the WalR negative regulation was shown to relieve repression of *sasD,* while the impact on SAUSA300_0602 was most pronounced into stationary phase of growth. Only a subtle difference in expression was observed for *ltaS*; mutational abrogation of WalR binding prevented “turning off” of gene expression, resulting in prolonged expression into stationary phase [Figure 7B]. We then examined the impact on the additional essential genes identified from the analysis of the direct regulon. No regulation was detected for *tagG, tarF* or *dnaD* [Figure S8]. However, the very low level of *dnaD* and *tarF* promoter activity under the conditions tested precluded a definitive determination. Whereas a strong positive regulation was observed for *rplK. prs* was also shown to be positively regulated*. hup* was shown to be weakly negatively regulated by WalR, which caused a reduction in the expression upon the transition into stationary phase, similar to *ltaS*. For hup there was a bi-phasic change at 190 min present in the Wt strain, which was lost when the binding site was mutated [Figure 7B]. Closer inspection of the large (730 bp) upstream intergenic region between the divergent *dnaA* and *rpmH* genes revealed an enrichment of potential WalR binding motifs upstream of *dnaA* [Figure 6B]. The three sites proximal to the *dnaA* gene were chosen for further analysis (site X was not investigated, [Figure 7C]). Sites ccc II and ccc III impacted the expression of *dnaA* while no change in expression was observed for the mutation of ccc I [Figure 7C]. We observed both a negative regulation for ccc II and a complete abrogation of expression for ccc III, suggesting a complex fine tuning of *dnaA* expression afforded by WalR. Recently, a suppressor mutant with a deletion in the *dnaA* promoter that reduced the level of DnaA activity (initiation of DNA replication) was identified in a *S. aureus Δnoc* strain ^47^. This deletion removed a 56 bp region surrounding the WalR binding site denoted by ccc II [Figure 7C]. Here, in agreement with this previous observation, we identified the reduced expression from the *dnaA* promoter upon abrogation of the ccc II WalR binding site.

**Figure 7.**
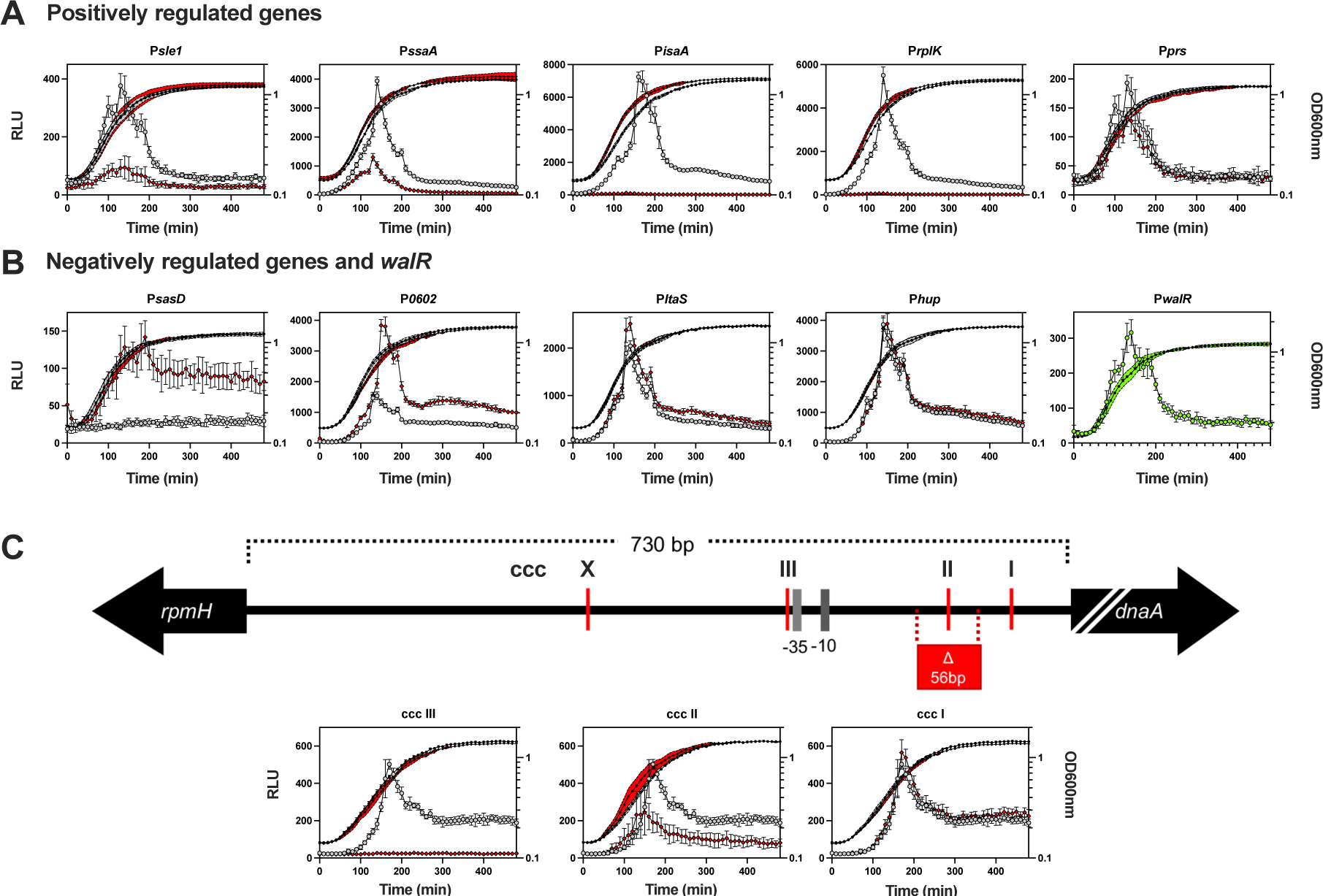
Characterising the temporal control of WalR on essential S. aureus genes. A) Positively and B) negatively WalR regulated genes coupled to bacterial luciferase reporters showing the changes in promoter activity of the Wt (open grey circle: RLU; filled black circles: OD600nm) or the ccc (open red diamond: RLU; filled black diamond: OD600nm) mutated WalR binding site in LB media over time. Data represent the mean of three independent experiments (± standard deviation). Expression of walR is represented by green open circles (RLU), with growth indicated by filled black circles (OD600nm). C) Schematic (to scale, except dnaA –truncation indicated by white back slashes) of the dnaA promoter region with identified WalR sites denoted. The Wt and the three dnaA motif mutants (ccc I-III) were analysed as described above. Highlighted in red is the 56 bp deletion^47^ and the −35 and −10 sites for the dnaA promoter.

### Modulation of WalKR activity alters *S. aureus* lipoteichoic acid structure

Lipoteichoic acid (LTA) is an anionic polymer composed of a repeating chain of glycerol phosphate units anchored to the cell membrane via a diacylglycerol group and decorated with side chain modifications of D-alanine and/or glycosyl moieties ^48^. Here, the physiological impact of negative *ltaS* regulation by WalR was assessed by extracting LTA from strains of representing a spectrum of WalKR activities. Polyacrylamide gel electrophoresis (PAGE) of LTA enables visualisation via laddered banding patterns, with each subsequent band representing a polymer chain length of n + 1 ^49^. Side chain modifications can also be observed, with high levels of heterogenous modification resulting in loss of single band resolution, i.e., smearing ^49^. In the Wt or a ‘down’ mutant yielded a characteristic smeared banding pattern, indicative of heterogenous side-chain modification, whereas in a WalKR ‘up’ mutant (WalK_T389A_) LTA chain length changes were observed. This manifested as a reduction in mid-length LTA [Figure 8, orange line] and increase in high molecular weight LTA [Figure 8, black line]. Analysis of the supernatant of the ‘up’ strain resulted in LTA shedding, consistent with the compromised cell wall [Figure 1C].

**Figure 8.**
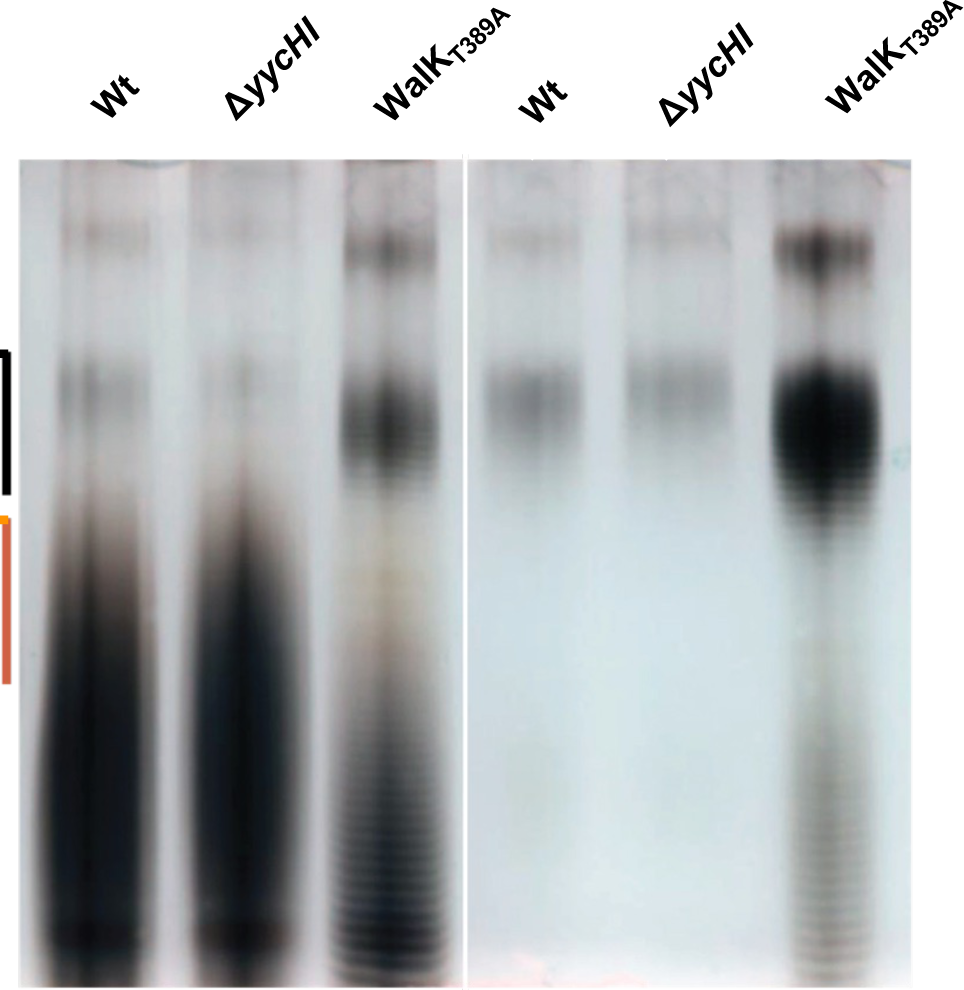
Assessing the influence of WalKR activity on LTA production. PAGE analysis of purified LTA from NRS384 and isogenic mutant derivative strains grown in LB. Each band indicates a glycerol phosphate polymer length of n+1. Where bands are not clearly visible, this is due to heterogenous side chain modifications resulting in smearing patterns. Black and orange lines are shown and represent regions of distinct difference in LTA banding patterns between the strains.

### WalR controls DNA compaction by regulation of the DNA binding protein HU

The essential gene *hup* ^50^, encodes the sole *S. aureus* DNA-binding protein HU, was indicated to be negatively regulated by WalR [Figure 7B]. *S. aureus* HU belongs to a family of low molecular weight nucleoid-associated proteins (NAPs) and although it is largely uncharacterised, its structure has been determined ^51^. Orthologs of HU NAPs from other bacteria have previously been shown to control DNA compaction, introducing negative supercoils into relaxed DNA ^52, 53^, and play an essential role in the initiation of DNA replication in *B. subtilis* ^54^. To investigate potential changes in DNA topology mediated by *S. aureus* HU and WalR regulation of HU, we performed CRISPRi knockdown of *walR, hup,* and as a negative control, *hla*. The CRISPRi knockdown titrated the expression of each targeted gene [Figure 9A], resulting in 57-, 20-, and 78-fold downregulation of *hup*, *walR*, and *hla*, respectively. Notably, a degree of knockdown in the absence of inducer was observed, attributable to the leaky expression of the CRISPRi guide and the location of the guide (overlapping the promoter for *hup*). HU knockdown resulted in a relaxation of plasmid supercoiling, shown by increased DNA band intensity during chloroquine gel electrophoresis, due to more slowly migrating topoisomers in comparison to the control [Figure 9A pSD1 vs pSD1*hup*, white up arrow]. Knockdown of *walR* had the opposite effect, indicated by increased DNA band intensities arising from faster migrating topoisomers with greater supercoiling density [Figure 9A pSD1 vs pSD1*walR*, white down arrow]. This observation is consistent with a model where WalR negatively regulates HU and is further supported by the knockdown of *hla* or the empty vector having no impact on DNA topology [Figure 9A pSD1 vs pSD1*hla*]. These results show that *S. aureus* HU, as observed in other bacteria, increases the supercoiling density of DNA. We propose that the negative regulation of *hup* by WalR has the opposite effect, causing the relaxation of supercoiling and leading to decreased compaction of cellular DNA.

**Figure 9.**
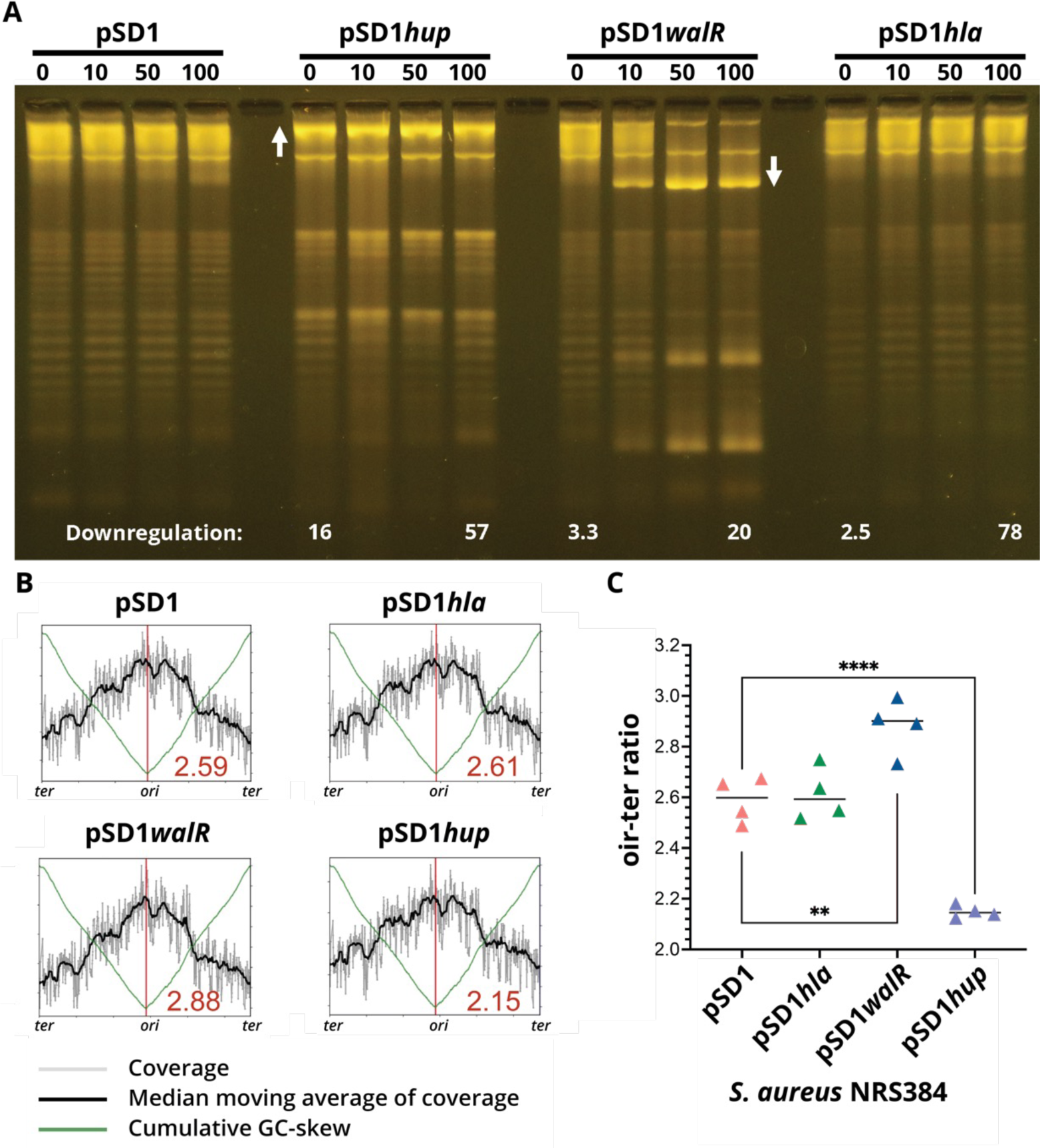
Impact of CRISPRi mediated downregulation of hup or walR on DNA topology. A) NRS384 containing either pSD1 (empty vector), pSD1hup, pSD1walR or the negative control pSD1hla were induced with increasing concentrations of aTc (0, 10, 50 and 100 ng/mL). At 5 h post induction total RNA and plasmids were isolated. Plasmids were run on a 1% (w/v) TBE gel containing 2.5 µg/mL chloroquine to allow for the resolution and visualisation of discrete topoisomers of supercoiled plasmid DNA. Relative fold downregulation of each targeted gene compared to uninduced vector control (pSD1, 0) as determined by RT-qPCR is denoted at the bottom of the image. The image is representative of three repeat experiments. B) A representative whole genome sequencing read-coverage graph showing the ori-ter ratios for the four CRISPRi constructs under 100 ng/mL aTc induction. Each plot represents one of four biological sequencing replicates for each CRISPRi construct. Shown in the bottom right quadrant of each graph is the mean ori-ter ratio across the four replicates. The ori-ter ratio between the sequence read coverage at the origin and terminus. C) Summary graph of ori-ter ratios for each of the four CRISPRi constructs induced with 100 ng/mL of aTc to downregulate expression of the three targeted genes (hla, walR and hup). Shown are the mean of quadruplicate, independent biological replicate sequencing experiments. Differences between means assessed using an unpaired student’s t test, **p=0.0059, ***p=<0.0001.

### Changes in initiation of DNA replication mediated by WalR and HU

We next assessed whether WalKR modulation of *hup* could also impact the initiation of DNA replication in *S. aureus*, building on the observation that this was impaired in *B. subtilis* by depletion of orthologous protein ^55^. This was addressed by genome sequencing of the CRISPRi knockdown constructs of *S. aureus* for *walR, hup, hla* and vector control, and aTc induction (0 or 100 ng/mL^-1^) and determining the number of chromosome replication origins per cell during exponential growth ^47, 54^. Mean *ori*-to-*ter* ratios were calculated and revealed a significant reduction in the *ori*-to-*ter* ratio from ∼2.60 in the vector control and non-target *hla* control compared to 2.15 when the expression of *hup* was repressed [Figure 9BC]. A significant increase (2.88) in the mean *ori*-to-*ter* ratio, was observed when *walR* was repressed [Figure 9BC], consistent with the negative regulation of *hup* by WalR [Figure 7B]. Taken together, these data show that *S. aureus* HU contributes to promoting initiation of DNA replication and is directly influenced by WalKR-mediated regulation.

Collectively, our findings above link diverse and crucial cellular processes, including DNA replication and cell wall homeostasis, to the activity of WalKR. Distinct from other bacilliota, WalKR appears to serve as a crucial nexus for both regulatory and temporal coordination of these diverse activities. Thus, this work provides a mechanistic basis for the essentiality of WalKR in *S. aureus*.

## Discussion

Here we use ChIP-seq to define genes directly under the control of WalKR. Subsequently, we generated a *S. aureus* specific WalR consensus binding motif and to inform physiologically relevant cut-offs for interrogating transcriptomic data obtained when mutationally activating WalK. These experiments allowed us to define a direct and indirect WalKR regulon. These results confirm and expand the pioneering discoveries of Dubrac and Msadek *et al*., with WalKR directly regulating several autolysins with activation increasing the production SaeRS regulated virulence genes ^7, 12, 24^. Additionally, consistent with our earlier work ^56^, we observed upregulation of a variety of genes involved in central metabolism upon WalK activation, particularly those involved in amino acid, purine, and fatty acid biosynthesis. Filtering our results on essential genes directly controlled by WalKR, we identified and defined new members of the WalKR direct regulon; *rplK*, *hup, ltaS, prs* and *dnaA*. Of which, *prs* and *rplK*, belong to the central metabolic pathways of purine and protein biosynthesis, respectively.

We find that WalKR negatively regulates the expression of *ltaS,* dampening expression from late exponential into stationary phase. To our knowledge, this is the first report of direct transcriptional regulation of *ltaS*, although post-translational regulation has been described ^57^. We also detected VraR binding to a second site further upstream in the *ltaS* promoter. Though comparative transcriptomics *ltaS* (N315 locus tag SA0674) has previously been mapped to the VraSR regulon as a positive regulator ^58^, however, direct control was not demonstrated ^38^. It is not surprising that WalKR and VraSR regulate *ltaS* transcription as both TCSs are intimately connected to cell wall homeostasis. WalKR maintains a balance of peptidoglycan cleavage that allows the cell to grow but not lyse ^10, 18^, and VraSR governing the cell wall stress stimulon in response to extracellular insult ^59, 60^. Together, LTA and wall teichoic acid are present in the Gram-positive cell wall in roughly equal proportion to peptidoglycan ^61^. Modulation of *ltaS* expression by two TCSs that integrate different signals, alongside post-translational regulation of the enzyme, presumably ensures tight and finely tuned control of *ltaS* activity, enabling co-ordination of LTA synthesis with peptidoglycan remodelling. Upon WalK activation, we observed a change in LTA chain length and reduction in modification. As LTA chain length is an intrinsic property of the LtaS enzyme that is dictated by the availability of lipid starter units ^62^, it is unlikely that the chain length differences observed upon activation of WalK are solely attributable to direct negative regulation of *ltaS*. We speculate that the global transcriptional rewiring of the cell upon WalK activation may affect the availability of lipid starter units, although this remains to be investigated.

In addition to its role in teichoic acid biosynthesis, we found that WalKR controls essential genes involved in the initiation of DNA replication. In *S. aureus*, DNA replication is initiated by binding of DnaA to AT-rich regions at the origin of replication, *oriC* ^63^. This process is tightly controlled, as mistiming of initiation results in aberrant cell division ^64^. We show that WalKR can both positively regulate the expression of *dnaA* and negatively regulate *hup.* The role of *hup* in staphylococcal DNA replication initiation has not previously been investigated but recently in *B. subtilis* the *hup* homologue *hbs* has been shown to promote initiation ^65^. We show this function is conserved in *S. aureus*; knocking down expression of *hup* resulted in reduced initiation of DNA replication. In addition to *hup,* two other regulators of DNA replication initiation have been characterised in *S. aureus*; Noc and CcrZ ^64^. Noc is a negative regulator of DnaA driven initiation ^47^, whereas CcrZ is a positive regulator ^64^. The mechanisms underlying the control of DnaA by these proteins are yet to be fully defined, however it is unlikely that Noc directly regulates *dnaA* expression ^47^ and CcrZ may act post-translationally, by phosphorylating an unknown intermediate factor ^64^. A recent investigation into the role of Noc in *S. aureus*, a suppressor mutant down regulating the activity of DnaA was identified with a deletion of a 56bp region [Figure 7C] within the 5’UTR of *dnaA* ^47^. We show this function is conserved in *S. aureus*; knocking down expression of *hup* resulted in reduced initiation of DNA replication. In addition to *hup,* two other regulators of DNA replication initiation have been characterised in *S. aureus*; Noc and CcrZ ^64^. Noc is a negative regulator of DnaA driven initiation ^47^, whereas CcrZ is a positive regulator ^64^. The mechanisms underlying the control of DnaA by these proteins are yet to be fully defined, however it is unlikely that Noc directly regulates *dnaA* expression ^47^ and CcrZ may act post-translationally, by phosphorylating an unknown intermediate factor ^64^. In a recent investigation into the role of Noc in *S. aureus*, a suppressor mutant down regulating the activity of DnaA was identified with a deletion of a 56bp region [Figure 7C] within the 5’UTR of *dnaA* ^47^. As this region encompasses one of two characterised WalR binding motifs upstream of the gene, we propose that the loss of positive WalR control through deletion of the binding site explains the observed decease in DnaA activity. Additionally, RNA-seq showed an overall increase in *dnaA* expression (1.05 Log_2_FC - WalK_T389A_ vs Wt) in exponential phase upon WalK activation, further highlighting the positive role WalR has on *dnaA* expression.

That WalR was found to positively regulate *dnaA* and negatively regulate *hup*, both of which are promoters of initiation, is somewhat counterintuitive. Knockdown of *walR* expression caused over-initiation, consistent with its negative control of *hup* but contradictory to its positive regulation of *dnaA* [Figure 8BC]. It may be that loss of HU is a dominant phenotype, reducing initiation even in the presence of higher levels of DnaA, or alternatively, that regulation of each gene is temporally distinct. The latter hypothesis is supported by our observation that *dnaA* regulation occurred during exponential phase whereas regulation of *hup* was in stationary phase [Figure 7C]. However, exactly how WalKR transcriptional regulation of both *dnaA* and *hup* works in concert with post-translational control mediated by HU, Noc and CcrZ to ensure tight, spatiotemporally accurate, control of initiation of DNA replication remains to be elucidated. It is of note that during the discovery of WalR in *B. subtilis* using a temperature-sensitive mutant, anucleate cells were observed at the non-permissive temperature. This observation hints at the possibility of a wider association between WalR and DNA replication in other Bacillota ^9^.

HU is a multifunctional protein, in addition to its role in DNA replication initiation, it also contributes to DNA compaction, introduces negative supercoils into DNA ^52^ and can impact localised gene regulation ^66^. As we observed opposite changes to DNA supercoiling caused by depletion of *walR* and *hup*, we propose that the negative regulation of HU by WalR causes relaxation of supercoiling and may lead to decreased compaction of cellular DNA. Transcriptional regulation of *hup* expression has not previously been described in *S. aureus*. However, qualitative western blots have shown HU to be continuously present in the staphylococcal nucleoid throughout all growth phases ^67^. Taking this into account, we propose that WalKR regulation of *hup* is not a binary switch, but rather provides tuneable control of this essential system. Intriguingly, in *Mycobacterium tuberculosis* the serine/threonine kinase PknB negatively regulates HU DNA binding through phosphorylation ^68^ and in *S. aureus*, PknB mediated phosphorylation activates WalR ^69^. Furthermore, a PknB phosphorylation site on HU has been experimentally identified in *S. aureus* ^70^. Therefore, PknB may directly repress HU DNA binding through post-translational modification and indirectly represses the expression of *hup* through activation of WalR.

The reason for essentiality of WalKR differs across Bacillota species ^71^. In *S. aureus,* essentiality is proposed to result from polygenic control of non-essential autolysins involved in the cleavage and relaxation of peptidoglycan crosslinks, allowing expansion of the cell wall ^6^. Here, we extend this hypothesis, showing WalKR direct control of at least five essential genes (*rplK*, *hup, ltaS, prs* and *dnaA*) not directly involved in peptidoglycan biosynthesis but intimately linked with cell growth and cell division. Thus, we propose that WalKR essentiality arises though polygenic co-ordination of multiple cellular processes; ribosome assembly, peptidoglycan homeostasis, LTA polymerisation, DNA topology, and the initiation of DNA replication, ultimately making WalKR an indispensable link between cell wall homeostasis and DNA replication. We propose a model in which WalK senses a currently unknown ligand during logarithmic growth through the extra-cytoplasmic PAS domain, resulting in autophosphorylation, dimerisation, and maximal interaction between WalK and WalR [Figure 10A]. The level of WalK activation can also be dynamically tuned in response to the metalation state of the intracellular PAS domain ^28^. WalK:WalR interaction allows phosphotransfer to WalR residue D53, whilst a second WalR site T101 can be phosphorylated through the PknB kinase (recognises muropeptide fragments^72^). Phosphorylated WalR binds the cognate recognition motifs of its direct regulon as a dimer, causing either negative or positive changes to gene expression, primarily dependent on the position of the binding motif in relation to the transcriptional start site [Figure 10B]. The direct WalR regulon has three broad functions; i) control of cell wall metabolism through regulation of a suite of autolysins governing peptidoglycan homeostasis and fine tuning of LTA biosynthesis through negative regulation of lipoteichoic acid synthase; ii) linking initiation of DNA replication to cell wall homeostasis through regulation of *dnaA* and *hup*; iii) signal amplification through modulation of transcription factors, selected other TCSs, and via negative regulation of DNA binding protein HU [Figure 7B]. Together with the downstream effects of changes in cell wall metabolism, signal amplification drives changes in expression of the indirect WalKR regulon, producing a large shift in cellular transcription that includes the increased expression of virulence factors and metabolic genes [Figure 10C].

**Figure 10.**
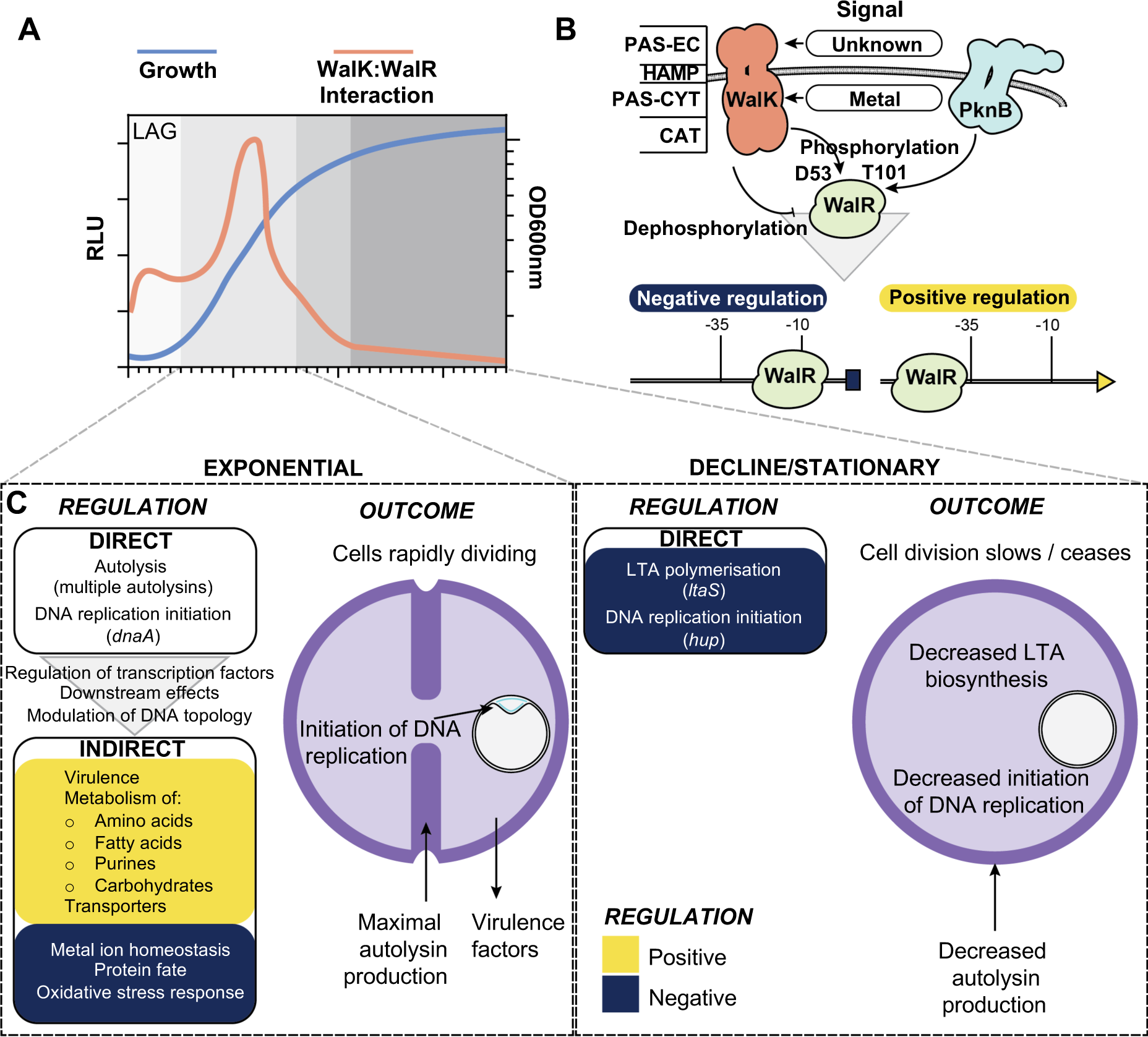
A model of the role of WalKR in cell division. A) Changes in the interaction of WalK with WalR throughout growth. B) Mechanism of signal transduction and transcriptional control of WalKR. −35 and −10 denote promoter regions. PAS-EC: Per-Arnt-Sim extracellular, HAMP: present in Histidine kinases, PAS-CYT: PAScytoplasmic, CAT: catalytic domain. C) Transcriptional changes and their outcomes for the direct and indirect WalKR regulon during different growth phases. Yellow and dark blue denote positively, and negatively, regulated gene sets, respectively.

WalKR has long been considered a promising target for the development of novel anti-Gram-positive agents ^73^, although to date no WalKR targeted compounds have been successfully developed. Despite this, the system remains a viable target for conventional antibacterial chemotherapy due to its role as a signal integrating nexus of essential cellular functions and the presence of two PAS domains capable of binding small molecules inside and outside of the cell ^18, 28^. Increasingly, alternative strategies to traditional antibacterial chemotherapy are being explored, including the development of so called “antibiotic resistance breaking (ARB) compounds” that re-sensitise resistant strains to existing antibiotics ^74, 75^. Here, we found that mutationally activated WalKR ‘up’ mutants were more sensitive to three different antibiotic classes targeting the cell wall: oxacillin, tunicamycin, and vancomycin. In contrast, recent work conducted using the methicillin sensitive *S. aureus* (MSSA) strain ATCC29213 found the opposite; a WalKR ‘down’ mutation (WalR_T101M_) increased tunicamycin sensitivity ^76^. Tunicamycin is a dual targeted antibiotic, binding to MraY and TarO, which belong to the peptidoglycan recycling and wall teichoic acid synthesis pathways, respectively ^77^. In MSSA ATCC29213, decreased WalKR activity results in indirect downregulation of enzymes within the peptidoglycan recycling *mupG* operon, as well as direct downregulation of autolysins. Together these gene expression changes starve MraY of precursor molecules, making the enzyme exquisitely sensitive to tunicamycin ^76^. In MRSA NRS384 the genes of the *mupG* operon were not differentially regulated upon WalKR activation. Consequently, we conclude that tunicamycin sensitivity in NRS384 ‘up’ mutants likely arises through a different mechanism. More broadly, differences in the indirect WalR regulon between MRSA and MSSA may reflect regulatory adaptions necessary to accommodate the exogenous, *mecA* encoded, PBP2a transpeptidase into the complex process of cell wall homeostasis ^78^. Here, we found that mutational stimulation of normal WalKR activity rendered NRS384 MRSA susceptible to oxacillin despite the presence of *mecA*, opening an alternative avenue for future WalKR focused drug development; the discovery of ARB compounds that phenocopy WalKR activating mutations.

## Materials and Methods

### Strains, oligonucleotides, media, and reagents

Bacterial strains and plasmids are listed in Table S5. Oligonucleotides (IDT) used in this study are listed in Table S6. *Escherichia coli* were routinely cultured in LB (Merck) or on L-agar (1.5% [w/v] agar added) unless stated otherwise. *S. aureus* were routinely grown on Brian Heart Infusion (BHI) agar (Bacto, BD Biosciences) or Sheep blood agar. When cultured in broth, they were grown in Brain heart infusion broth Trypticase soy broth (TSB, Oxoid), or LB with shaking at 200 rpm. For selection, antibiotics (Sigma) were added at the following concentrations for *E. coli* (*E.c*) and *S. aureus* (*S.a.*): Ampicillin 100 µg/mL^-1^ – *E.c*., Kanamycin 50 µg/mL^-1^ – *E.c*./*S.a*., Chloramphenicol 10 µg/mL^-1^ – *E.c.*/*S.a*. Restriction enzymes, Phusion DNA polymerase, and T4 ligase were purchased from New England Biolabs. Phire Hotstart II DNA polymerase for colony PCR was purchased from Thermo Fisher.

### *S. aureus* site-directed mutagenesis by allelic exchange

Upstream and downstream regions of the point mutation for *walK*^T389A^ (IM7/IM120/IM121/IM10), *walR*^T101A^ (IM31/IM232/IM233/IM10) or *spa* deletion (IMT275/IMT353/IMT354/IMT278) were PCR amplified and then a SOE-PCR was performed on the gel extracted template to generate an amplicon for SLiCE cloning into pIMAY-Z ^79^. This yielded plasmids pIMAY-Z *walK*^T389A^, pIMAY-Z *walR*^T101A^ and pIMAY-Z Δ*spa*. To construct pIMAY-Z *walK*^Y32C^, genomic DNA from a sectored mutant of NRS384 Δ*yycHI* (containing an additional *walK*^Y32C^ mutation) was amplified with primers IM107/IM10 and the amplicon cloned as described above. Construction of isogenic mutants of NRS384 by allelic exchange was performed as described previously ^79^. The WalK enhancing mutations could visually be discriminated from the wild type due to reduced colony size and opacity. For the WalR^T101A^, the mutation was screened by colony PCR (70°C annealing temperature) with primers IM233/IM181. From putative mutants, genomic DNA was extracted from 1 ml of overnight culture (DNeasy Blood and Tissue Kit—Qiagen) pre-treated with 100 μg of lysostaphin (Sigma cat. no. L7386) and sequenced on an Illumina NextSeq by the Doherty Applied Microbial Genomics facility (University of Melbourne). Resultant reads were mapped to a NRS384 reference genome ^80^ and mutations identified using Snippy (https://github.com/tseemann/snippy).

### Construction of pRAB11-FLAG

To construct a vector for the C-terminal FLAG tagging of *S. aureus* proteins, the anhydrotetracycline inducible vector pRAB11 ^34^ was digested with KpnI to linearise and gel extracted. The 6.4kb vector was then amplified with primers IM512/IM513 to add in a consensus ribosome binding site (IM512), a 1xFLAG tag and downstream *tonB* transcriptional terminator (IM513). The amplimer was digested with KpnI, gel extracted, re-ligated to yield pRAB11-FT. The sequence of the plasmid was verified by sequencing on the Illumina platform. To clone into pRAB11-FT, the vector was digested with KpnI, gel extracted and used as template with primers IM514/IM515 to amplify the vector backbone. Response regulators (WalR - IM516/IM517), (SaeR - IM518/IM519), (VraR - IM520/IM521), (HptR - IM522/IM523) were amplified from the start codon and omitting the stop codon with NRS384 genomic DNA. The products were gel extracted, SLiCE cloned into amplified pRAB11-FT and transformed into IM08B, yielding pRAB11:*walR*^FLAG^, pRAB11:*saeR*^FLAG^, pRAB11:*vraR*^FLAG^ and pRAB11:*hptR*^FLAG^. The plasmids were electroporated into NRS384Δ*spa*.

### Construction of pIMC8-YFP reporter strains and assay for YFP activity

Promoter regions for *sasD* (IM1127/IM1107), *sle1* (IM1108/IM1109), *P602* (IM1129/IM1110), *ltaS* (IM1111/IM1112), *ssaA* (IM1130/IM366) *isaA* (IM1128/IM364 were PCR amplified from NRS384 genomic DNA and gel extracted. The vector pIMC8-YFP ^28^ was digested with KpnI, gel extracted and PCR amplified with IM1/IM385. The amplified promoters and vector were SLiCE cloned, transformed into IM08B and subsequently electroporated into NRS384, NRS384 Δ*yycHI* or NRS384 *walK*^Y32C^. Mutations disrupting the WalR motif (1^st^ TGT to CCC) were introduced by SOE-PCR with the bracketed primers sets for *hpt* (IMT300/IMT301); *sle1* (IM1108/IM1115; IM1114/1109), *P0602* (IM1129/1119; IM1118/IM1110), *ltaS* (IM1111/IM1117; IM1116/IM1112), *ssaA*^CCCI^ (LS371/LS376; LS375/LS372), *ssaA*^CCCII^ (LS371/IM1121; IM1120/LS372), *isaA*^CCCI^ (IM1128/IM1123; IM1122/IM364), *isaA*^CCCII^ (IM1128/IM1125; IM1124/IM364). The resultant mutated promoters were gel extracted, SLiCE cloned, transformed into IM08B and subsequently electroporated into NRS384. For the *sasD* promoter the mutation was incorporated into the reverse primer (IM1113) in combination with IM1127. To assess YFP production, each strain was grown in 5 ml of LB containing 10 µg/mL^-1^ chloramphenicol in a 50 ml tube for overnight at 37°C with shaking at 200 rpm. The culture was diluted 1:100 in 5 ml of fresh LB containing chloramphenicol and incubated overnight. The fluorescence (excitation 512 nm, emission 527 nm) of each strain (200 µl – Nunc black well plates) was read in triplicate using an Ensight multimode plate reader (PerkinElmer) set to 100 flashes per well. Resultant data was plotted using the GraphPad Prism (v9.3.1) software package.

### Production and purification of proteins

For production of WalR, Rosetta 2 (DE3) pET28(a):*walR* was grown in 2 L of autoinduction media ^81^ at 25°C for four days with vigorous shaking, and cells were harvested by centrifugation. For production of VraR Rosetta 2 (DE3) pET21(d): *vraR* was grown in 2 L of LB at 37°C with vigorous shaking to an OD600 nm of 0.6, chilled on ice for 10 min then induced with 1 mM IPTG. Subsequently, cells were grown for a further 16-20 h at 18 °C with vigorous shaking and harvested by centrifugation. For purification of both proteins, cell pellets were resuspended at 3 ml/g in buffer A (50 mM NaH_2_PO_4_ [pH 8.0]) with 300 mM NaCl and 10 mM imidazole. To enhance lysis and prevent proteolysis, 7000 U chicken egg-white lysozyme (Sigma), two complete EDTA-free protease inhibitor tablets (Roche) and 20 U DNase I (NEB) were added. Cells were lysed by sonication and lysates were clarified by centrifugation at 30,000 x *g* for 30 minutes at 4°C. The cleared lysate was loaded onto a 25 mL free-flow gravity column (GeneFlow) packed with 3 ml TALON® Metal Affinity Resin (Takara Bio), washed with 10 column volumes (CV) of buffer A containing 20 mM imidazole and 2 M NaCl. The protein was eluted in 2 CV of buffer A containing 150 mM imidazole and 300 mM NaCl and dialysed overnight 6 kDa MWCO, CelluSep) into storage buffer (25 mM Tris [pH 8.0], 300 mM NaCl, 20 mM KCl, 10 mM MgCl_2_, 1 mM DTT, 5% glycerol). The proteins were spin concentrated to 5 mg / ml (3 kDa MWCO, vivaspin 20, Sartorius), aliquoted, snap frozen in liquid nitrogen, and stored at −80°C until required.

### Electrophoretic mobility shift assays (EMSAs)

DNA duplex probes for electrophoretic mobility shift assay binding assays were generated by annealing two 70 bp single-stranded oligonucleotides, one of which was labelled with Cy5 fluorophore on the 5’ end (Table S6). Annealing was performed in duplex buffer (30 mM HEPES [pH 7.5], 100 mM potassium acetate) by heating equimolar concentrations of complimentary single-stranded oligonucleotide at 94°C for two min and then cooling to room temperature over 30 min. Duplexes were subsequently gel-extracted from a 2% (w/v) agarose 1xTBE gel. Binding reactions were performed in a final volume of 25 μl of binding buffer (25 mM Tris [pH 8.0], 300 mM NaCl, 20 mM KCl, 10 mM MgCl_2_, 1 mM DTT, 4 µg BSA, 0.5 µg salmon sperm DNA). Initially, varying concentrations of WalR/VraR were incubated for 5 min at 25°C in binding buffer, then DNA probe was added to a final concentration of 16 nM and the reaction was incubated for a further 15 min. After incubation, reactions were mixed (5:1) with 6x Orange G loading buffer ^82^ and 5 µl was electrophoresed on an 8% polyacrylamide native gel (29:1 acrylamide:bisacrylamide ratio) in 1xTBE at 4 °C. Bands were visualised using a GE Amersham 600 imager in the Cy5 channel with a 10 min exposure time.

### Total RNA extraction and rRNA depletion

A 10 ml LB (50 ml tube) culture was grown overnight at 37°C with shaking at 200rpm. The saturated culture was diluted 1:100 into fresh 10 ml TSB and grown to an OD600nm of 0.8-1.0. A 5 ml aliquot of culture was removed and added to 10 ml of RNAprotect Bacteria Reagent (Qiagen) and mixed by vortexing. The sample was incubated at room temperature for 5 min. Cells were then harvested by centrifugation (7,000*xg*/5min/22°C), the supernatant discarded and the cell pellet resuspended in 1 ml of TRIzol (Invitrogen). Cells were lysed by bead beating (Precellys 24 instrument - 6000 rpm, 1 min, 100 µm zirconium beads) and cell lysates were clarified by centrifugation (20,000*xg*/10 min/4°C). Subsequently, 700 µl of supernatant was removed and mixed with 700 µl of ethanol. RNA was extracted using a Direct-Zol RNA miniprep plus kit (ZymoResearch) according to manufactures instructions, including the on-column DNase I treatment step. Following RNA extraction, an additional DNA removal step was performed using a TURBO DNA-free kit (Invitrogen) according to manufacturer’s instructions. The absence of DNA was accessed *by gyrB* PCR (IM1020/IM1021) on 1µl of RNA template, yielding no amplification. RNA quality was determined on the Bioanalyser (Agilent) with all yielding a RIN score of above 8. For each strain three independent RNA extractions were made. A 5 µg aliquot of total RNA was depleted for rRNA with the mRNA then converted into cDNA with the Scriptseq complete bacteria kit (Epicentre). The libraries were sequenced on the Illumina HiSeq platform for 50 bp single end reads.

### RNA-seq data analysis

RNA-seq data was analysed using the *S. aureus* USA300 FPR3757 reference genome (accession number: NC_007793) and Kallisto ^83^, a kmer based pseudoalignment tool, with analysis and visualization using Degust [https://github.com/drpowell/degust]. Degust uses Voom/Limma ^84^ and generates an interactive website to analyse and explore the data.

### Preparation of samples for ChIP-seq

An overnight culture (10 ml LB in 50 ml tube) of each strain NRS384Δ*spa* containing either pRAB11: *walR*^FLAG^ / *vraR*^FLAG^ / *hptR*^FLAG^ / *saeR*^FLAG^ or pRAB11^FLAG^ only were grown at 37°C with shaking at 200 rpm. Overnight cultures were diluted 1:100 in fresh LB (100 ml) in a 1 L baffled flask and grown to an OD600nm of 0.5, induced with 100 ng/mL^-1^ of anhydrotetracycline and grown for a further hour. Cells were crosslinked by direct addition of methanol free formaldehyde (Pierce) to cultures (final concentration of 1% (v/v)) and incubated with gentle mixing (rotating platform) for 15 min at room temperature. The crosslinking reaction was quenched by addition of glycine to a final concentration of 400 mM and incubated a further 15 min. Cells were pelleted (7,000xg/10min/4°C), washed three times with ice-cold Phosphate buffered saline (PBS) and the pellet stored at −80°C. For cell lysis, cells were suspended in 1ml of lysis buffer (50 mM Tris-HCl [pH 7.4], 150 mM NaCl, 0.1% Triton X-100, 100 μg/mL^-1^ lysostaphin (Ambi), 500 µg/mL^-1^ RNaseA, complete mini protease inhibitor EDTA free containing 100 um zirconium beads) and incubated for 20 min at 37°C. The weakened cells were then disrupted using bead beating (6000 rpm for 1 min; Precellys 24 instrument) and the cell lysate clarified by centrifugation (20,000*xg/*10min/4°C). The cation concentration of the lysate was adjusted with MgCl_2_ and CaCl_2_ to 100 mM. The DNA was sheared by the addition of a range of DNase I (NEB) concentrations (0.5 – 2U), in a 200 μL volume, followed by incubation at 37°C for 10 min. The reaction was quenched by the addition of EDTA to 50 mM on ice. The degree of DNA fragmentation was assessed by electrophoresis of samples on a 2% (v/v) agarose TAE gel. Samples showing maximal fragmentation at between 100-300 bp were subjected to immunoprecipitation. Lysis buffer (made up to 4 ml - omitting lysostaphin and RNase A) containing 10 µg of M2-anti FLAG antibody was added and incubated overnight at 4°C on a rotating platform in a 15 ml tube. A 100 μl aliquot of Protein G agarose (Pierce) was added to the lysate and incubated a further 2h at room temperature. The sample was centrifuged (2,500*xg*/3 min/22°C) and the agarose pellet washed three time with 4 ml of IP buffer (25mM Tris.Cl [pH 7.2], 150mM NaCl) and finally resuspended in 200 µl of elution buffer (10mM Tris.Cl [pH 8], 1mM EDTA, 1% SDS containing 100µg of proteinase K). Crosslinks were reversed incubation for 2h at 37°C and then 9h at 65°C with shaking at 1400 rpm. Finally, eluted DNA was cleaned up by PCR purification (QiaQuick PCR purification kit, Qiagen). DNA libraries were prepared with NEBNext® Ultra™ II DNA Library Prep Kit for Illumina and sequenced using an Illumina MiSeq.

### ChIP-seq read mapping, peak identification, and motif searching

MiSeq reads from each of the five experiments (pRAB11: *walR*^FLAG^ / *vraR*^FLAG^ / *hptR*^FLAG^ / *saeR*^FLAG^ or pRAB11^FLAG^ only) were first mapped to the *S. aureus* USA300 NC_007793 reference chromosome using *samtools*. The resulting *.bam* and *.sam* files were used to create tag counts (*i.e.,* mapped reads) with the *makeTagDirectory* script within *homer* ^85^. The *homer* peak identification tool (*findPeaks*) was extensively explored but high levels of background reads were detected and precluded further use of *homer* for peak detection. An alternative strategy was developed by building read coverage plots for viewing in Artemis ^86^, using *samtools* and the command *% samtools depth -aa [target_file_name].bam [subtraction_file_name].bam | cut −f3,4 | perl-nae ’use List::Util qw(max); print max(0, $F[0]-$F[1]),"\n";’ > [target_peaks].userplot*. This created a coverage plot of those regions of the *S. aureus* USA300 NC_007793 chromosome specifically bound by a given response regulator, relative the response regulator sequence reads in the subtraction set. These subtraction sets were a concatenation of a random selection of 20% of the sequence reads for the three response regulators and the plasmid-only control combined. Thus, for peak discovery of WalR-binding sites, an Artemis userplot was prepared from *vraR*^FLAG^ / *hptR*^FLAG^ / *saeR*^FLAG^ and pRAB11^FLAG^ subtracted from *walR*^FLAG^. The same method was used to generate userplots and discover binding peaks for the remaining three response regulators. The ‘*create feature from graph function*’ in *Artemis* was then used to define chromosome regions represented by the subtraction coverage plots. These regions were mapped to the NRS384 genome in Geneious Prime (v2023.03) and searched using TGTNNNNNNNNTGT +/− 5bp as input. Output sequences were combined with motif regions +/− 5bp for the following previously experimentally validated WalR regulon members; SAUSA300_0739, SAUSA300_0955, SAUSA300_2051, SAUSA300_2253, SAUSA300_2503 ^12, 24^ and input to WebLogo ^87^. The resultant sequence logo was converted to IUPAC code and used to search the NRS384 genome using Geneious Prime (v2023.03).

### Mapping TSS in relation to WalR binding sites

The *S. aureus* NRS384 genome was annotated in Geneious Prime (v2023.03) with predicted transcriptional start sites (TSS) as defined in a previous study ^39^. The 500 bp upstream of a predicted TSS were extracted and manually annotated with −35 and −10 elements and predicted WalR binding sites. T-tests (run in Stata v16.0) were used to test associations between WalR binding site position and orientation, with gene expression data from RNA-seq.

### Essentiality analysis

Essentiality of *S. aureus* genes was called if a locus had been described as essential in two previous studies ^41, 42^ and did not harbour a transposon insertion in the Nebraska transposon library ^43^. Genes were defined as belonging to the core genome if they were present in every strain of the *Aureo*Wiki orthologue table (https://aureowiki.med.uni-greifswald.de/download_orthologue_table)^88^. Data from RNA-seq, ChIP-seq, and the essentiality and core genome analysis were integrated in R (v4.0.3, https://www.r-project.org/) with RStudio 2022.02.0+443 using dplyr (v1.0.8) and tibble (v3.1.6), then visualised using UpSetR (v1.4.0) ^43^ with ggplot2(v3.3.5).

### Growth curves, other phenotypic testing

For measurement of growth curves, *S. aureus* was grown overnight in BHI broth at 37°C and subsequently diluted into fresh BHI broth to an OD600_nm_ of 0.05. Growth was measured for 8 hours at 37°C with 300 rpm dual orbital shaking in a 96 well plate (Corning) using a Clariostar Plus (BMG) plate reader.

### Antibacterial susceptibility testing

Minimum inhibitory concentrations of antibacterial agents were determined by broth microdilution; bacteria were exposed to 2-fold serial dilutions of antibacterial agents in Mueller-Hinton broth 2 (BBL, BD) according to the guidelines provided by the Clinical and Laboratory Standards Institute. Vancomycin susceptibility was assessed using gradient plates as previously described ^89^.

### Lysostaphin sensitivity

An overnight 5ml BHI culture of each strain was diluted 1:100 in an Eppendorf tube containing fresh BHI containing different final concentrations (0 – 1.6 µg/mL^-1^) of lysostaphin (Ambi). Cells were then incubated statically for 90 min at 37°C in a water bath with the CFU/mL^-1 determined^ by dilution and spot plating onto BHI agar. Plates were incubated for 18 h at 37°C before enumeration.

### LTA extraction, purification, and analysis by PAGE

For extraction and analysis of LTA, 50 mL of LB was inoculated with a single colony of NRS384, NRS384 Δ*yycHI,* NRS384 *walK*_T101A_, NRS384 *walK*_T389A_, or RN4220, and grown at 37°C with vigorous shaking for 18 h. Cells were harvested from 30 ml of saturated culture by centrifugation of (5000 *x g*, 10 min) and LTA was extracted and analysed by PAGE as described previously ^90^.

#### CRISPRi constructs and knockdown analysis

To generate pSD1 CRISPRi knockdown constructs for *walR, hla* and *hup,* primers corresponding to the previously described for *walR* (IM1180/IM1181) and *hla* (IM1182/IM1183) were synthesised from Zhao *et al* ^91^. Knockdown primers for *hup* (targeted the region overlapping and upstream of the *hup* start codon (IM1559/IM1560)) were designed with the annealed primer pairs for *walR*, *hla* and *hup* cloned into the SapI site of pSD1 as described previously ^92^. Plasmids were then transformed into NRS384. Overnight cultures (5ml LB, chloramphenicol 10 μg/mL^-1 in^ 50 ml tubes) were then diluted in fresh media 1:100 containing different concentrations of aTc to induce expression of dCAS9, with the optical density of the cultures followed.

RNA isolation: RNA was isolated from 1ml of cells (induced with either 0 or 100 ng/mL^-1^ of aTc) after 5h of growth, as described above. A 1ug aliquot of total RNA was converted into cDNA with Superscript IV and random hexamers as described previously ^93^. For RT-qPCR, 1 μl of cDNA was used as template with primers for *gyrB* (IM1020/IM1021), *hla* (IM1026/IM1027), *walR* (IM1153/IM1154) and *hu* (IM1586/IM1587) with Luna Universal qPCR Master Mix (NEB) on a Quantstudio 1 PCR machine. The data was normalised to the *gyrB* gene and analysed with the ΔΔ CT method ^94^.

Plasmid isolation: Plasmid DNA was isolated from 10 ml of cells at 5h post induction (induced with 0, 10, 50 and 100 ng/mL^-1^ aTc). Cells were centrifuged 7,000*xg*/2min pellet was washed in 1ml of PBS and then resuspended in 400 μl of the resuspension buffer (Monarch Miniprep kit – NEB) containing 50 μg of lysostaphin. The cells were lysed at 37°C for 30 min and then processed following the kit instructions through one column with elution in 30 µl of elution buffer. DNA was quantified with the Qubit BR DNA quantification kit and normalised to 15 ng/µl, with the normalised loading (150ng of purified plasmid) assessed on a 1% TAE gel. Subsequent 1% agarose gels (in 2xTBE) containing 2.5 µg/mL^-1^ chloroquine were run as described by Cameron *et al* ^95^. Gels (10cm) were run at 10V for 16h which were washed twice (30 min each wash) in dH_2_O and then stained with Sybr Gold for 30 min and subsequently imaged.

Genomic isolation: Genomic DNA was isolated from the equivalent of OD600nm of 5 after 5h of growth (induced with 0 or 100 ng/mL aTc). The cell pellet was washed with 1ml of PBS and resuspended in 90 μl of PBS containing 5 µl of 20 mg/mL^-1^ RNase A, 50 µg of lysostaphin and 100 µl of the tissue lysis buffer (Monarch Genomic DNA purification kit, NEB). Cells were lysed at 37°C for 30 min and then processed following the manufactures instructions.

### Analysis of *ori*-*ter* ratios

To measure to *ori*-*ter* ratios, genomic DNA prepared after CRISPRi knockdown (as above) was sequenced using the Illumnia NextSeq (by the Doherty Applied Microbial Genomics facility, University of Melbourne). Illumina reads were processed and analysed using iRep, as previously described (v1.1 https://github.com/christophertbrown/iRep ^96^).

#### Construction of pSmBIT and pLgBIT split luciferase vectors

The vector pRAB11(pC194 replicon) ^34^ was modified by PCR to restore the consensus *tetO* upstream of the *tetR* gene (IM1290/IM1291) ^97^. As described previously, this reduced the impact of elevated level TetR production and allowed leaky expression of the target gene in the absence of aTc. The above 6.4 kb PCR product was gel extracted, treated with SLiCE and transformed into *E. coli* IM08B, yielding pRAB11*. To introduce a consensus ribosome binding site and 9 nucleotide spacer before the start codon (<u>AGGAGG</u>AATTGGAAA) downstream of the two *tetO* sites (proceeding the gene of interest), pRAB11* was first digested with KpnI and gel extracted. This was used a template in a PCR (IM513/IM1355), the product digested with KpnI, gel extracted and ligated. The ligation product was transformed into IM08B yielding pRAB11*RBS. The *tetR*-*RBS fragment was digested from pRAB11*RBS (SphI/KpnI) and ligated into complementary digested pCN34 (pT181 replicon) ^98^ yielding pCN34*RBS. Both pRAB11*RBS and pCN34*RBS were digested with KpnI, gel extracted and used as template in a PCR (IM515/IM1356). The following combinations were combined with 50 ng of each 1. pRAB11*RBS PCR and LINKER(GSSGGGGSGGGGSSG)-SmBIT gBlock. 2. pCN34*RBS PCR and LINKER-LgBIT gBlock. gBlock sequences were codon optimised for *S. aureus* The SLiCE reactions were transformed into IM08B yielding either pSmBIT or pLgBIT, with both vectors were fully sequenced to validate.

#### Cloning into split luciferase vectors

Either pSmBIT and pLgBIT were digested with KpnI, gel extracted and used as template for PCR with primers IM515/IM1360. The pSmBIT or pLgBIT amplimers were combined with amplified open reading frames with stop codon removed and tailed with 5’-GATAGAGTATGATGAGGAGGAATTGGAAA-3’ forward or 5’-GAACCACCACCACCACTAGAACC-3’ sequences complementary to the vector.

WalR and WalK alleles were PCR amplified with IM1363/IM1364 and IM1365/IM1366, respectively, then SLiCE cloned into pSmBIT (for *walR* alleles) or pLgBIT (for *walK* alleles) and transformed into IM08B. For *S. aureus* transformations, at least 1 µg of pLgBIT(+*walK* allele) and pSmBIT(+ *walR* allele) were purified from IM08B and co-electroporated into NRS384 with selection on BHI agar containing 10 ug/mL^-1^ chloramphenicol and 50 ug/mL^-1^ kanamycin.

#### Chromosomal tagging of WalR-SmBIT and WalK-LgBIT

To access the functional interaction of WalR/WalK under native levels of protein of production, the native copy on the chromosome was tagged with SmBIT for WalR and LgBIT for WalK. The regions of DNA were assembled as follows. 1. *walR*-SmBIT-*walK*: *walR*-SmBIT was amplified from pSmBIT-*walR* with IM107/IM1517 and a downstream fragment encompassing *walK* was amplified with IM1516/IM10 from NRS384 genomic DNA. Both were gel extracted and joined by SOE-PCR. 2. *walK*-LgBIT-*yycH*: *walK*-LgBIT was amplified from pLgBIT-*walK* with IM7/IM1519 and a downstream fragment encompassing 500bp of *yycH* was amplified with IM1518/IM44 on NRS384 genomic DNA. Both were gel extracted and joined by SOE-PCR. Either amplimer was SLiCE cloned into pIMAY-Z and transformed into IM08B. The above cloning steps and *S. aureus* allelic exchange was performed as described by Monk and Stinear ^79^. Presence of *walR-*SmBIT and *walK-*LgBIT were screened by colony PCR with IM1360/IM1368 and IM1360/IM44, respectively. Genomic DNA was isolated from the strains and whole genome sequenced.

#### Growth and luciferase curves

An overnight 5 ml LB containing antibiotics (in a 50ml tube) were grown overnight. The culture was diluted 1:100 in fresh LB supplemented with chloramphenicol and kanamycin including a 1:5000 dilution of the Nano-Glo® Luciferase Assay Substrate (Promega). Preliminary growth curves in the presence of the substrate showed no impact on growth at this concentration. The culture was then dispensed in triplicate (200 µl) black/clear bottom 96 well plates (Cat no. 165305, Themofisher). The plates were sealed with MicroAmp™ Optical Adhesive Film (Thermofisher) and incubated at 37°C with dual orbital shaking at 300 rpm (Clariostar Plus, BMG). Every 10 minutes the plate was read at OD600nm and light emission (1s exposure) collected over an 8 h period.

#### Bacterial luciferase reporter plasmid

To construct pIMK1-LUX, the Listeria phage integrase vector pIMK was digested with SphI/BglII to excise the PSA integrase and replace it with the PCR amplified (IM1241/IM1242) low copy number pSK41 replicon from pLOW, yielding pIMK1. Vectors pIMK1 and pPL2*lux* were then digested with SalI/PstI and the gel extracted pIMK1 backbone ligated to the bacterial luciferase operon from pPL2*lux*. The vector pIMK1-LUX produces exact promoter fusions which can be cloned into the SalI/SwaI digested vector, as described previously ^99^. Promoters for *walR* (IM1222/IM32), *sasD* (IM1216/IM248 (Wt) or IM1217(ccc)), *sle1* (IM1295/IM1115; IM1114/IM1296), *0602* (IM1294/IM1119; IM1118/IM1062), *ltaS* (IM1218/IM1117; IM1116/IM1219), *ssaA^CCCII^* (IM1297/IM1121; IM1120/IM1298), *isaA^CCCI^* (IM1220/IM1123; IM1122/IM1221), *isaA^CCCII^* (IM1220/IM1125; IM1124/IM1221), *tarF* (LS451/LS448; LS449/LS452), *tagG* (LS457/LS454; LS455/LS458), *dnaA* (IM1745/IM1744; IM1743/IM1746), *dnaD^CCCI^* (LS471/LS466; LS467/LS472), *dnaD^CCCII^* (LS471/LS468; LS469/LS472,) *rplK* (IM1734/LS143; LS144/IM1735), *hup* (IM1289/IM1290; IM1291/IM1292), *dnaA^CCCI^* (LS443/LS440; LS441/IM1746), *dnaA^CCCII^* (LS443/LS438; LS439/IM1746), *dnaA^CCCII^* (LS443/IM1743; IM1744/IM1746), and *prs* (IM1285/LS139; IM140/IM1288) were PCR amplified from genomic DNA with either the outer set (Wt promoter) or SOE-PCR with the four primers (ccc promoter). The amplimers were digested with SalI, gel extracted and cloned into the above double digested vector. Plasmids isolated from IM08B were transformed into NRS384 Wt. An overnight 5ml LB culture with kanamycin was diluted 1:100 in fresh LB containing kanamycin with growth curves and light emission measured as described above for the split luciferase with the substrate omitted.

## Data visualisation

Graphs were generated in R (v4.0.3, https://www.r-project.org/)or GraphPad Prism (v9.3.1) software packages.

## Data availability

The DNA sequence reads for ChIP-seq and RNA-seq have been submitted to the NCBI Gene Expression Omnibus (GEO) repository under accession number GSE212321. DNA sequence reads for *ori*-*ter* analysis have been submitted to NCBI under Bioproject ID PRJNA886746.

## Acknowledgements

We are grateful to the following colleagues of for their advice, assistance, and input into this work: Justin Clarke, Charlie Higgs, Danielle Ingle, Karinna Saxby. T.P.S. is supported by NHMRC Research Grant (APP1105525) and NHMRC project grant GNT1145075. H. E. G. M. is supported through an NHMRC ideas grant 2003192. A.M.T. is supported by an Australian Government Research Training Program scholarship.

