## Supplemental data file for "The two-component regulator WalKR provides an essential link between cell wall homeostasis with DNA replication in *Staphylococcus aureus*"

- Supplementary Data File -

Liam K. R. Sharkey[1], Romain Guerillot[1], Calum Walsh[1], Adrianna M. Turner[1], Jean Y. H. Lee[1], Stephanie L. Neville [1], Stephan Klatt[2], Sarah L. Baines[1], Sacha Pidot[1], Fernando J. Rossello[3,4], Torsten Seemann[1,5], Hamish McWilliam[1], Ellie Cho[6], Glen P. Carter [1], Benjamin P. Howden[1,5], Abderrahman Hachani[1], Christopher A. McDevitt[1], Timothy P. Stinear[1,5],* and Ian R. Monk[1],*

1. Department of Microbiology and Immunology, Doherty Institute for Infection and Immunity, University of Melbourne, Melbourne, Victoria, 3000, Australia
2. The Florey Institute of Neuroscience and Mental Health, Melbourne Dementia Research Centre, The University of Melbourne, Parkville, Victoria, 3010, Australia.
3. University of Melbourne Centre for Cancer Research, The University of Melbourne, Melbourne, Victoria, 3000, Australia.
4. Australian Regenerative Medicine Institute, Monash University, Melbourne, Victoria, 3800, Australia.
5. Centre for Pathogen Genomics, Department of Microbiology and Immunology, Doherty Institute for Infection and Immunity, University of Melbourne, Melbourne, Victoria, 3000, Australia
6. Biological Optical Microscopy Platform, University of Melbourne, Melbourne, Victoria, 3000, Australia

* Joint senior authors

| **Supplementary figures and tables:** | | **Page** |
|---|---|---|
| Figure S1 | Growth in LB of C-terminally tagged WalR-SmBIT or WalK-LgBIT. | 2 |
| Table S1 | The indirect WalR regulon defined by RNA-seq | 3-9 |
| Figure S2 | ChIP-seq reveals binding sites of response regulators. | 10 |
| Table S2 | Putative response-regulator binding sites identified by ChIP-seq. | 11-13 |
| Figure S3 | Generation of an *S. aureus* WalR binding motif. | 14 |
| Table S3 | The predicted WalR direct regulon. | 15-18 |
| Figure S4 | Diversity of WalR motifs. | 19 |
| Table S4 | Relative positions of predicted WalR binding and transcriptional start sites (TSS). | 20-23 |
| Figure S5 | Purification of recombinant WalR. | 24 |
| Figure S6 | Competition EMSAs. | 25 |
| Figure S7 | WalR and VraR binding at the *ltaS* locus. | 26 |
| Figure S8 | Impact of WalR motif mutation on *tarF*, *tagG* and *dnaD.* | 27 |
| Figure S9 | The effect of *walR*, *hup*, and *hla* on *ori*:*ter* ratio. | 28 |
| Table S5 | Strains and plasmids used in this study | 29-32 |
| Table S6 | Oligonucleotides used in this study | 33-40 |
| References | | 41 |

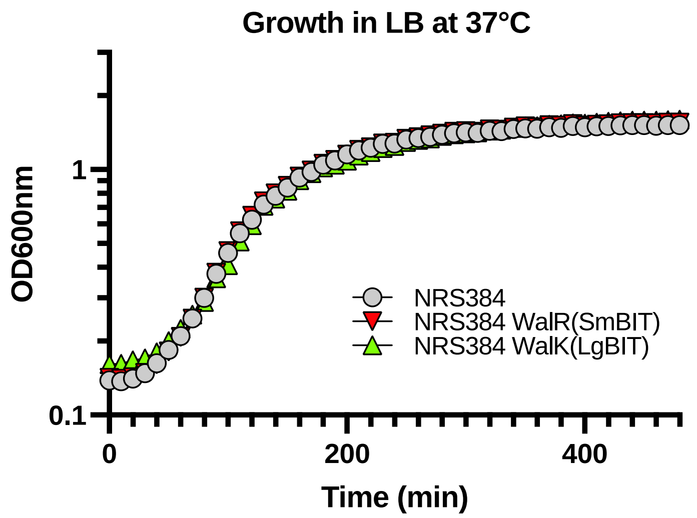

***Figure S1. Growth in LB of the chromosomally C-terminally tagged WalR-SmBIT or WalK-LgBIT in LB compared to the Wt.*** *No difference in growth profile was observed for the mutants.*

| Cellular function | Locus tag | Gene | Product | RNA-seq significance (FDR) | $Log_2FC$ |
|---|---|---|---|---|---|
| Amino acid biosynthesis and transport | SAUSA300_2006 | *ilvD* | Dihydroxy-Acid Dehydratase | 4.00E-07 | 2.84 |
| | SAUSA300_2007 | *ilvB* | Acetolactate Synthase Large Subunit | 1.00E-07 | 2.46 |
| | SAUSA300_2013 | *leuD* | Isopropylmalate Isomerase Small Subunit | 2.30E-06 | 2.06 |
| | SAUSA300_0185 | *argJ* | Bifunctional Ornithine Acetyltransferase/N-Acetylglutamate Synthase Protein | 3.30E-05 | 2.02 |
| | SAUSA300_2010 | *leuA* | 2-Isopropylmalate Synthase | 1.10E-06 | 2.01 |
| | SAUSA300_0186 | *argC* | N-Acetyl-Gamma-Glutamyl-Phosphate Reductase | 1.10E-06 | 1.95 |
| | SAUSA300_2012 | *leuC* | Isopropylmalate Isomerase Large Subunit | 4.00E-07 | 1.88 |
| | SAUSA300_0359 | *metC* | Trans-Sulfuration Enzyme Family Protein | 1.50E-06 | 1.87 |
| | SAUSA300_2009 | *ilvC* | Ketol-Acid Reductoisomerase | 9.00E-07 | 1.87 |
| | SAUSA300_2011 | *leuB* | 3-Isopropylmalate Dehydrogenase | 9.00E-07 | 1.68 |
| | SAUSA300_0184 | *argB* | Acetylglutamate Kinase | 2.10E-05 | 1.58 |
| | SAUSA300_0066 | *argR* | Arginine Repressor | 6.30E-05 | -1.75 |
| Carbohydrate uptake and metabolism | SAUSA300_0186 | *argC* | N-Acetyl-Gamma-Glutamyl-Phosphate Reductase | 1.10E-06 | 1.95 |
| | SAUSA300_0448 | *treP* | PTS System, Trehalose-Specific Iibc Component | 2.20E-05 | 1.72 |
| | SAUSA300_0315 | *nanA* | N-Acetylneuraminate Lyase | 1.10E-04 | 1.7 |
| | SAUSA300_0236 | *glcC* | PTS System, IIBC Components | 1.70E-06 | 1.64 |
| | SAUSA300_0333 | - | BglG Family Transcriptional Antiterminator | 2.20E-04 | 1.56 |
| | SAUSA300_1258 | *dmpI* | 4-Oxalocrotonate Tautomerase | 2.60E-04 | -1.56 |
| | SAUSA300_0791 | *gcvH* | Glycine Cleavage System Protein H | 1.20E-06 | -1.76 |
| | SAUSA300_0930 | *lplA1* | Lipoate-Protein Ligase A Family Protein | 5.00E-07 | -1.79 |
| | SAUSA300_0030 | *ugpQ* | Glycerophosphoryl Diester Phosphodiesterase | 3.00E-07 | -2 |
| Cell wall organisation | SAUSA300_2585 | *asp3* | Accessory Secretory Protein Asp3 | 5.70E-05 | 1.92 |
| | SAUSA300_0153 | *capB* | Capsular Polysaccharide Biosynthesis Protein Cap5B | 5.10E-05 | 1.6 |
| | SAUSA300_0156 | *capE* | Capsular Polysaccharide Biosynthesis Protein Cap5E | 5.20E-05 | 1.57 |
| | SAUSA300_0603 | - | Response To Antimicrobial Peptides | 1.80E-05 | -1.59 |
| | SAUSA300_0604 | - | Alpha/Beta Fold Family Hydrolase | 7.80E-06 | -1.61 |
| Cofactor and carrier biosynthesis | SAUSA300_0697 | *queC* | ExsB Protein | 8.40E-06 | 1.85 |
| DNA replication, recombination, and repair | SAUSA300_0367 | *ssb* | Single-Strand Binding Protein | 1.90E-06 | 1.94 |
| | SAUSA300_1142 | *dprA* | DNA Protecting Protein DprA | 2.10E-06 | 1.83 |
| | SAUSA300_1732 | *tnp2* | Putative Transposase | 3.10E-03 | -4.62 |
| Fatty acid biosynthesis | SAUSA300_0320 | *gehB* | Triacylglycerol Lipase | 4.00E-07 | 2.56 |

| | | | | | |
|---|---|---|---|---|---|
| | SAUSA300_1563 | - | Acetyl-CoA Carboxylase, Biotin Carboxylase | 3.20E-05 | 2.01 |
| | SAUSA300_0177 | - | Acyl-CoA/Acyl-Acp Dehydrogenase | 5.00E-07 | 1.96 |
| | SAUSA300_1125 | *acpP* | Acyl Carrier Protein | 9.90E-06 | -1.92 |
| Metal ion homeostasis | SAUSA300_0720 | *sstC* | Putative Iron Compound ABC Transporter ATP-Binding Protein | 9.00E-05 | 1.6 |
| | SAUSA300_0117 | *sirA* | Iron Compound ABC Transporter Iron Compound-Binding Protein SirA | 5.60E-05 | -1.51 |
| | SAUSA300_1874 | *ftnA* | Ferritins Family Protein | 2.20E-06 | -1.69 |
| | SAUSA300_0843 | *sufA* | Fe-S Cluster Carrier | 2.30E-04 | -1.7 |
| | SAUSA300_1373 | *fer* | Ferredoxin | 3.50E-05 | -2.14 |
| Nitrogen utilisation | SAUSA300_1566 | - | Allophanate Hydrolase | 2.10E-05 | 1.71 |
| Oxidative stress response | SAUSA300_2502 | *crtO* | Staphyloxanthin Biosynthesis | 6.80E-05 | -1.65 |
| | SAUSA300_1909 | - | Putative Thioredoxin | 7.00E-07 | -1.86 |
| | SAUSA300_1197 | *bsaA* | Glutathione Peroxidase | 3.50E-06 | -1.87 |
| | SAUSA300_2463 | *ddh* | D-Lactate Dehydrogenase | 1.70E-06 | -1.97 |
| | SAUSA300_1044 | *trxA* | Thioredoxin | 2.00E-06 | -2.2 |
| | SAUSA300_0789 | - | Putative Thioredoxin | 1.00E-07 | -2.3 |
| | SAUSA300_1690 | - | Putative Thioredoxin | 2.00E-07 | -2.35 |
| Phage associated | SAUSA300_1947 | - | Phi77 Orf031-Like Protein | 4.90E-03 | -1.71 |
| | SAUSA300_1944 | - | Phi77 Orf026-Like Protein Phage Transcriptional Activator | 8.20E-03 | -1.71 |
| Protein fate | SAUSA300_1656 | *uspA1* | Universal Stress Protein | 2.20E-06 | -1.64 |
| | SAUSA300_1984 | *mroQ* | Membrane-Embedded CaaX Protease | 2.10E-05 | -1.69 |
| | SAUSA300_1790 | *prsA* | Foldase Protein PrsA | 4.00E-07 | -1.8 |
| | SAUSA300_0752 | *clpP* | ATP-Dependent Clp Protease Proteolytic Subunit | 1.40E-06 | -1.89 |
| | SAUSA300_1295 | *cspA* | Csp Family Cold Shock Protein | 2.00E-07 | -3.19 |
| | SAUSA300_0067 | - | Universal Stress Protein | 2.00E-07 | -3.35 |
| Purine ribonucleotide biosynthesis | SAUSA300_0973 | *purM* | Phosphoribosylaminoimidazole Synthetase | 1.00E-07 | 2.65 |
| | SAUSA300_0971 | *purL* | Phosphoribosylformylglycinamidine Synthase II | 1.20E-06 | 2.48 |
| | SAUSA300_0970 | *purQ* | Phosphoribosylformylglycinamidine Synthase I | 3.10E-05 | 2.19 |
| | SAUSA300_0972 | *purF* | Amidophosphoribosyltransferase | 4.00E-07 | 2.15 |
| | SAUSA300_0974 | *purN* | Phosphoribosylglycinamide Formyltransferase | 5.00E-07 | 2.01 |
| | SAUSA300_0975 | *purH* | Bifunctional Phosphoribosylaminoimidazolecarboxamide Formyltransferase/Imp Cyclohydrolase | 8.00E-07 | 1.61 |
| | SAUSA300_2183 | *adk* | Adenylate Kinase | 4.30E-05 | 1.57 |

| | | | | | |
|---|---|---|---|---|---|
| Respiration | SAUSA300_0960 | *qoxD* | Quinol Oxidase, Subunit Iv | 9.90E-05 | 1.7 |
| Ribosome and Protein synthesis | SAUSA300_2186 | *rpmD* | 50S Ribosomal Protein L30 | 1.70E-06 | 2.05 |
| | SAUSA300_0531 | *rpsG* | 30S Ribosomal Protein S7 | 2.70E-06 | 1.79 |
| | SAUSA300_2179 | *rpsK* | 30S Ribosomal Protein S11 | 3.40E-05 | 1.75 |
| | SAUSA300_2187 | *rpsE* | 30S Ribosomal Protein S5 | 7.40E-06 | 1.7 |
| | SAUSA300_2180 | *rpsM* | 30S Ribosomal Protein S13 | 2.10E-05 | 1.52 |
| | SAUSA300_2188 | *rplR* | 50S Ribosomal Protein L18 | 2.10E-05 | 1.51 |
| | SAUSA300_2648 | *rpmH* | 50S Ribosomal Protein L34 | 1.10E-03 | -1.56 |
| | SAUSA300_0991 | *def* | Peptide Deformylase | 1.50E-06 | -1.81 |
| | SAUSA300_1233 | *rpmG2* | 50S Ribosomal Protein L33 | 1.00E-04 | -1.92 |
| | SAUSA300_1545 | *rpsT* | 30S Ribosomal Protein S20 | 3.10E-05 | -2.13 |
| | SAUSA300_0053 | *speG* | Spermidine N(1)-Acetyltransferase | 6.50E-06 | -2.19 |
| | SAUSA300_0736 | *saHPF* | Hibernation-Promoting Factor | 4.00E-07 | -2.26 |
| | SAUSA300_1511 | *rpmG* | 50S Ribosomal Protein L33 | 6.30E-05 | -2.33 |
| | SAUSA300_1027 | *rpmF* | 50S Ribosomal Protein L32 | 7.40E-06 | -2.38 |
| | SAUSA300_1234 | *rpsN2* | 30S Ribosomal Protein S14 | 4.70E-06 | -2.44 |
| | SAUSA300_1535 | *rpsU* | 30S Ribosomal Protein S21 | 7.90E-06 | -3.11 |
| Transcriptional regulation | SAUSA300_0691 | *saeR* | DNA-Binding Response Regulator SaeR | 5.00E-07 | 2.28 |
| | SAUSA300_0690 | *saeS* | Sensor Histidine Kinase SaeS | 1.00E-07 | 2.2 |
| | SAUSA300_0928 | *comK1* | Competence Transcription Factor | 9.90E-06 | 1.86 |
| | SAUSA300_0187 | *argD* | Ornithine Aminotransferase | 2.10E-06 | 1.85 |
| | SAUSA300_1992 | *agrA* | Accessory Gene Regulator Protein A | 2.40E-06 | 1.77 |
| | SAUSA300_1991 | *agrC* | Accessory Gene Regulator Protein C | 1.40E-06 | 1.57 |
| | SAUSA300_2337 | *nreC* | DNA-Binding Response Regulator Nrec | 3.00E-03 | -1.53 |
| | SAUSA300_1708 | *rot* | Accessory Regulator Rot | 3.10E-06 | -1.63 |
| | SAUSA300_0114 | *sarS* | Accessory Regulator | 1.80E-06 | -1.76 |
| | SAUSA300_2218 | *sarV* | Staphylococcal Accessory Regulator V | 2.40E-05 | -1.89 |
| | SAUSA300_0954 | - | MarR Family Transcriptional Regulator | 1.10E-06 | -2.3 |
| | SAUSA300_2599 | *icaR* | Intercellular Adhesion Operon Transcription Regulator | 5.00E-07 | -2.86 |
| | SAUSA300_2437 | *sarT* | Accessory Regulator T | 9.40E-06 | -3.14 |
| Transport | SAUSA300_0176 | *ssuC* | ABC Transporter Permease | 4.70E-06 | 1.95 |
| | SAUSA300_0436 | *gmpB* | ABC Transporter Permease | 8.00E-07 | 1.94 |
| | SAUSA300_0202 | - | Peptide ABC Transporter Permease | 4.40E-05 | 1.8 |
| | SAUSA300_0203 | - | Nickel-Peptide/Transporter Substrate-Binding Protein | 1.10E-06 | 1.74 |
| | SAUSA300_0201 | - | Peptide ABC Transporter Permease | 1.20E-06 | 1.68 |

| | | | | | |
|---|---|---|---|---|---|
| | SAUSA300_0143 | *phnE2* | Phosphonate ABC Transporter Permease | 2.00E-05 | 1.6 |
| | SAUSA300_0200 | - | Peptide ABC Transporter ATP-Binding Protein | 1.10E-06 | 1.59 |
| | SAUSA300_0208 | *malK* | Putative Maltose ABC Transporter ATP-Binding Protein | 5.00E-05 | 1.52 |
| | SAUSA300_2307 | *hrtB* | ABC Transporter Permease | 2.00E-05 | -1.61 |
| Virulence | SAUSA300_1755 | *splD* | Serine Protease SplD | 2.00E-07 | 5.79 |
| | SAUSA300_1757 | *splB* | Serine Protease SplB | 0.00E+00 | 4.45 |
| | SAUSA300_1758 | *splA* | Serine Protease SplA | 0.00E+00 | 4.15 |
| | SAUSA300_1754 | *splE* | Serine Protease SplE | 8.00E-07 | 3.83 |
| | SAUSA300_1756 | *splC* | Serine Protease SplC | 2.00E-07 | 3.44 |
| | SAUSA300_1753 | *splF* | Serine Protease SplF | 3.00E-07 | 3.05 |
| | SAUSA300_2600 | *icaA* | N-Glycosyltransferase | 3.40E-06 | 2.66 |
| | SAUSA300_1382 | *lukS-PV* | Panton-Valentine Leukocidin, LukS-PV | 1.00E-07 | 2.59 |
| | SAUSA300_0951 | *sspA* | V8 Protease | 1.00E-07 | 2.55 |
| | SAUSA300_1975 | *lukH* | Aerolysin/Leukocidin Family Protein | 8.50E-06 | 2.37 |
| | SAUSA300_2364 | *sbi* | IgG-Binding Protein Sbi | 1.00E-07 | 2.15 |
| | SAUSA300_1920 | *chp* | Chemotaxis-Inhibiting Protein Chips | 2.90E-06 | 2 |
| | SAUSA300_1918 | *hlb-1* | Truncated Beta-Hemolysin | 3.80E-05 | 1.93 |
| | SAUSA300_2586 | *asp2* | Accessory Sec System Protein | 1.10E-06 | 1.91 |
| | SAUSA300_0950 | *sspB* | Cysteine Protease | 1.20E-06 | 1.88 |
| | SAUSA300_0285 | *esxB* | Type VII Secretion System Extracellular Protein B | 2.80E-06 | 1.87 |
| | SAUSA300_2367 | *hlgB* | Gamma-Hemolysin Component B | 1.10E-06 | 1.83 |
| | SAUSA300_1381 | - | Panton-Valentine Leukocidin, LukF-PV | 3.00E-07 | 1.82 |
| | SAUSA300_1058 | *hla* | Alpha-Hemolysin | 1.20E-06 | 1.78 |
| | SAUSA300_2184 | *secY* | Preprotein Translocase Subunit SecY | 2.50E-06 | 1.76 |
| | SAUSA300_1974 | *lukG* | Leukocidin/Hemolysin Toxin Family Protein | 5.50E-05 | 1.71 |
| | SAUSA300_0949 | *sspC* | Cysteine Protease | 1.80E-05 | 1.71 |
| | SAUSA300_2587 | *asp1* | Accessory Secretory Protein Asp1 | 5.30E-05 | 1.69 |
| | SAUSA300_2601 | *icaB* | Intercellular Adhesion Protein B | 1.40E-05 | 1.54 |
| | SAUSA300_2572 | *aur* | Zinc Metalloproteinase Aureolysin | 1.10E-06 | 1.5 |
| | SAUSA300_0899 | *trfA* | Adaptor Protein | 1.10E-06 | -1.64 |
| | SAUSA300_1784 | *traP* | Signal Transduction Protein TraP | 4.90E-06 | -1.65 |
| | SAUSA300_0401 | *ssl7* | Superantigen-Like Protein 7 | 9.90E-06 | -1.65 |
| | SAUSA300_2262 | *spdB* | Abi Domain-Containing Protein | 1.30E-05 | -1.69 |
| | SAUSA300_0762 | *secG* | Preprotein Translocase Subunit SecG | 4.40E-04 | -1.74 |
| | SAUSA300_1594 | *yajC* | Preprotein Translocase Subunit YajC | 4.60E-05 | -1.94 |
| | SAUSA300_0289 | *esaG* | Component Of the Type VII Secretion System | 1.00E-05 | -2.02 |
| | SAUSA300_0395 | *ssl1* | Superantigen-Like Protein | 2.30E-05 | -2.07 |

| | | | | | |
|---|---|---|---|---|---|
| | SAUSA300_1988 | *hld* | Delta-Hemolysin | 6.50E-05 | -2.09 |
| | SAUSA300_2573 | *isaB* | Immunodominant Antigen B | 8.00E-07 | -2.28 |
| | SAUSA300_0399 | *ssl5* | Superantigen-Like Protein 5 | 7.80E-06 | -2.54 |
| | SAUSA300_0654 | *sarX* | Hypothetical Protein | 1.30E-06 | -3.03 |
| Unknown | SAUSA300_0692 | *saeQ* | Regulator of SaeS | 2.00E-07 | 3.12 |
| | SAUSA300_0175 | *ssuA* | Putative Lipoprotein | 1.20E-06 | 2.3 |
| | SAUSA300_1877 | - | Hypothetical Protein. Transmembrane | 5.00E-07 | 2.06 |
| | SAUSA300_0174 | *ssuB* | Hypothetical Protein, Putative ABC Transporter | 3.60E-06 | 2 |
| | SAUSA300_0326 | - | Hypothetical Protein, Protein-Adp-Ribose Hydrolase | 9.90E-06 | 1.79 |
| | SAUSA300_1565 | - | Hypothetical Protein, Ahs2 Domain-Containing | 5.10E-05 | 1.72 |
| | SAUSA300_1734 | *rppH* | Hypothetical Protein, Nudix Hydrolase Domain Containing | 2.00E-05 | 1.52 |
| | SAUSA300_2448 | - | Hypothetical Protein, Putative Transmembrane | 3.40E-06 | 1.5 |
| | SAUSA300_0427 | *mpsC* | Hypothetical Protein, MpsC Domain Containing | 2.20E-06 | -1.52 |
| | SAUSA300_2352 | - | Addiction Module Antitoxin | 1.00E-06 | -1.53 |
| | SAUSA300_2632 | - | Hypothetical Protein, Putative Transmembrane | 2.00E-05 | -1.53 |
| | SAUSA300_2401 | - | Addiction Module Antitoxin | 8.10E-05 | -1.54 |
| | SAUSA300_0385 | - | Hypothetical Protein, Putative Transmembrane | 1.80E-05 | -1.54 |
| | SAUSA300_2132 | - | Hypothetical Protein Of Unknown Function | 1.60E-04 | -1.54 |
| | SAUSA300_2528 | - | Hypothetical Protein, Pepsy Domain-Containing | 3.70E-05 | -1.54 |
| | SAUSA300_2093 | - | Hypothetical Protein, Protein-Disulfide Reductase Activity | 8.80E-06 | -1.55 |
| | SAUSA300_1054 | - | Hypothetical Protein, Putative Transmembrane | 7.80E-05 | -1.57 |
| | SAUSA300_2624 | - | Hypothetical Protein, Putative Transmembrane | 2.50E-06 | -1.57 |
| | SAUSA300_1977 | - | Hypothetical Protein, Putative Methyltransferase | 1.50E-04 | -1.58 |
| | SAUSA300_0609 | - | Phage Integrase Family Protein | 1.40E-05 | -1.61 |
| | SAUSA300_1685 | - | Hypothetical Protein, Putative Transmembrane | 1.10E-04 | -1.61 |
| | SAUSA300_0940 | - | Hypothetical Protein, Putative Transmembrane | 1.40E-05 | -1.68 |
| | SAUSA300_0304 | - | Hypothetical Protein Of Unknown Function | 6.70E-06 | -1.69 |
| | SAUSA300_1803 | - | Hypothetical Protein, Putative Transmembrane | 5.00E-07 | -1.7 |
| | SAUSA300_2402 | - | Hypothetical Protein, Putative Antitoxin | 1.40E-05 | -1.71 |
| | SAUSA300_0094 | - | Hypothetical Protein Of Unknown Function | 1.30E-04 | -1.72 |
| | SAUSA300_1495 | - | Hypothetical Protein, Rhodanese-Like Domain Containing | 5.50E-06 | -1.74 |
| | SAUSA300_1213 | - | Hypothetical Protein Of Unknown Function | 4.20E-04 | -1.74 |
| | SAUSA300_2334 | - | Hypothetical Protein, Peptidase Activity | 1.80E-05 | -1.75 |
| | SAUSA300_1606 | - | Hypothetical Protein Of Unknown Function | 7.50E-06 | -1.75 |
| | SAUSA300_2252 | - | Hypothetical Protein, Putative Transmembrane | 4.00E-06 | -1.79 |
| | SAUSA300_1856 | - | Hypothetical Protein, Putative PfpI Endopeptidase | 1.70E-06 | -1.79 |
| | SAUSA300_1099 | - | Hypothetical Protein Of Unknown Function | 3.90E-04 | -1.8 |

| | | | | |
|---|---|---|---|---|
| SAUSA300_0831 | - | Hypothetical Protein Of Unknown Function | 2.70E-05 | -1.82 |
| SAUSA300_2080 | - | Hypothetical Protein Of Unknown Function | 2.50E-06 | -1.84 |
| SAUSA300_2418 | - | Hypothetical Protein, Putative Peroxiredoxin | 3.60E-06 | -1.87 |
| SAUSA300_1802 | - | Hypothetical Protein Of Unknown Function | 1.30E-05 | -1.89 |
| SAUSA300_0668 | - | Hypothetical Protein Of Unknown Function | 1.10E-06 | -1.9 |
| SAUSA300_1057 | - | Hypothetical Protein Of Unknown Function | 8.10E-06 | -1.9 |
| SAUSA300_2368 | *bioX* | Hypothetical Protein, Putative Transmembrane | 1.20E-04 | -1.93 |
| SAUSA300_2637 | - | Hypothetical Protein, Putative Phage Tail Protein | 6.30E-06 | -1.93 |
| SAUSA300_1272 | - | Hypothetical Protein, Swim-Type Domain-Containing | 1.80E-05 | -1.97 |
| SAUSA300_1212 | - | Hypothetical Protein, Ntox50 Domain-Containing | 5.40E-05 | -2 |
| SAUSA300_2354 | *dsbA* | Putative Lipoprotein | 1.10E-06 | -2.02 |
| SAUSA300_0342 | | Hypothetical Protein Of Unknown Function | 1.30E-05 | -2.04 |
| SAUSA300_0292 | - | Hypothetical Protein, Putative Transmembrane | 2.40E-05 | -2.05 |
| SAUSA300_2206 | - | Hypothetical Protein, Putative Transmembrane | 4.00E-07 | -2.05 |
| SAUSA300_0942 | - | Hypothetical Protein, Pepsy Domain-Containing | 3.00E-05 | -2.05 |
| SAUSA300_2527 | - | Hypothetical Protein Of Unknown Function | 8.60E-06 | -2.06 |
| SAUSA300_1041 | - | Hypothetical Protein, Putative Transmembrane | 8.80E-06 | -2.07 |
| SAUSA300_0172 | - | Hypothetical Protein Of Unknown Function | 4.40E-06 | -2.07 |
| SAUSA300_2543 | - | Hypothetical Protein, Signal Transduction Protein Trap | 1.80E-05 | -2.14 |
| SAUSA300_1692 | - | Hypothetical Protein, Putative Transmembrane | 2.80E-05 | -2.15 |
| SAUSA300_1230 | - | Hypothetical Protein Of Unknown Function | 1.10E-05 | -2.17 |
| SAUSA300_2053 | *ywpF* | T6SS Effector-Like Protein Of Unknown Function | 4.00E-06 | -2.18 |
| SAUSA300_1221 | - | Hypothetical Protein Of Unknown Function | 3.10E-04 | -2.18 |
| SAUSA300_0884 | - | Hypothetical Protein Of Unknown Function | 9.40E-06 | -2.23 |
| SAUSA300_1004 | - | Hypothetical Protein Of Unknown Function | 8.40E-06 | -2.27 |
| SAUSA300_0781 | - | Hypothetical Protein Of Unknown Function | 2.30E-05 | -2.3 |
| SAUSA300_2562 | - | Hypothetical Protein Of Unknown Function | 6.60E-05 | -2.32 |
| SAUSA300_0565 | - | Hypothetical Protein, Putative Transmembrane | 1.70E-06 | -2.32 |
| SAUSA300_1326 | *rnhA* | Putative Cell Wall Enzyme EbsB | 8.10E-06 | -2.34 |
| SAUSA300_0929 | - | Hypothetical Protein Of Unknown Function | 1.30E-05 | -2.34 |
| SAUSA300_0372 | - | Hypothetical Protein, Pepsy Domain | 1.00E-07 | -2.36 |
| SAUSA300_1240 | - | Hypothetical Protein, Putative Transmembrane | 1.10E-05 | -2.37 |
| SAUSA300_0050 | - | Hypothetical Protein, Putative Transmembrane | 1.40E-05 | -2.41 |
| SAUSA300_1107 | - | Hypothetical Protein, Putative Transmembrane | 2.50E-05 | -2.41 |
| SAUSA300_1277 | - | Hypothetical Protein Of Unknown Function | 1.10E-05 | -2.46 |
| SAUSA300_1484 | - | Hypothetical Protein, Putative Transmembrane | 3.50E-05 | -2.49 |
| SAUSA300_1478 | - | Putative Lipoprotein | 8.00E-07 | -2.6 |
| SAUSA300_1478 | - | Putative Lipoprotein | 8.00E-07 | -2.6 |

| | | | | |
|---|---|---|---|---|
| SAUSA300_2481 | - | Hypothetical Protein, YozE Sam Like Domain Containing | 2.00E-05 | -2.6 |
| SAUSA300_0373 | - | Hypothetical Protein, Hth Cro/C1-Type Domain-Containing Protein | 9.20E-06 | -2.66 |
| SAUSA300_2403 | - | Putative Lipoprotein | 5.00E-07 | -2.67 |
| SAUSA300_2403 | - | Putative Lipoprotein | 5.00E-07 | -2.67 |
| SAUSA300_2246 | - | Hypothetical Protein, Bph_3 Domain | 1.00E-07 | -2.67 |
| SAUSA300_2544 | - | Hypothetical Protein, YozE Sam Like Domain Containing | 9.90E-06 | -2.73 |
| SAUSA300_1432 | - | PhiSLT Orf78-Like Protein | 1.70E-05 | -2.74 |
| SAUSA300_2041 | - | Hypothetical Protein Of Unknown Function | 1.90E-05 | -2.74 |
| SAUSA300_1208 | - | Hypothetical Protein Of Unknown Function | 9.30E-06 | -2.81 |
| SAUSA300_0266 | - | Hypothetical Protein Of Unknown Function | 6.70E-06 | -2.86 |
| SAUSA300_1581 | - | Hypothetical Protein Of Unknown Function | 4.00E-07 | -2.89 |
| SAUSA300_2493 | *cwrA* | Hypothetical Protein, Cell Wall Stress Stimulon | 5.80E-06 | -2.92 |
| SAUSA300_0937 | - | Hypothetical Protein, Putative Transmembrane | 9.90E-06 | -2.93 |
| SAUSA300_1493 | - | Hypothetical Protein Of Unknown Function | 6.40E-06 | -2.95 |
| SAUSA300_1008 | - | Hypothetical Protein Of Unknown Function | 7.80E-06 | -3.03 |
| SAUSA300_1795 | - | Hypothetical Protein Of Unknown Function | 1.00E-07 | -3.04 |
| SAUSA300_2328 | - | Hypothetical Protein Of Unknown Function | 2.00E-07 | -3.1 |
| SAUSA300_2460 | - | Acetyltransferase Family Protein | 9.00E-07 | -3.17 |
| SAUSA300_2460 | - | Acetyltransferase Family Protein | 9.00E-07 | -3.17 |
| SAUSA300_1180 | - | Hypothetical Protein Of Unknown Function | 1.00E-06 | -3.78 |
| SAUSA300_1215 | - | Hypothetical Protein Of Unknown Function | 1.20E-06 | -3.83 |
| SAUSA300_0816 | - | CsbB-Like Superfamily Protein | 4.00E-07 | -4 |
| SAUSA300_1904 | - | Hypothetical Protein, Putative Transmembrane | 5.00E-07 | -4.36 |

***Table S1. The indirect WalR regulon defined by RNA-seq.*** *Genes undergoing a ≥1.5 log2FC (FDR ≤0.05) change in gene expression upon WalK activation (WalKT389A mutant vs wild type) are grouped by function. FDR = P value adjusted for false discovery rate, Log2FC = fold-change.*

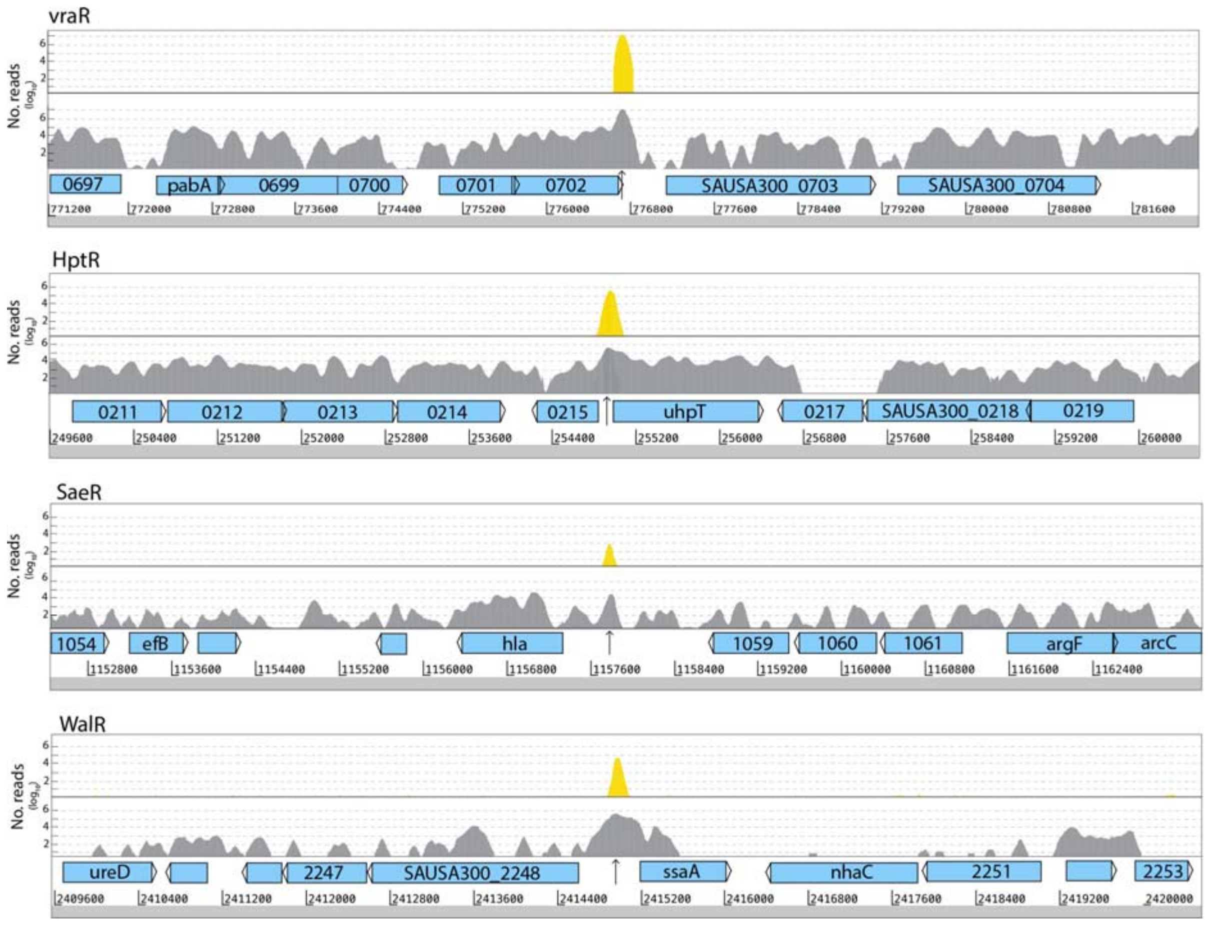

***Figure S2 S. aureus ChIP-seq reveals binding sites of response regulators.*** *Artemis userplots showing examples of the impact of non-target in silico read-subtraction (grey reads – below) for the identification of target-specific response regulator DNA-binding regions (yellow reads – above).*

**Table S2:** Putative response-regulator binding sites identified by ChIP-seq

| Locus ID | Gene name | Product | Function | Impact on gene regulation | Nebraska Tpn insert | ChIP-seq bind coords | Peak prediction score | Motif detected | Motif coords |
|---|---|---|---|---|---|---|---|---|---|
| **WalR** | | | | | | | | | |
| SAUSA300_0136 | *sasD* | Cell-wall associated protein | Contains LPXAG motif – likely covalently anchored to peptidoglycan | Negative (5.8-log2 decrease, 56-fold) | Yes | 154986..155175 | 4.24929 | Yes | 155136..155152 |
| SAUSA300_0438 | *sle1* | N-acetylmuramoyl-L-alanine amidase (autolysin) | cleaves amide bond between N-acetylmuramoyl and L-amino acids in peptidoglycan | Positive (3.5-log2 increase, 12-fold) | Yes | 491543..491550 | 0.97290 | Yes | 491559..491575 |
| SAUSA300_0524 | *rplJ* | 50S ribosomal protein L10 | binds the two ribosomal protein L7/L12 dimers and anchors them to the large ribosomal subunit | No regulation | No | 583428..583441 | 0.62689 | No | |
| SAUSA300_0602 | | Hypothetical protein | unknown | Negative (3.3-log2 decrease, 10-fold) | Yes | 675677..675892 | 5.77914 | Yes | 675820..675847 |
| SAUSA300_0653 | | AraC family transcriptional regulator | Transcriptional regulation | Negative (1.2-log2 decrease, 3-fold) | Yes | 728668..728678 | 0.64084 | No | |
| SAUSA300_0681 | | hypothetical protein | | Positive (2.3-log2 decrease in down mutant | No | 755739..755812 | 2.21698 | No | |
| SAUSA300_0703 | *ltaS* | Glycerol phosphate lipoteichoic acid synthase | Synthesis of polyglycerol-phosphate lipoteichoic acid (LTA) from phosphatidylglycerol | Negative (0.7-log decrease, 1.6-fold) | No | 776863..776913 | 2.43150 | Yes | 776866..776882 |
| SAUSA300_0860 | *rocD* | ornithine--oxo-acid transaminase | | Positive (1.2-log2 increase, 2-fold) | Yes | 937633..937638 | 0.462098 | Yes | 937577..937593 |
| SAUSA300_1081 | | Hypothetical protein | | No regulation | Yes | 1182912..1182918 | 0.610952 | No | |
| SAUSA300_1089 | *lspA* | lipoprotein signal peptidase | lipoprotein signal peptidase; integral membrane protein that removes signal peptides from prolipoproteins during lipoprotein biosynthesis | No regulation | Yes | 1191626..1191643 | 0.917025 | Yes | 1191606..1191622 |
| SAUSA300_1188 | *mutS* | DNA mismatch repair protein MutS | This protein performs the mismatch recognition step during the DNA repair process | No regulation | Yes | 1306553..1306561 | 0.539114 | No | |
| SAUSA300_1203 | | Hypothetical protein | | | Yes | 1325832..1325849 | 1.02376 | No | |
| SAUSA300_1304 | | Hypothetical protein | | No regulation | Yes | 1435403..1435418 | 1.04382 | No | |
| SAUSA300_1344 | *dnaD* | putative DNA replication protein DnaD | | No regulation | No | 1509472..1509481 | 0.554517 | Yes | 1509483..1509499 |
| SAUSA300_1461 | | Hypothetical protein | | No regulation | Yes | 1615853..1615880 | 1.549569 | No | |
| SAUSA300_1841 | *rrsC* | 16S rRNA | | | | 2002908..2002944 | 1.675390 | Yes | 2003159..2003174 |

| Locus ID | Gene name | Product | Function | Impact on gene regulation | Nebraska Tpn insert | ChIP-seq bind coords | Peak prediction score | Motif detected | Motif coords |
|---|---|---|---|---|---|---|---|---|---|
| SAUSA300_1890 | | staphopain A | | Positive (0.85-log2 increase, 1.8-fold) | Yes | 2055015..2055025 | 0.740715 | No | |
| SAUSA300_2124 | *rrsE* | 16S rRNA | | | | 2297762..2297799 | 2.261965 | No | |
| SAUSA300_2209 | | Hypothetical protein | | Negative (0.84-log2 decrease, 1.8-fold) | Yes | 2375987..2375993 | 0.553028 | No | |
| SAUSA300_2249 | *ssaA_1* | N-acetylmuramoyl-L-alanine amidase (autolysin) | cleaves amide bond between N-acetylmuramoyl and L-amino acids in peptidoglycan | Positive (1.3-log2 increase, 2.5-fold) | Yes | 2414865..2415012 | 4.061749 | Yes (2x) | 2414918..2414933 2415046..2415061 |
| SAUSA300_2506 | *isaA* | Lytic transglycosylase | cleaves the β-1,4 glycosidic bond between N-acetylmuramic acid and N-acetylglucosamine residues of peptidoglycan | Positive (1.1-log2 increase, 2.2-fold) | No | 2712808..2712836 (peak-1) | 1.933075 | Yes | 2712897..2712912 |
| SAUSA300_2506 | *isaA* | Lytic transglycosylase | | Positive (1.1-log2 increase, 2.2-fold) | No | 2712870..2713227 (peak-2) | 4.523005 | Yes | 2713089..2713104 |
| **VraR** | | | | | | | | | |
| SAUSA300_0703 | *ltaS* | Glycerol phosphate lipoteichoic acid synthase | Synthesis of polyglycerol-phosphate lipoteichoic acid (LTA) from phosphatidylglycerol | | | 776616..776813 | 5.536103 | Yes | |
| SAUSA300_1357 | *aroC* | chorismate synthase | catalyzes the formation of chorismate from 5-O-(1-carboxyvinyl)-3-phosphoshikimate in aromatic amino acid biosynthesis | | | 1525199..1525211 | 0.8984054 | | |
| SAUSA300_2016 | *rrlD* | 23S ribosomal RNA | | | | | | | |
| **HptR** | | | | | | 2179043..2179049 | 4.352250 | | |
| SAUSA300_0151 | *adhE* | bifunctional acetaldehyde-CoA/alcohol dehydrogenase | | | | | | | |
| SAUSA300_0216 | *uhpT* | Sugar phosphate antiporter | cytoplasmic membrane protein that functions as a monomer; catalyzes the active transport of sugar-phosphates such as glucose-6-phosphate with the obligatory exchange of inorganic phosphate or organophosphate" | | | 254804..254999 | 4.867318 | Yes | |
| SAUSA300_0303 | | Hypothetical protein | | | | 352882..352918 | 1.4709476 | | |
| SAUSA300_0445 | *gltB* | glutamate synthase, large subunit | an essential enzyme in the nonmevalonate pathway of isopentenyl diphosphate and dimethylallyl diphosphate biosynthesis | | | 497269..497311 | 2.7078340 | | |

| Locus ID | Gene name | Product | Function | Impact on gene regulation | Nebraska Tpn insert | ChIP-seq bind coords | Peak prediction score | Motif detected | Motif coords |
|---|---|---|---|---|---|---|---|---|---|
| SAUSA300_0769 | | Hypothetical protein | | | | 857300..857341 | 0.927730 | | |
| SAUSA300_0883 | | Putative surface protein | | | | 969867..969932 | 1.503311 | | |
| SAUSA300_1052 | | fibrinogen-binding protein | | | | 1150372..1150493 | 2.338896 | | |
| SAUSA300_1058 | | alpha-hemolysin | | | | 1157882..1157919 | 1.312439 | | |
| SAUSA300_1463 | | Hypothetical protein | | | | 1617905..1617937 | 0.817594 | | |
| SAUSA300_1654 | | proline dipeptidase | | | | 1818897..1818945 | 1.959034 | | |
| SAUSA300_2012 | *leuC* | isopropylmalate isomerase large subunit | dehydratase component, catalyzes the isomerization between 2-isopropylmalate and 3-isopropylmalate | | | 2172237..2172454 | 5.527080 | | |
| SAUSA300_2504 | | acyltransferase | | | | 2710979..2711032 | 2.532463 | | |
| SAUSA300_2538 | | amino acid permease family protein | | | | 2738847..2738937 (peak-1) | 4.517702 | | |
| SAUSA300_2538 | | amino acid permease family protein | | | | 2739312..2739437 (peak-2) | 5.807741 | | |
| **SaeR** | | | | | | | | | |
| SAUSA300_0220 | *pflB* | formate acetyltransferase | | | | 260249..260326 | 2.245046 | | |
| SAUSA300_0472 | *ipk* | 4-diphosphocytidyl-2-C-methyl-D-erythritol kinase | An essential enzyme in the nonmevalonate pathway of isopentenyl diphosphate and dimethylallyl diphosphate biosynthesis | | | 530663..530696 | 1.313321 | | |
| SAUSA300_0892 | *oppA* | oligopeptide ABC transporter oligopeptide-binding protein | | | | 979938..980000 | 2.499846 | | |
| SAUSA300_0919 | *murE* | UDP-N-acetylmuramoylalanyl-D-glutamate--L-lysine ligase | involved in cell wall formation; peptidoglycan synthesis; cytoplasmic enzyme; catalyzes the addition of lysine to UDP-N-acetylmuramoyl-L-alanyl-D-glutamate forming UDP-N-acetylmuramoyl-L-alanyl-D-glutamyl-L-lysine | | | 1008357..1008398 | 1.681862 | | |
| SAUSA300_1058 | *hla* | Alpha-hemolysin | | | | 1157746..1157818 | 2.372307 | Yes | |
| SAUSA300_1648 | | putative NADP-dependent malic enzyme | | | | 1811157..1811173 | 1.025598 | | |
| SAUSA300_2144 | | Hypothetical protein | | | | 2320758..2320829 | 2.399629 | | |
| SAUSA300_2158 | | Hypothetical protein | | | | 2336211..2336232 | 0.992981 | | |
| SAUSA300_2423 | | Hypothetical protein | | | | 2609040..2609080 (peak-1) | 2.380980 | | |
| SAUSA300_2423 | | Hypothetical protein | | | | 2609182..2609231 (peak-2) | 2.487435 | | |

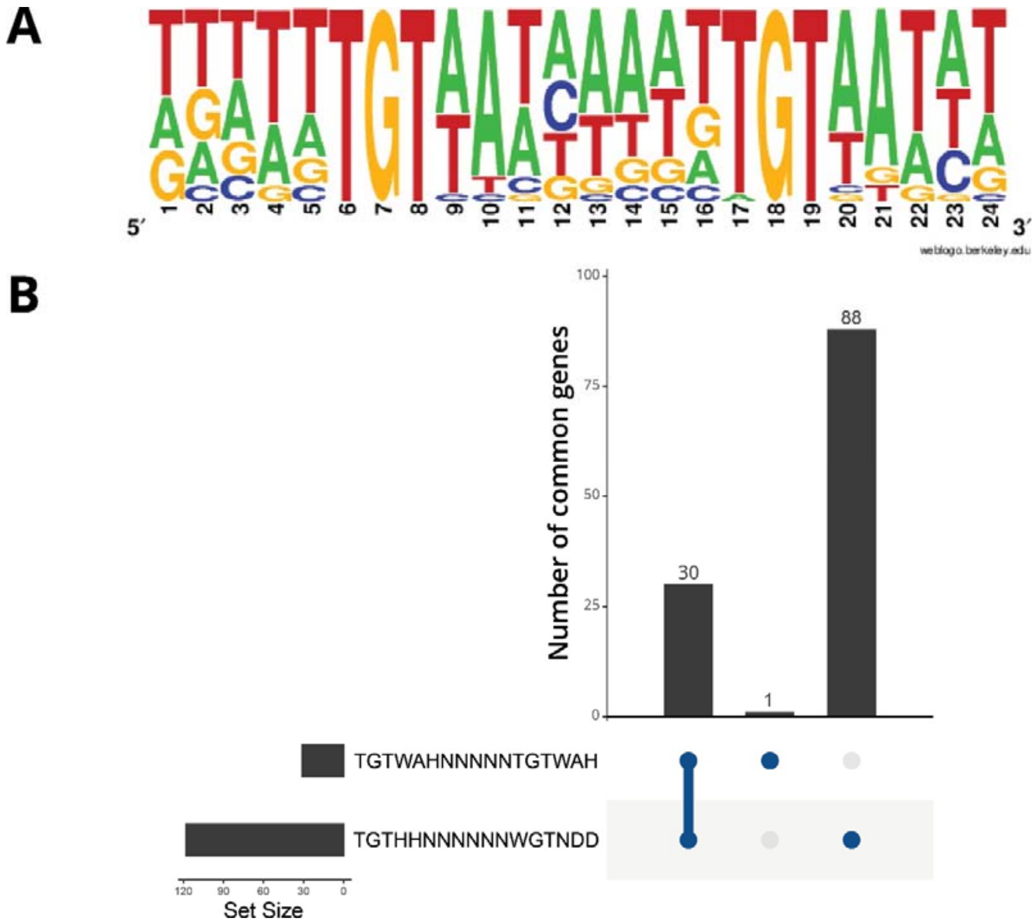

***Figure S3. Generation of an S. aureus WalR binding motif.*** *A) A combined frequency plot of experimentally validated WalR binding motifs and those detected through ChIP-seq analysis.* B) Upset plot comparison of overlap between intergenic WalR binding sites identified using a previously published motif (TGTWAHN5TGTWAH) and that defined in A) (TGTHHNNNNNNWGTNDD).

| Cellular function | Locus tag | Gene | Product | RNAseq significance (FDR) | $Log_2FC$ |
|---|---|---|---|---|---|
| Amino acid biosynthesis and transport | SAUSA300_0012 | *metX* | Putative Homoserine O-Acetyltransferase | 9.2E-06 | 1.60 |
| | SAUSA300_1808 | *glnP* | Amino Acid ABC Transporter Permease/Substrate-Binding Protein | 1.2E-05 | 1.38 |
| | SAUSA300_0860 | *rocD* | Ornithine-Oxo-Acid Transaminase | 1.6E-04 | 1.21 |
| | SAUSA300_0451 | - | Acetyltransferase | 2.2E-02 | 0.39 |
| | SAUSA300_1711 | *putA* | Proline Dehydrogenase | 6.7E-01 | -0.10 |
| Autolysis | SAUSA300_2253 | - | Secretory Antigen Precursor SsaA_3 | 6.0E-07 | 3.90 |
| | SAUSA300_0438 | *sle1* | Cell Wall Amidase | 6.0E-07 | 3.50 |
| | SAUSA300_2503 | - | Secretory Antigen Precursor SsaA_2 | 3.2E-06 | 3.09 |
| | SAUSA300_0955 | *atl* | Autolysin | 6.0E-07 | 2.85 |
| | SAUSA300_0651 | - | CHAP Domain-Containing Protein | 2.7E-06 | 1.68 |
| | SAUSA300_0277 | - | CHAP Domain-ContainingProtein, Predicted Autolysin | 4.7E-06 | 1.48 |
| | SAUSA300_2482 | - | CHAP Domain-Containing Protein, Predicted Autolysin | 2.7E-04 | 1.34 |
| | SAUSA300_2249 | *ssaA* | Secretory Antigen Precursor SsaA | 3.0E-05 | 1.31 |
| | SAUSA300_2506 | *isaA* | Lytic Transglycosylase | 9.3E-06 | 1.09 |
| | SAUSA300_0270 | *lytM* | Peptidoglycan Hydrolase | 5.9E-04 | 0.73 |
| | SAUSA300_0739 | - | LysM Domain-Containing Protein, Predicted Autolysin | 7.0E-03 | 0.61 |
| | SAUSA300_2051 | *sceD* | Lytic Transglycosylase | 5.5E-01 | -0.18 |
| Carbohydrate uptake and metabolism | SAUSA300_0216 | *hpt* | Glucose-6-Phosphate Antiporter | 9.6E-06 | 3.26 |
| | SAUSA300_2155 | *lacA* | Galactose-6-Phosphate Isomerase Subunit LacA | 3.0E-03 | 2.20 |
| | SAUSA300_1191 | *glpF* | Glycerol Uptake Facilitator | 2.9E-05 | 1.45 |
| | SAUSA300_0862 | *glpQ* | Glycerophosphoryl Diester Phosphodiesterase | 2.2E-05 | 1.18 |
| | SAUSA300_2096 | *manA* | Mannose-6-Phosphate Isomerase | 2.5E-04 | 0.60 |
| | SAUSA300_1014 | *pycA* | Pyruvate Carboxylase | 4.3E-03 | 0.46 |
| | SAUSA300_1192 | *glpK* | Glycerol Kinase | 1.9E-01 | 0.21 |
| | SAUSA300_0151 | *adhE* | Bifunctional Acetaldehyde-CoA/Alcohol Dehydrogenase | 1.8E-03 | -0.47 |
| | SAUSA300_1678 | *fhs* | Formate-Tetrahydrofolate Ligase | 1.1E-03 | -0.50 |
| | SAUSA300_1657 | *ackA* | Acetate Kinase | 2.5E-04 | -0.66 |
| | SAUSA300_1510 | - | 5-Formyltetrahydrofolate Cyclo-Ligase | 1.0E-03 | -0.79 |
| | SAUSA300_2254 | - | Glycerate Dehydrogenase-Like Protein | 1.5E-05 | -1.07 |
| Cell wall organisation | SAUSA300_2589 | *sasA* | Cell Wall Anchor Domain-Containing Protein | 4.2E-06 | 1.98 |
| | SAUSA300_0958 | *lcpB* | LytR-CpsA-Psr attach WTA to peptidoglycan | 5.2E-04 | 0.63 |
| | SAUSA300_1574 | *reoM* | Regulator of murA Degradation | 2.7E-04 | 0.58 |

| | | | | | |
|---|---|---|---|---|---|
| | SAUSA300_0876 | - | OatA Homologue, Predicted Peptidoglycan O-Acetyltransferase | 2.1E-02 | -0.39 |
| | SAUSA300_0703 | *ltaS* | Lipoteichoic Synthetase | 7.0E-04 | -0.64 |
| | SAUSA300_0248 | *tarF* | Putative Teichoic Acid Biosynthesis Protein F | 2.6E-04 | -0.76 |
| | SAUSA300_0625 | *tagG* | Teichoic Acid ABC Transporter Protein | 1.0E-05 | -1.73 |
| | SAUSA300_0992 | - | Predicted Cell-Wall Binding Lipoprotein | 1.4E-06 | -2.48 |
| | SAUSA300_0602 | - | Hypothetical secreted Protein, Linked To Response To Antimicrobial Peptides | 6.0E-07 | -3.24 |
| Cofactor and carrier biosynthesis | SAUSA300_0229 | *fadX* | Putative Acyl-Coa Transferase FadX | 4.2E-03 | 1.05 |
| | SAUSA300_0822 | *sufB* | Fes Assembly Protein SufB | 1.0E-01 | 0.20 |
| DNA replication, recombination, and repair | SAUSA300_0001 | *dnaA* | Chromosomal Replication Initiation Protein DnaA | 9.1E-05 | 1.05 |
| | SAUSA300_1259 | *umuC* | Impb/Mucb/Samb Family Protein | 1.0E-04 | 0.90 |
| | SAUSA300_1344 | *dnaD* | Putative Dna Replication Protein DnaD | 4.4E-03 | 0.41 |
| | SAUSA300_0383 | *saoC* | DNA Binding Protein of the Stress Associated Operon | 2.5E-01 | -0.13 |
| | SAUSA300_1362 | *hup* | DNA-Binding Protein HU | 2.4E-05 | -2.11 |
| Metal ion homeostasis | SAUSA300_0078 | *copA* | ATPase Copper Transport CopA | 4.1E-05 | 1.32 |
| | SAUSA300_1005 | *mntH* | Manganese Transport Protein MntH | 1.0E-04 | 0.82 |
| Nitrogen utilisation | SAUSA300_2238 | *ureA* | Urease Subunit Gamma | 2.4E-02 | 0.98 |
| Oxidative stress response | SAUSA300_1895 | *nos* | Nitric Oxide Synthase Oxygenase | 9.1E-04 | 0.55 |
| | SAUSA300_0253 | *scdA* | Cell Wall Biosynthesis Protein ScdA | 2.9E-04 | -0.72 |
| | SAUSA300_1463 | - | Bacilliredoxin homologue | 1.9E-04 | -0.74 |
| | SAUSA300_1513 | *sodA* | Fe/Mn Family Superoxide Dismutase | 1.4E-05 | -1.14 |
| | SAUSA300_2626 | *bstA* | Bacillithiol transferase | 1.7E-05 | -1.22 |
| Protein fate | SAUSA300_1089 | *lspA* | Lipoprotein Signal Peptidase | 1.9E-01 | -0.15 |
| | SAUSA300_1278 | - | Oligoendopeptidase F | 2.0E-04 | -0.79 |
| | SAUSA300_0536 | *hchA* | Chaperone Protein HchA | 1.7E-04 | -0.80 |
| | SAUSA300_0777 | *cspC* | Cold Shock Protein | 7.9E-06 | -1.45 |
| Purine ribonucleotide biosynthesis | SAUSA300_0478 | *prs* | Ribose-Phosphate Pyrophosphokinase | 1.2E-03 | 0.68 |
| Ribosome and Protein synthesis | SAUSA300_0529 | *rplGB* | Putative Ribosomal Protein L7Ae-Like | 3.0E-04 | 0.77 |
| | SAUSA300_0522 | *rplK* | 50S Ribosomal Protein L11 | 9.9E-04 | 0.54 |
| | SAUSA300_1312 | - | Acetyltransferase | 4.6E-02 | -0.23 |

| | | | | | |
|---|---|---|---|---|---|
| | SAUSA300_1365 | *rpsA* | 30S Ribosomal Protein S1 | 3.1E-04 | -0.60 |
| | SAUSA300_2316 | *paiA* | Acetyltransferase | 1.1E-03 | -0.91 |
| | SAUSA300_1131 | *rpsP* | 30S Ribosomal Protein S16 | 1.0E-03 | -1.09 |
| Transcriptional regulation | SAUSA300_0350 | - | Cro/Ci Family Transcriptional Regulator-Like Protein | 2.7E-02 | 0.79 |
| | SAUSA300_0033 | *mecR1* | Methicillin-Resistance Mecr1 Regulatory Protein | 4.9E-01 | 0.10 |
| | SAUSA300_1453 | *rnz* | Ribonuclease Z | 8.5E-02 | -0.25 |
| | SAUSA300_1639 | *phoP* | Alkaline Phosphatase Synthesis Transcriptional Regulatory Protein | 3.4E-04 | -0.64 |
| Transport | SAUSA300_0435 | *gmpA* | ABC Transporter ATP-Binding Protein | 6.0E-07 | 2.58 |
| | SAUSA300_0107 | *nptA* | Na/Pi Co-transporter Family Protein | 1.0E-06 | 1.85 |
| | SAUSA300_0231 | *nikA* | Nickel Substrate-Binding Protein for ABC transporter | 8.5E-01 | 0.02 |
| | SAUSA300_1979 | *ktrB* | Potassium Uptake Protein B | 7.0E-03 | -0.43 |
| | SAUSA300_1283 | *pstS* | Phosphate-Binding Protein of Phosphate ABC Transporter | 3.3E-01 | -0.45 |
| Virulence | SAUSA300_2366 | *hlgC* | Gamma-Hemolysin Component C | 1.8E-05 | 2.34 |
| | SAUSA300_0099 | *plc* | 1-Phosphatidylinositol Phosphodiesterase | 1.9E-06 | 1.52 |
| | SAUSA300_0224 | *coa* | Staphylocoagulase | 1.7E-05 | 1.37 |
| | SAUSA300_2440 | *fnbB* | Fibronectin Binding Protein B | 9.5E-03 | 0.43 |
| | SAUSA300_0548 | *sdrE* | Serine aspartate repeat protein E | 4.1E-01 | -0.11 |
| | SAUSA300_1370 | *ebpS* | Cell Surface Elastin Binding Protein | 2.0E-01 | -0.21 |
| | SAUSA300_1101 | *rqcH* | Putative Fibronectin/Fibrinogen Binding Protein | 6.1E-03 | -0.36 |
| | SAUSA300_1922 | *sak* | Staphylokinase | 6.4E-03 | -0.36 |
| | SAUSA300_1028 | *isdB* | Iron Regulated Surface Determinant Protein B | 2.2E-03 | -0.65 |
| | SAUSA300_2581 | *sasF* | Putative Surface Anchored Protein | 2.9E-06 | -1.35 |
| | SAUSA300_1296 | *msaA* | Modulator of SarA | 3.8E-05 | -1.53 |
| | SAUSA300_0113 | *spa* | Immunoglobulin G Binding Protein A | 3.4E-05 | -1.60 |
| | SAUSA300_0136 | *sasD* | Cell Wall Surface Anchor Family Protein | 0.0E+00 | -5.82 |
| | Teg41_srn_1080 | *NA* | sRNA Teg41 | *NA** | *NA** |
| Unknown | SAUSA300_0178 | - | Hypothetical Protein | 2.2E-04 | 0.72 |
| | SAUSA300_0464 | - | Hypothetical Protein | 8.0E-04 | 0.71 |
| | SAUSA300_0362 | - | Hypothetical Mechanosensitive Channel Protein | 5.0E-03 | 0.52 |
| | SAUSA300_0606 | - | Hypothetical Membrane Protein | 1.8E-02 | 0.46 |
| | SAUSA300_0371 | - | Hypothetical Protein | 2.4E-01 | 0.37 |
| | SAUSA300_0655 | - | Hypothetical Protein | 7.4E-03 | 0.35 |
| | SAUSA300_1534 | - | Hypothetical Protein | 1.7E-01 | 0.32 |

| | | | | |
|---|---|---|---|---|
| SAUSA300_1339 | - | Hypothetical Protein | 1.0E-01 | 0.29 |
| SAUSA300_2456 | - | Hypothetical Protein | 9.5E-01 | -0.01 |
| SAUSA300_1905 | - | Hypothetical Protein | 8.6E-01 | -0.02 |
| SAUSA300_0390 | - | Hypothetical Protein | 5.3E-01 | -0.11 |
| SAUSA300_1846 | - | Hypothetical Protein | 6.7E-03 | -0.48 |
| SAUSA300_1902 | - | Hypothetical Protein | 2.1E-04 | -0.64 |
| SAUSA300_1026 | - | Hypothetical Protein | 2.7E-04 | -0.66 |
| SAUSA300_2592 | - | Hypothetical Protein | 3.2E-03 | -0.66 |
| SAUSA300_0365 | - | Hypothetical Protein | 1.9E-02 | -0.72 |
| SAUSA300_0657 | - | Hypothetical Protein | 1.7E-04 | -0.73 |
| SAUSA300_0024 | *walJ* | Metallo-Beta-Lactamase Family Protein | 2.0E-04 | -1.20 |
| SAUSA300_0374 | - | Hypothetical Protein | 9.6E-05 | -1.49 |
| SAUSA300_1070 | - | Hypothetical Protein | 3.4E-06 | -1.78 |
| SAUSA300_1582 | *csbD* | Similar to general stress response protein | 1.1E-05 | -1.83 |
| SAUSA300_0682 | - | YbaK/EbsC Protein | 8.1E-05 | -1.86 |
| SAUSA300_0985 | - | Hypothetical Protein | 1.2E-04 | -2.14 |
| SAUSA300_1965 | - | Hypothetical phage Protein | 1.7E-04 | -2.39 |
| SAUSA300_0031 | *maoC* | MaoC domain Protein in SCC*mec* | 2.9E-06 | -2.93 |

***Table S3. The predicted WalR direct regulon.*** *S. aureus NRS384 genes with in silico detected WalR binding sites within 500 bp of transcriptional start sites. RNA-seq analysis of gene expression comparing NRS384 Wt with the WalKT389A 'up' mutant is shown for each locus. FDR = P value adjusted for false discovery rate, Log2FC = fold-change.*

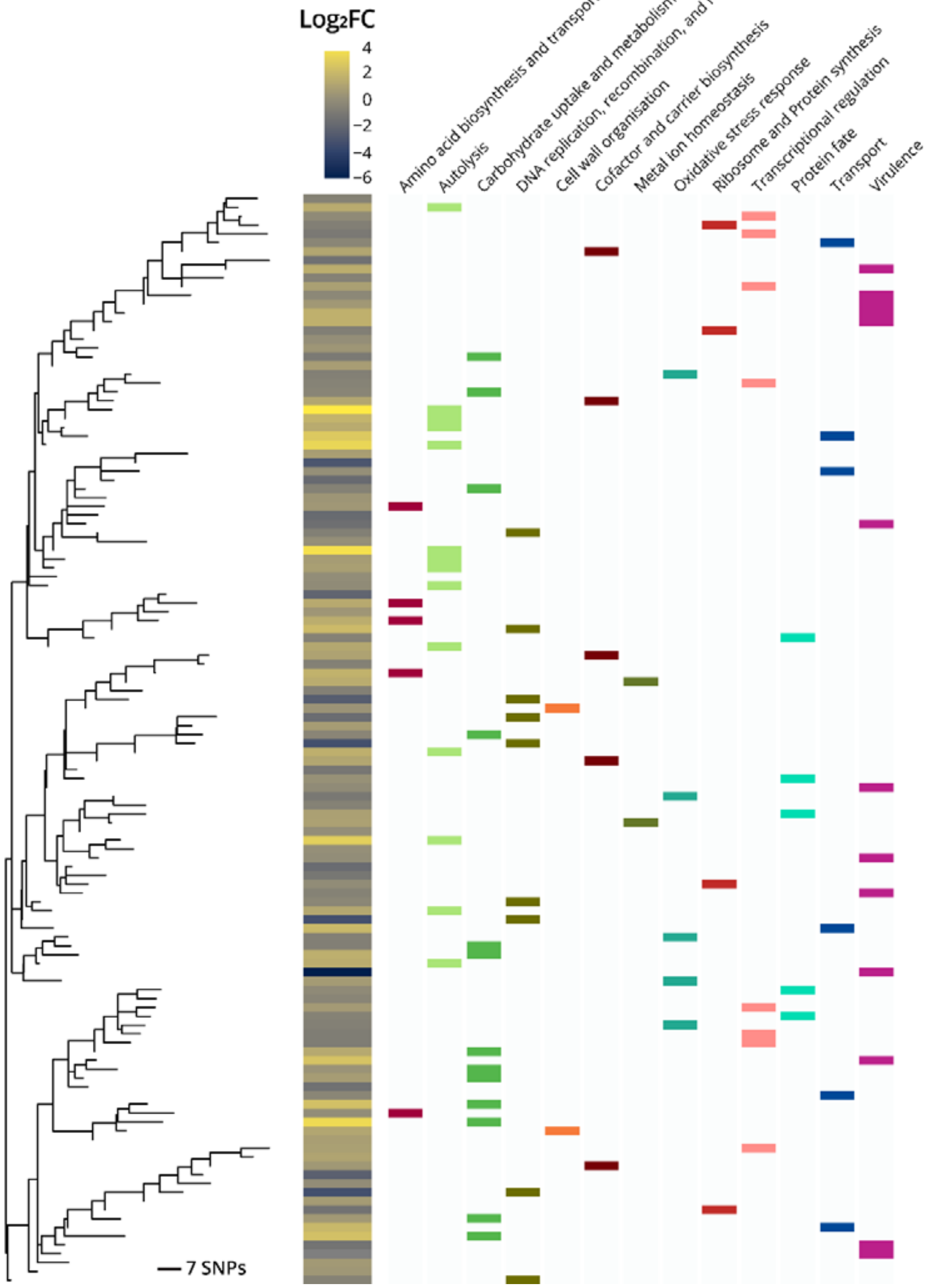

***Figure S4. Diversity of WalR motifs****. Analysis of intergenic WalR motif diversity to investigate grouping by alterations to gene expression (left) or the functional class of regulated genes (right).*

| Gene | Locus tag | Direction | Log2 FC | Sequence |
|---|---|---|---|---|
| - | SAUSA300_2253 | Rev | 3.90 | CTAATTTTTATTACAAAATTAGTCAAAGTTTTTTTATTCTTTTACAAATCAATAACAATTC**CCCACA**AGAAGCATTAATTTTGT**TAAAGT**AACTATTGTCA( 79nt)ATG |
| *sle1* | SAUSA300_0438 | Fwd | 3.50 | AATTATAAAATTTGATGATACAGTATATGATTTTTTTGTAATCATAATGTCATCAAACA**TCAACC**TATTATACATAATAAAATCG**TATAAT**GATGTAGTA( 56nt)ATG |
| *uhpT* | SAUSA300_0216 | Fwd | 3.26 | AATATGTTTCTAAGTATGTGTTTATGTTCAGTATTTTGGATAATTTAATAATTTTAAGGATATTAAGCGCTTACACCGACGTGA**TATATT**TGGCTTAACG( 21nt)ATG |
| - | SAUSA300_2503 | Rev | 3.09 | ATTGGTATATTACATTCAATGAAGCTTTATTAGGAACAGATTACATTATGATAACAAAG**CCCGCA**AGACACCTAATCTCTGT**TATAGT**TTGTTTTGTCGC(121nt)ATG |
| *atl* | SAUSA300_0955 | Fwd | 2.85 | TTTTTCATTTTTTTACAGTGAAAATGTAAATTAAGAGTATATTACAAATTGG**TTAAAT**ACGCACAGGTATATAAAACAGGTAC**TATAAT**GTTAGTAATAA( 28nt)ATG |
| *gmpA* | SAUSA300_0435 | Fwd | 2.58 | ATGTTAATATTTAGTAATTGCGTTGTGGAGGATTGGTGGACATTTTTATGGATTTAATTTGATAAATGTCATAGTAGTCTCACAATTCGTCATTGTCACATGAATTACTT ATTTTTTAATTTTTTAGAAAATTCGGCAAATTGTA**TTGACG**ATATATGCAATTTTACA**TATAAT**GCTCTTATTC( 36nt)ATG |
| *hlgC* | SAUSA300_2366 | Fwd | 2.34 | TGGCATTTTTAACTTAATTGTAAAAAAAGTTGATAATGGTCAATTGTTAATGAACAG**TTAATT**ATAATAACGCCCAAAATA**TATTATTA**TTTAATTAAGT( 31nt)ATG |
| *lacA* | SAUSA300_2155 | Fwd/Rev | 2.20 | TGAATGTTCACTTATTGTTTGAAAATACGTCATAAACAAACATATTATAAACATATTATG**TTGATT**TATTGTTTGTTTATGTT**TATAAT**TTAAACGTAAT( 99nt)ATG |
| *sasA* | SAUSA300_2589 | Rev | 1.98 | ATTACACATATATTACACTAGGTTTATTGAAAAAATATTGGTATTACTTTAATATTAAAGATGTATTTTGCATATAATATTTATCGTTTAGT**TGTAAA**TTAACTATTTAT AAATAAAAGGTGGATGGTAT**AGATATATT**AATACAATAA(255nt)ATG |
| *nptA* | SAUSA300_0107 | Rev | 1.85 | CGATATTTTACAAGTCATATACAAATAACATATATTGTTAAATAATTTTACCTAATCTTAACATTAAAT**TTACA**ATTATAAGC**GATAAT**CTAAATATAAA( 23nt)ATG |
| - | SAUSA300_0651 | Rev | 1.68 | TTATAAAAATATGAACCGATATCCTAAAATGTTAATAATATTACAAGATAATAACAAAC**CACACA**AAGCTACTTATTTTTGA**TAATAT**GGAAATCGTAAT( 83nt)ATG |
| *metX* | SAUSA300_0012 | Rev | 1.60 | CGCATGACAATGATAACAACATTTTAAATGATAAAAGTAATTCATCACTGAATCTCAACTAACAC**ATAACA**ATTTCATATT**TCTTATTGT**GAGAAGTTGA(138nt)ATG |
| *plc* | SAUSA300_0099 | Rev | 1.52 | TGTCAATAATTTCTATAATACATTATTAAAACATCAATTAAATAAGTTTTAAAAT**TTTACA**CATATTTTTATTTAAAAAAGATG**TATAAT**TAATGTATTA( 23nt)ATG |
| - | SAUSA300_0277 | Fwd | 1.48 | GTATTTTTAGCTTTTTTTAAATAAATATATTGAATATACCCATATATTTTTAA**TTAACC**ATTCATTTTTGTAATATAAATGTG**TATACT**AAAATTAAATT( 27nt)ATG |
| *glpF* | SAUSA300_1191 | Rev | 1.45 | TTAAACTACTTTTATAAAGACTTGAAAAATTAAATTTGACAAAATTTTTACAAAAACCA**TTGACA**ACGCTTTCATATCGATAG**TATCAT**AACAAATATAA( 92nt)ATG |
| *glnP* | SAUSA300_1808 | Rev | 1.38 | ACAGTCTTTTAAAAAGAAGCTTTTCTTTATATTGATAAAAATACATTTATAAAACACT**ATTGCA**TCAAAATGTATTAATATG**TATAAT**TATGAATATTAA( 18nt)ATG |
| *coa* | SAUSA300_0224 | Fwd | 1.37 | TAATTAACTATAACAAAAAGATAGTTAATGCTTTGTTTATTCTAGTTAATATA**TAGTTA**ATGTCTTTTAATATTTTGTTTCTT**TAATGT**AGATTGGGCAA( 26nt)ATG |
| - | SAUSA300_2482 | Rev | 1.34 | TACAATTCAACACTTTATATGACAACTTCATTACAGTTACTTTTATTGTTGATTG**CTTACA**TTGTTTTCTAAAAAAAATTTGTTAT**CATAAT**TAACGTTG( 32nt)ATG |
| *copA* | SAUSA300_0078 | Fwd | 1.32 | TAACCTTGGTTAGAGTGTATTGTCATGTGATAGTTAAGATTAGATCTTTTAATTAAAACA**TTGTTT**TAAATATCGATGACA**AGGTCTAAT**GTAGGACGTG(125nt)ATG |
| *ssaA* | SAUSA300_2249 | Rev | 1.31 | ATTACAAATTTGTAACAGACTTATTTTCGAAACATTCAGTTTTCACAAAGAATGCCGTAGTTATAAGTTTTGCAAAATGCAAATTTCTATACTTTTAAAATTCTATTGTG TCAAGATATTAGCAAAAAATTACAAAACGTTAACAA**AGCACA**CAAGGACGCTAATTTCTGT**TATAGT**CTTTCTTGTCG( 94nt)ATG |
| *rocD* | SAUSA300_0860 | Fwd | 1.21 | TGTATGCAATTTGTTAGTGGTTTTCAATATTTATAAATAAAAATATTCTTAGTAAAATATACTTTTGATTAAGTGTAAAATTATCAATTCTTTTGC**TTGTAA**TCATTTAA GTAATTATGT**CATTAT**AAATGTAAGG(142nt)ATG |
| *glpQ* | SAUSA300_0862 | Fwd | 1.18 | TAATTCTTTTAAATTTATATTACAAAAATGTTATAAATTGTAAAAGAAATGTGTAAAGCGTT**TTCACA**AGCAGGTTTTTGTAG**TATTTT**AAAATTGTTAG( 32nt)ATG |
| *isaA* | SAUSA300_2506 | Rev/Fwd | 1.09 | TATTACAGCTATGTAACAAAAATACAATCTGTAATATTACGAAAGCTGTAATAGTTGAGCTAAATATAGTAAAAATCTTATGTGTTAAAGACTATTTCAAGGGTTGATTG GATTCTAAAGGGCACATATTTCACATTACAGTATAAAACAGCCATATTTCAACATTACAAATAACACTTGATATTGTAATGTTTTGTAAAGAAAGTGTAA**TTTACT**GGCT GGTTTTTTGTGA**TATAGT**GAGTACCGTTGA(100nt)ATG |
| *dnaA* | SAUSA300_0001 | Rev | 1.05 | GTGGATAATTAGAAATTACACACAAAGTTATACTATTTTTAGCAACATATTCACAGGTAT**TTGACA**TATAGAGAACTGAAAAAG**TATAAT**TGTGTGGATA(200nt)ATG |
| *fadX* | SAUSA300_0229 | Fwd | 1.05 | TTATCGACCGTCAAAATGTGTCTATACTGTGTCTATTATCAGAATAAAAAGATATTTCTAA**TTGATT**TTAAACGTCGTTATGT**TATATT**TCTTGTAAGGG( 40nt)ATG |
| *ureA* | SAUSA300_2238 | Fwd | 0.98 | TCAATATCGGATATGCTAAAAATATAATTTTCTAATTACACATTGAACTTTGTAT**ATTACT**TGTTTATCACAACAATTAGGATTAA**TATAAA**TTTTGTAA( 32nt)ATG |
| *umuC* | SAUSA300_1259 | Fwd | 0.90 | TATATCATGTAATACTTAAAAATTCATTTGCAAAATAACAGAACATTTGTTCTAAAATAG**TTGAAA**TAGAACACGTGTTCGTA**TATAAT**AAGATTCAAGT( 15nt)ATG |
| *mntH* | SAUSA300_1005 | Fwd | 0.82 | GAACGTTCAACATAATAATTCATACTTTTAAAAAAATTAAAATAAATTTAGG**TTGACC**TAAACATTTTATTAGGTTAT**TATATT**GTCCATAAGAAGTAGAA( 11nt)ATG |
| - | SAUSA300_0350 | Fwd | 0.79 | CATAGGTGGATAATCATATTTTTTGGTGTAGATTTGACAAATATAG**TTGTCA**AAAAGACGCATATCATT**TACAGT**GTAATTAAGGAGTTGATGCGATGTG( 12nt)ATG |
| *rplGB* | SAUSA300_0529 | Fwd | 0.77 | ATATAACAGAGGCTAATGCTTTAGCCTCTTGTTATTTTTATGTAAATTATTTGATTTAATG**TTGACG**AATTCTCTTGTTCAA**TGTTAAT**ATATTAAAGGT( 31nt)ATG |
| *lytM* | SAUSA300_0270 | Fwd | 0.73 | TGTAATGACAATGTAATGAGTTTAGTAAAACATTTCGGGAATATTAAATAGTTGGAAAATGAGAATTAAATCC**TTTACT**CAGTTGTCTAATTCTTT**TAGTAT**GTGCAGTA CAG( 45nt)ATG |

| | | | | |
|---|---|---|---|---|
| - | SAUSA300_0178 | Fwd | 0.72 | TAATTTGTTATATCCTTTTAACTAGGAAAATATACATTTCGTAATAATAATAATCGTTATCA**TTGAAA**AAGTGTTAATAAGGTG**TATAAT**GAAAATGTGA( 30nt)ATG |
| - | SAUSA300_0464 | Fwd | 0.71 | TGTTATTATTAAGTGTGCACGCAGTATCATTAGTTATAAAATGTAGCTGTTAAAAG**TCAAAA**ATACATCGAATGTAGTTAG**GCATATAAT**ATAAAAAGAG( 46nt)ATG |
| *prs* | SAUSA300_0478 | Fwd | 0.68 | TAGGATAAAAGGATAATCCTATGTAATATTAATGTAATCTTTATGATTTAATGA**TTCGCA**TAGTAATGGAGTTACATTTTATA**TATAAT**AGTAATTGCGT( 24nt)ATG |
| - | SAUSA300_0958 | Rev | 0.63 | AAAAATGCTTTTTGATTTTAACAAAACATTCAAATTACAAGGAACTTTGACATAACA**TTTGTA**TTTTTTACTATAAAGTAC**TACAAT**TTAAGGCTATAAT( 95nt)ATG |
| - | SAUSA300_0739 | Rev | 0.61 | ATGATAAAAGGTTGTATTCTTTTTATATTGTTAACCATTTGATTACATCGTTATAACAATAGC**TTTTGA**CAAAATGTATTGTGC**TATAGT**ATTTGCATAC( 33nt)ATG |
| *manA* | SAUSA300_2096 | Fwd | 0.60 | TATAAATTTTTGAAAATGTTAAAGTACTGTAAATTTAAGTGAAGCGCTTTAA**TTGGCA**GTACTGCAATGTTAGTAAATATGG**TACAAT**CACTAATAGTTA(111nt)ATG |
| - | SAUSA300_1574 | Fwd | 0.58 | GTACACAACCTGAAAATATCTCAAAATCATTAAGCTTTATTAAAGATTACATTAAAAA**TCTATA**ATTTAAAGCCATAATATTG**TATGAT**GATAGTGAGTT( 27nt)ATG |
| *nos* | SAUSA300_1895 | Rev | 0.55 | TTAATTTCCATACTTTTAATTAAAATCAACCAACAATTTAATGACATATACATAATTT**TTAAGA**GTATTTTAATAATGTAGAC**TATAAT**ATAAAGCGAGG( 8nt)ATG |
| *rplK* | SAUSA300_0522 | Fwd | 0.54 | ATAGAAAAGCTTTAATTAACAATTAAAGTTATTAAACTAACCAAAAGATAAAAAAGAGTA**TTGATT**TTTTAATTAGAAAAGTGT**TAAAAT**TATGTGGTCG( 95nt)ATG |
| - | SAUSA300_0606 | Fwd | 0.46 | ATTGTCTAGGGTGATTTTGTTGTAATAATGTCCTGTTTTTTTCAACATTATTGTAAATGAGT**TGTATA**TTTTCTCTATAAT**GATAAT**GTAAATATATGAT( 36nt)ATG |
| *pycA* | SAUSA300_1014 | Fwd | 0.46 | CTGAAGCGGAGGTTTTAATTGTCTCAAGAATGTTAAATTTTGAAAATTTGAAACATAACTA**TAGCAA**AACAGAGCGCTTTAAGA**TAAAAT**TTTATTATCT( 42nt)ATG |
| *fnbB* | SAUSA300_2440 | Rev | 0.43 | AAAAACTCATTATACATAAAGTTAACAACAATCTTAACTTTTTATTAACTCGCTTT**TTTTCA**TTGCTTTTAAAAACCGAACAATA**TAGAAT**TGCATTTAT( 47nt)ATG |
| *dnaD* | SAUSA300_1344 | Fwd | 0.41 | TTAAAAAAATACTGTTATTTAAAATTTGTTTGCTACTAGTTAAATATATTAAGCATTTTAG**TCGAAA**ATTAAAGAAAATGATT**TATACT**ATTAAGTAATGG( 24nt)ATG |
| - | SAUSA300_0451 | Fwd | 0.39 | GTTTGAATGATTTTTGTGTCAATGAAAAGTAAGAAGTTATAATTTGATGATAAAGAAATGATG**GTGAAA**TGAGGGGGAGTATCT**TACAAT**AGAATTATTA( 47nt)ATG |
| - | SAUSA300_0371 | Rev | 0.37 | TAACTTAGTTTAAAAACGATTCGTATCTTTCAGATTCAAATACCATCATTTTCTCCTAATA**CTTACA**CTTTAATTACAATT**ATGTAAGTT**GTTTTCAGAT( 34mt)ATG |
| - | SAUSA300_0655 | Fwd | 0.35 | TTTTAAAACTAAGTAACAGTTTGAAGAAATCGTAGTTCAATAATGTTAATTGTGAAAA**TGTATA**TAAACATAAAAAAATCATG**TATAAT**ATATGTTGTTA( 29nt)ATG |
| - | SAUSA300_1534 | Fwd | 0.32 | ATTAAAAAAAGTTATATAGTTTATAAATAATCAAATTGATATTCTATAGGTTCTTATAAC**TATAAA**GTATATTCAATTTCATG**TATAAT**TAATGTGAGGG( 36nt)ATG |
| - | SAUSA300_1339 | Rev | 0.29 | ATTTAACAGATAACACATTTGAAAAAATAGTTAAATCAAATTATATAGAGTGTTAAAAC**ATGACG**AATACAGTATATTGTGA**TAAATT**AAAAAATGTAGG( 10nt)ATG |
| *glpK* | SAUSA300_1192 | Rev | 0.21 | AATAAAAAGAAACGTAAATAGCATAATTTAACATGTTTGATTCATGGATTATGCTATTTTT**TCGCCA**AAATTTAACAGATTTTG**TACAAT**GGGTTAGCGA( 28nt)ATG |
| *sufB* | SAUSA300_0822 | Fwd | 0.20 | GAATAGAATGCTGTTAATCATAGATAATTTTGATATTAGACATATAAAAGTATAAAAATT**TTTATA**AGATGTCATGTCATTGT**TATAAT**ATGGTTTACAT( 56nt)ATG |
| *mecR1* | SAUSA300_0033 | Fwd | 0.10 | TTTTATCTTTTTCATCAATATACTCCTTATATAAGACTACATTTGTAGTATAT**TACAAA**TGTAGTATTTATGTCAAAATAATGT**TATAAT**TTTTGTGATA( 14nt)ATG |
| *opp-5A* | SAUSA300_0231 | Rev | 0.02 | TAAAAAGCATAATTACTACAATTAATTGAACTTTAATAATTACTAAACTTGAACAACAT**TTTACT**TTAACAAAATAAAGTTT**TAAAAT**ATTAATTGTTGG( 99nt)ATG |
| - | SAUSA300_2456 | Rev | -0.01 | TTAATGAAATGACCGACACCCTCGCTTCTACTCCAATTTAACAAAATACTATA**TTAGTA**CTATATTTTTCATTAACATGTATGT**TACATT**TTCTATATTA(204nt)ATG |
| - | SAUSA300_1905 | Rev | -0.02 | ACCTCAAACAATTTTATATTTCATTTAGTAAAATATAACTTTATTTAACATGACATTCAA**TATACA**TACGATTACTATTTATT**TACAAT**ATATTACAACT( 21nt)ATG |
| *putA* | SAUSA300_1711 | Rev | -0.10 | TCAACATCAATAATAGTGTAATTATACATAATTATTTTTGATTGTTTTTGATGAAAACGCT**TTCTCG**AATATTTTTTTCATGC**TAAACT**TATTGTAAACA( 21nt)ATG |
| - | SAUSA300_0390 | Rev | -0.11 | TATAAGCCACATGTCACTATTATCGCAGCTAGCTGCGTCACTCAAAACGAATACGGAACGCG**TTGGTT**ATTTCAATCATAA**TATTACTCT**GCAAATACAC( 74nt)ATG |
| *sdrE* | SAUSA300_0548 | Fwd | -0.11 | TTAATTTATTCAAACCAAAGTGTAAAATAACAGTCTTAGATAAATAAATTTATTTAAAGTA**TTGTGC**TTTATCTAAAAATGTAT**TACGAT**GGGAATACAA( 59nt)ATG |
| - | SAUSA300_0383 | Rev | -0.13 | TTTAAACAAACTTGAAATCACTAACAATAACGTTTAAACTATGTATGAAGCAAGACTGACGT**TTCACT**GCCCGTAGACAAA**CGTCAGTAT**ACATAAAGTA( 22nt)ATG |
| *lspA* | SAUSA300_1089 | Fwd | -0.15 | TGTCAAAACATAGTAGTTTATCAAGTATTGAGTTAGTAACATTAGATTTAATGTAATATG**CTTACT**TTTTTTATTAGCAGGTGTAAGC**TATAAT**ATAAAGAGTT( 31nt)<br>ATG |
| *sceD* | SAUSA300_2051 | Fwd | -0.18 | TATGGGGGATTTCGTTGTAATCACTGTGTAAAAAACAAGGCTATTTTTTTATTTTG**TTGTAA**CCTGCACGGATTTTATAGATGT**TACAGT**AGTCTCTGTA( 44nt)ATG |
| *ebpS* | SAUSA300_1370 | Fwd | -0.21 | AATTAGTGTATACACAATTGAAAAATGATTGAAATAATTTTGAAAAATATACATAAACAT**ATGTCA**TGTGGGTATATTTTATG**TAAAAT**CATTGTAATAG( 19nt)ATG |
| - | SAUSA300_1312 | Fwd | -0.23 | CTTTTATTGGTTATTTTTATCCCATAGTGTGATAATTACTATTTTTCATTCATAATAAAGG**TTTAAA**GCATGTTAATAGTGTG**TAAGAT**TAACATGTACT( 40nt)ATG |
| *rnz* | SAUSA300_1453 | Rev | -0.25 | GAAAAATTAACTAGCAAAGACGAATTTTTAACAGATTTTGATTCAAGTATAAA**TTAAAA**CTAAATTGATACAAATTTTATGA**TAAAAT**GAATTGAAGAAA( 13nt)ATG |
| *rqcH* | SAUSA300_1101 | Rev | -0.36 | ATTGAATTGCAAACTTTTAGATAATGTAAAATGTATGGCATAATGTATGGTTCAATAAC**TATACT**GAAAAGTTACAATCATGT**TAAAAT**GAAACGAATGA( 24nt)ATG |
| - | SAUSA300_0876 | Rev | -0.39 | TTCCAAGAGTGTACAATTCATTTTCTAGTTGAATATTTTCTTGAAAATTTTTATAATAAATTTAATTTATATTACAGTTATAT**TACAAT**ACATAATACTT(216nt)ATG |
| *ktrB* | SAUSA300_1979 | Rev | -0.43 | TCATTTTCAATTATAAAATATTAAAAGAATTAGTCAACGCCTGTTAGTAATACACATCA**GTAACA**TTTCTATTTTCATTTATGA**TATTAT**CTAATTATTA( 30nt)ATG |

| | | | | |
|---|---|---|---|---|
| *pstS* | SAUSA300_1283 | Rev | -0.45 | AAGCATTTTAATTTTACTAATGAAGCAATATTTTTTAGATTAACAAAAATTAATA**TTTACA**TTTTCTTAACAATTTTTTATG**TAACAT**TTACAGTTTCTA( 84nt)ATG |
| *adhE* | SAUSA300_0151 | Rev | -0.47 | AAAACACCGAAATAACAATGATTTCATGAAAACATTTATTTTAAAATTTGATATTTGTTCAAATAATATTCGAAATTAAACTTTTTTGTATAGAATTTTC**TTTATA**TCCT GAGAGACATG**TACTATAAT**GTTTGTGAAA( 30nt)ATG |
| - | SAUSA300_1846 | Fwd | -0.48 | ACTAAAGTTTTACATCGCTATCAGTATTATGTATGATTTATTTTAACAACTATAAATAAGC**TTGAAT**TTTGTAACTAGTAAGTG**TATAAC**AGTAATGAAT( 45nt)ATG |
| *fhs* | SAUSA300_1678 | Rev | -0.50 | ATAATACAAAACTTTAATAAGTGAATTTATTGCAAAAATGAAAGCGCTAACCCGATTTAG**TCGACA**AGTTTTTAACAGTTCGT**TATTAT**ATGAATGTAAG( 67nt)ATG |
| *rpsA* | SAUSA300_1365 | Fwd | -0.60 | TAAAGTGACATATTTATGTATATGACTATTTCGCAAAATGTAATCGAGGTAGAATTTC**TTGACA**ATTCTGTCAGTTTATAAGA**TGTTAT**AAATATGTAGT( 19nt)ATG |
| - | SAUSA300_1902 | Fwd | -0.64 | CCTCTTTTTTACTTACACGAGCATTAATTATTAACGCGTTTATGCTTCCACAAAATTTTTG**TTGCAA**AATAGAACACATGTTCG**TATAAT**GTAATTATCA( 18nt)ATG |
| *ltaS* | SAUSA300_0703 | Rev | -0.64 | CAGTAAAATGAAAAGATAAAGTGTGTTTTTAC**TTGAAT**TTTGACTAAAATTACTC**TATATT**TATTAATTGAGCTATGCTTATTATTACAATTTGATTACA(262nt)ATG |
| *phoP* | SAUSA300_1639 | Fwd | -0.64 | TGTCCCAGAGTAAGTCATGCAAATGATAGAAGGAAATATAAGAAGTATTTTA**TTTATA**TGTAAAGTAATGTAAAAAATGTGTAAA**GATAAT**GTGTAGGAT( 61nt)ATG |
| *isdB* | SAUSA300_1028 | Fwd | -0.65 | AACCTATGTCATAGATATTTCATAATCTATAACATAGGTTATTTTTTTATAAAATAACG**TTGCAA**TTAACTAACATTTCAATG**TACAAT**ACAAGTAATCA( 52nt)ATG |
| - | SAUSA300_1026 | Fwd | -0.66 | ATTCGTCCTTTCGGTATTCATTATTAATATTGTAACAGAACATGATATGTTAAGAAAAAATC**TTGACA**ACTTGTTCTTAGAAAGT**TAAAAT**AAATTTTGT( 21nt)ATG |
| *ackA* | SAUSA300_1657 | Fwd | -0.66 | TGACAGAGTTAAATCAGTGGATGGACACAAATCGTCCTAAAAAAATAATCTGTAATAGTA**TAGTCA**TAAAACTGTATGAATGA**TAAAAT**GAAAAATGAAAT( 34nt)ATG |
| - | SAUSA300_2592 | Rev | -0.66 | GAGTGAACAGAATTATATTTCCATAGCAAACATTCCTAAACTACACATACGTTACATAA**TTGATT**CA**TTTTTA**TAGAAACGGGTAAAAATGATAAAGTAA( 40nt)ATG |
| *scdA* | SAUSA300_0253 | Rev | -0.72 | TTGTGAGCCAACATGATTTGAGGGCTTTATTTTGCTGTTTATGACATGATTATGACAT**TTCCCT**GATTTTCATTTTCATATA**CATTAAATT**GTATACACT( 22nt)ATG |
| - | SAUSA300_0365 | Fwd | -0.72 | TTCATATCATTTTTGATATTAATTCATTTGAAACTTTCTATGATATTTTTAAAAATACAC**TTCACA**AAAGCGAACATATGTTC**TATAAT**AGTTGTGAGGT( 8nt)ATG |
| - | SAUSA300_0657 | Rev | -0.73 | TAGTAAAATAATAATAAATATTTTCATATGATTAACAAAAACTATAAAACTGTATC**ATGACA**TGACATCATTTATGTAGTTTTA**TATAAT**AATGACAAGG( 11nt)ATG |
| - | SAUSA300_1463 | Fwd | -0.74 | ATAATGTTTGGTATATTGTTAAAATTGTGTCTAAATATAGGTGTGATTCAGATTAGTTTA**TTGAAC**AATATGTTATTAATTAG**TAGAAT**GAGGATAGTTT( 28nt)ATG |
| *tarF* | SAUSA300_0248 | Fwd | -0.76 | TGTTATGTTAAAGTGGTACATTAATCATGTATTTCGTATGATAATTAACGACAAGTGTAA**TGGTTA**AATGTATTTTATGATGAAATGC**TATAAT**AGGCATGGTT( 57nt) ATG |
| - | SAUSA300_1278 | Fwd | -0.79 | GCAATATTCTATCTACTAATAGAAAAATCATTGTTCCTTGCGACATGGAAATC**GTAACA**TTATCGTTTAGGAGACAAAATTATG**TATAAT**GAATGTATTA( 18nt)ATG |
| - | SAUSA300_1510 | Rev | -0.79 | AAGATTTTCAATTGAAATAATAAATTTTAGAATTGTCGCATAATTCATCATGACAACATA**ATGACA**TGTATTGTTATTAACGA**TATAAT**AGAATTGAATA( 14nt)ATG |
| *hchA* | SAUSA300_0536 | Fwd | -0.80 | ATTTAGTTGTATTTTTTTCAAAGAAATTCATTTTGATTATTTTTGATAATGAGCATT**TTAATA**GTAATACATGTTTATAGTG**TGTAGT**ATATGTCTATAC( 35nt)ATG |
| *paiA* | SAUSA300_2316 | Rev/Fwd | -0.91 | TGAAAATACGGCCATTTTACTGGAATAAATGTATCATTTATTGATGTTACACACAATC**AATACA**ATTTGTAATTGATTTGT**GTTAAT**GTATTTTTAAATA( 63nt)ATG |
| - | SAUSA300_2254 | Rev | -1.07 | GTTATATCACCTCCATCAAAAGTACGCGCTTACATCATTGATACACAAGACATC**ATGACA**TAGTTGTATACTGTCTAATCTTCTAC**TATACT**AAGATAAA( 14nt)ATG |
| *rpsP* | SAUSA300_1131 | Rev | -1.09 | TCACATAGTTATTATTGTAAGAAAAAGGTTCAATAATTTTCCTGGTAAAGAAAAAACTC**TTTACA**AACATTCATACACCTGTTAA**TATTAT**TTCTTGTAG( 40nt)ATG |
| *sodA* | SAUSA300_1513 | Rev | -1.14 | CATTTCATACTCCGTTCGTATGATTTTATGGCAAATCTGTTAACTTTTTATA**TACAAA**ATGTTTAATTATTTTTGATTTGAGT**TATATT**AAATGAGTAGA( 96nt)ATG |
| *walJ* | SAUSA300_0024 | Fwd | -1.20 | CGCATTTATTCATATTTTAAGTAGAACCGCATTGTAAAATTAGTGTAACTGTTATTTTAAAAAC**TTTAGT**ATTTGTCTAATCATTGT**TATAAT**AATTAAG(182nt)ATG |
| *bstA* | SAUSA300_2626 | Fwd | -1.22 | TTGAGAACTTTTCGTCAACTATCTTTTATTGTAAGGTAGTTGTTGTACACATTCCTTAA**ATGACT**AACAACTTTGTTAATAG**GGTAAT**ACTTACGGAAGT( 29nt)ATG |
| *sasF* | SAUSA300_2581 | Fwd | -1.35 | TTAAATATAAGCTAAGTAATAAGTAGATAATTACTAACAACAATAACTAGATAGATAAG**TGTAAA**TTTCTTGTAAACAGGTA**TATAAT**AGTATGTAATTC( 33nt)ATG |
| *cspC* | SAUSA300_0777 | Fwd | -1.45 | TATTAAAATGGAATGTTACTATATAGTTCAATGTGTATTATCACAGAAAATAAAATAATGC**TTTACT**TCTATATTTAAAAGTG**TATAAT**GAAAGTTAAGT(109nt)ATG |
| - | SAUSA300_0374 | Fwd | -1.49 | AATAACGATAGCTACATTGAATAAAATTGATATTCAATTACTACTTTTAAAAATATT**TGGATA**AAAATAATTTGAATTTGTTT**TAGAAT**TGTAAATAAGG( 50nt)ATG |
| - | SAUSA300_1296 | Fwd | -1.53 | TTTACTTAAAAAAACTAATAATTACTATAGTTACTACTTTGTTTGTTTCAAGTCGTCAAAC**TTGATT**TTCAGAGGATAAAGG**TATAAA**AATAAGTATAGA( 30nt)ATG |
| *spa* | SAUSA300_0113 | Rev | -1.60 | TTTATAAGTTGTAAAACTTACCTTTAAATTTAATTATAAATATAGATTTTAGTA**TTGCAA**TACATAATTCGTTATAT**TATGAT**GACTTTACAAATACATA( 13nt)ATG |
| *tagG* | SAUSA300_0625 | Fwd | -1.73 | TGATTGTATACTATAATGTATTTGTAATAAACTAATATTTTAAAGAACTAGACAATAATT**TTGATA**GCATCCATGTATAGTGA**TAGTAT**TTACAACAATT( 62nt)ATG |
| - | SAUSA300_1070 | Fwd | -1.78 | TGAAAATCAATTTAATCATGGAAAATTTATAATATTCATTGTTTTACATTTTCAAATCA**ATGAAA**AACACAAGTGGTTTAATG**TATAAT**AATAGTAGTAA( 21nt)ATG |
| *csbD* | SAUSA300_1582 | Rev | -1.83 | CATGTTTTATATTTAATCGTATAAATACTATACTATAACATAAAAACTTCATATTATAATGT**TTAGCG**AACCTCCTTAGTGGTA**TATAAA**TATATACATC( 15nt)ATG |
| - | SAUSA300_0682 | Fwd | -1.86 | TTAATTTAAAAAGGTGAATTCAACTTATAAAATGATGTAAATGTTATGTCAAAA**TCAACC**AATCCGTAATGTATTTTAAAATGT**TAATAT**AGTTCTGAAG( 21nt)ATG |

| | | | | |
|---|---|---|---|---|
| - | SAUSA300_0985 | Fwd | -2.14 | GTATTTTTAAAACATTATGCGCTATGAAATTGTGTATAATTTATGTCAGTTCACAATGTG**TTCACA**AATTTGAATTTCAATG**TATAAT**TGTGTATATTAC( 30nt)ATG |
| - | SAUSA300_1965 | Fwd | -2.39 | GTGATAAGTACATGTGTTTCAGGTATTGGAACTGTTACGTTTAAAATGT**GTGGCA**TTTCTATCTTTCCTTTCGTG**TATAAT**GTTGTTATCTCCTAGTGAA( 14nt)ATG |
| - | SAUSA300_0992 | Fwd | -2.48 | ATGACGTAACTGTCAACAGATATACTTAGTAATGAAGATGTGTAATGTAATTGTTTAAAA**TTGATT**TCCAAGCAGATTTTATT**TATCAT**TTAATTTAAAT( 20nt)ATG |
| *maoC* | SAUSA300_0031 | Rev | -2.93 | AGATAAAATGGTGCATTTGTCTCAATAGAAAGTTAGAGGTGAGTCTTACGTTTCAGTGACGG**TAGACT**TACCTTTAACATGTTACATACTAAAAAATTAA( 24nt)ATG |
| - | SAUSA300_0602 | Fwd | -3.24 | CAACAAACATCACGATGAAAAGCATGTTTAATTTTTAGTGATAAGTGAAAT**ATAAAA**GTAGTAATTATTATTTGTAA**TATAAT**TGTAATATGACTGTTGT( 66nt)ATG |
| *sasD* | SAUSA300_0136 | Fwd | -5.82 | AATCTAAAGTGAAATTTTTATAAAAAAATGTAATGATTCAAAAATTTTG**TTGCAT**TTCTTTTGTAATCGTAT**GATAAT**GTAAATGTAATCAAATTGTAAT( 16nt)ATG |
| *teg41* | srn_1080 | Rev | #N/A | TACAGGTGCATTTACAAAATCTTTACAATCATTGGTTTAATAGAGGTTAAGCTCATGAAT**TTGACA**TGAAGCAGAAATTTAAT**TATATT**TTATGTTATAA |

***Table S4. Relative positions of predicted WalR binding and transcriptional start sites (TSS).*** *Red; WalR binding, green; TSS , underlined; predicted -35 and -10 promoter elements, yellow; sRNA. Direction refers to the orientation of the WalR binding site in relation to the downstream gene.*

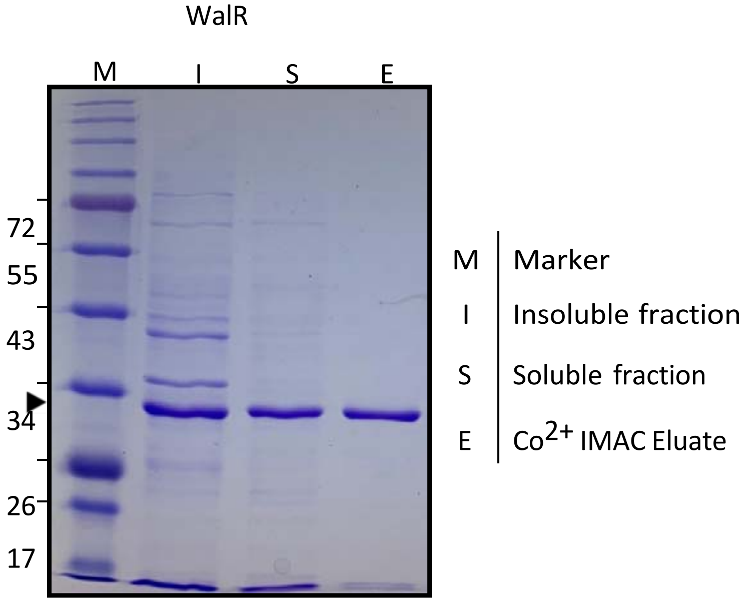

**Figure S5. Purification of recombinant WalR.** WalR was purified by single step immobilized metal-ion affinity chromatography using Cobalt TALON resin. SDS PAGE image shows cellular fractionation and purification. Black arrow denotes 6-His-WalR (28kDa).

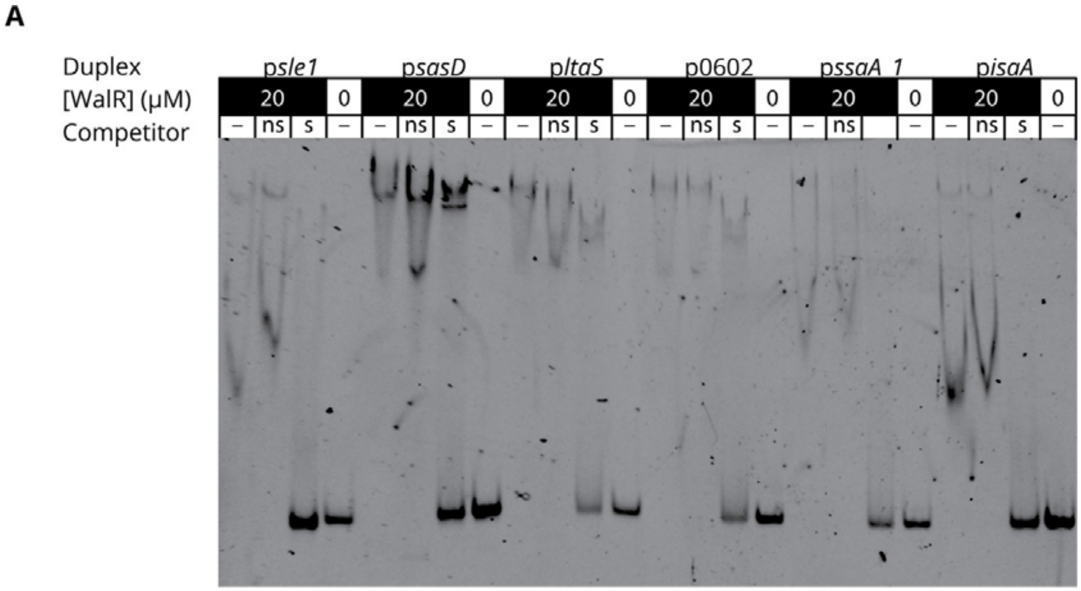

***Figure S6. DNA competition EMSAs with WalR.*** *Competition EMSAs show WalR binding to ChIP-seq identified promoter regions is specific. Competition electrophoretic mobility shift assay (EMSA) showing the effect on WalR binding of the presence or absence (–) of an excess of non-specific (ns) or specific (s).*

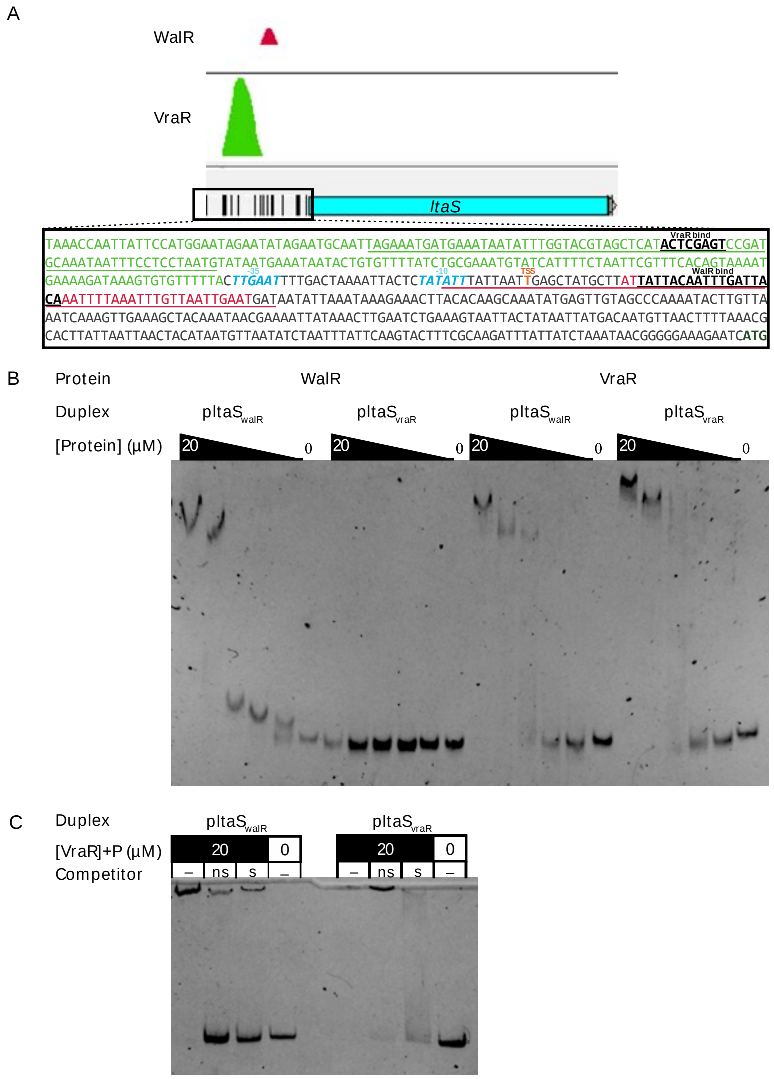

***Figure S7. WalR and VraR binding at the ltaS locus****. A) Artemis userplot showing VraR and WalR ChIP-seq peaks. Box shows sequence 529bp upstream of ltaS start codon. Bases coloured green correspond to VraR binding peak, red to WalR. Transcriptional start site in orange, -35 and -10 promoter elements are in blue, and VraR and WalR binding motifs are bold and underlined. Bases underlined in green and red correspond to positions of the pltaSvraR and pltaSwalR duplex probes respectably. These probes were used for EMSAs (see below). B) Dose dependent band shifts caused by WalR (left) and VraR (right) binding of ltaS promoter regions.C) Competition EMSA showing band shifts for phosphorylated VraR in the presence or absence (–) of an excess of non-specific (ns) or specific (s) non-labelled duplex competitor*

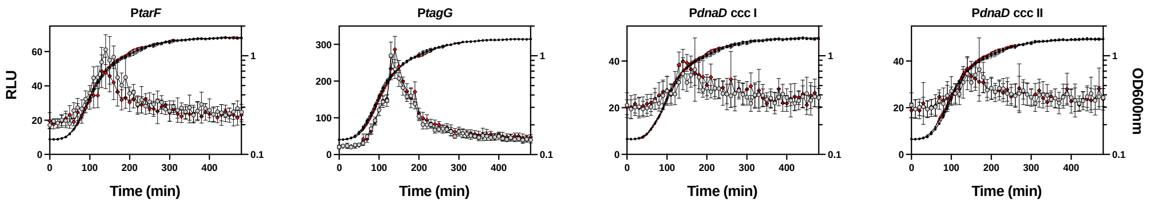

***Figure S8.*** **Impact of WalR motif mutation on the expression from *tarF, tagG and dnaD.*** *The above genes coupled to bacterial luciferase reporters showing the changes in promoter activity of the Wt (open grey circle: RLU; filled black circles: OD600nm) or the ccc (open red diamond: RLU; filled black diamond: OD600nm) mutated WalR binding site in LB media over time. Data represent the mean of three independent experiments (±standard deviation).*

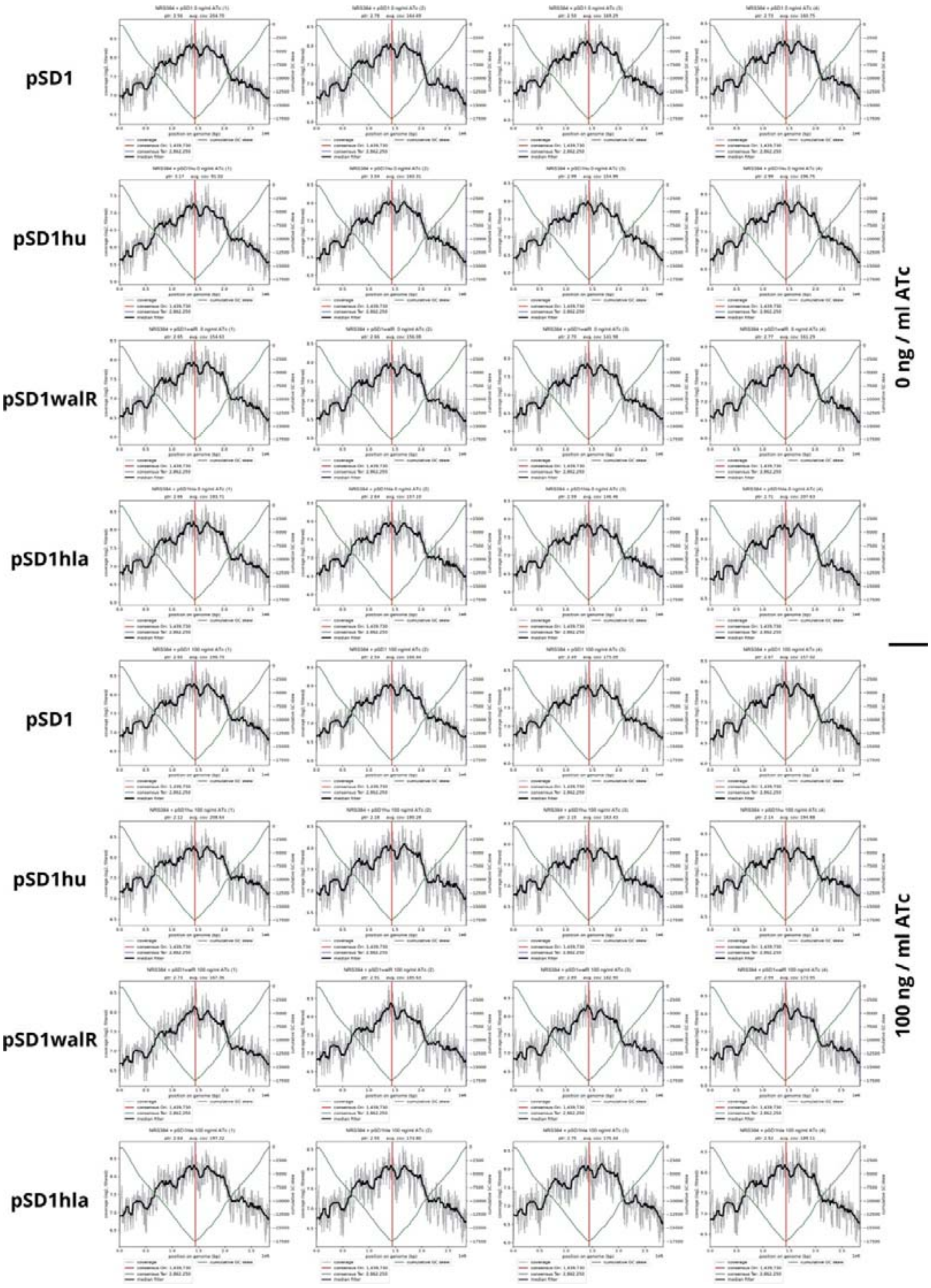

***Figure S9. The effect of walR, hup, and hla on Ori:Ter ratio.*** *Graphs show log2- transformed coverage values (y axis) at relative positions across the genome (x axis). Cumulative GC-skew is plotted on the second Y axis. The calculated position of the origin of replication (consensus ori) is marked in red. Four replicates of each condition are shown. Ptr is the peak to trough ratio, which is equal to the Ori:Ter ratio. Graphs and calculations were made using the iRep R package (https://github.com/christophertbrown/iRep).*

### Table S5. Strains and plasmids

| ***Escherichia coli* strains** | **Description** | **Reference / Supplier** |
|---|---|---|
| IM08B | DH10BΔ*dcm*. Expresses CC8 adenine methylation profile for direct transformation of NRS384. | [1] |
| Rosetta 2(DE3) | B-strain. $F^-$ *ompT hsdSB* ($r_B^-$ $m_B^-$) *gal dcm* (DE3); IPTG-inducible T7 RNA polymerase. For protein production. | Novagen |
| ***Staphylococcus aureus* strains** | | |
| RN4220 | ST8; CC8; chemically mutagenized derivative of 8325-4, transformable with *E. coli* DNA; premature stop codon in both *hsdR* and *sauUSI*. | [2] |
| NRS384 | USA300-14 clone. Tetracycline resistant. | BEI resources |
| NRS384 Δ*spa* | Clean deletion of protein A from TTG to TAA. | This study |
| NRS384 Δ*yycHI* | Deletion of WalKR positive regulators *yycHI*. Clean deletion from codon 5 of *yycH* to the TAA of *yycI*. | [3] |
| NRS384 WalR$_{T101A}$ | The mutation introduces a T->A amino acid change at residue 101. Serine threonine kinase phosphorylation site. Exhibits reduced WalKR activity. | This study |
| NRS384 WalK$_{Y32C}$ | The mutation introduces a Y->C amino acid change at residue 32 (first transmembrane domain) of WalK. Exhibits increased WalKR activity. | This study |
| NRS384 WalK$_{T389A}$ | The mutation introduces a T->A amino acid change at residue 389 (within the HE/DXXT/N of HisKA histidine kinases) predicted to prevent the desphosphorylation of WalR. Exhibits increased WalKR activity. Unstable. | This study |
| NRS384 *walR*-SmBIT | Small-Bit split luciferase tag incorporated onto the C-terminus of WalR (native location) | This study |
| NRS384 walK-LgBIT | Large-Bit split luciferase tag incorporated onto the C-terminus of WalK (native location) | This study |
| NRS384 $walR^{FLAG}$ | FLAG-tag introduced on the C-terminus of WalR in the Wt strain. | [3] |
| **Plasmids** | | |
| pIMAY-Z | Allelic exchange plasmid. Cm(R) | [1] |
| pET28(a) | IPTG inducible protein production plasmid. Amp(R) | Novagen |
| pET21d | IPTG inducible protein production plasmid. Kan(R) | |
| pRAB11FLAG | Anhydrotetracycline inducible E. coli / S. aureus shuttle vector. pC194 rep. C-terminal 1x FLAG. Amp(R) Cm(R) | This study and [4] |
| pIMC8-YFP | Enhanced YFP reporter plasmid. Cm(R) | [3] |
| pSD1 | CRISPRi knockdown plasmid. Amp(R) Cm(R) | [5] |
| pIMK1-LUX | Bacterial luciferase reporter plasmid. pSK41 low copy number replicon. Kan(R) | This study |

| | | |
|---|---|---|
| pSmBIT | pRAB11 backbone (pC194 replicon) with SmBIT split luciferase. Amp(R) Cm(R). | This study |
| pLgBIT | pCN34 backbone (pT181 replicon). TetR/tetO ex pRAB11 introduced. LgBIT split luciferase. Kan(R). | This study |
| pLOW | pSK41 low copy number replicon for IPTG inducible expression in S. aureus. Amp(R) Cm(R). | [6] |
| pCN34 | S. aureus-E. coli shuttle vector, pT181-cop-wt repC ColE1 Kan(R). | [7] |
| **Allelic exchange constructs for *S. aureus*.** | | |
| pIMAY-Z *Δspa* | For construction of protein A deficient strain by allelic exchange. | This study |
| pRAB11: *saeR*$^{FLAG}$ | For construction of WalR T101A mutation by allelic exchange. | This study |
| pIMAY-Z *walK*$^{Y32C}$ | For construction of WalK Y32C mutation by allelic exchange. | This study |
| pIMAY-Z *walK*$^{T389A}$ | For construction of WalK T389A mutation by allelic exchange. | This study |
| pIMAY-Z *walR*(SmBIT) | For construction of a SmBIT C-terminal WalR by allelic exchange. | |
| pIMAY-Z walR(LgBIT) | For construction of a LgBIT C-terminal WalK by allelic exchange. | |
| **ChIP-seq response regulator overexpression constructs for *S. aureus*.** | | |
| pRAB11: *walR*$^{FLAG}$ | ATc inducible plasmid containing of C-terminally FLAG-tagged WalR, for ChIP-seq. | This study |
| pRAB11: v*raR*$^{FLAG}$ | ATc inducible plasmid containing of C-terminally FLAG-tagged VraR, for ChIP-seq. | This study |
| pRAB11: *hptR*$^{FLAG}$ | ATc inducible plasmid containing of C-terminally FLAG-tagged HptR, for ChIP-seq. | This study |
| pRAB11: *saeR*$^{FLAG}$ | ATc inducible plasmid containing of C-terminally FLAG-tagged SaeR, for ChIP-seq. | This study |
| **YFP reporter plasmids for *S. aureus*.** | | |
| pIMC8: P*sasD*-YFP | Native *sasD* driven expression of YFP. | This study |
| pIMC8: P*sle1*-YFP | Native *sle1* driven expression of YFP. | This study |
| pIMC8: P*602*-YFP | Native *SAUSA300_0602* driven expression of YFP. | This study |
| pIMC8: P*ltaS*-YFP | Native *ltaS* driven expression of YFP. | This study |
| pIMC8: P*ssaA1*-YFP | Native *ssaA1* driven expression of YFP. | This study |
| pIMC8: P*isaA*-YFP | Native *isaA* driven expression of YFP. | [3] |
| pIMC8: P$sasD_{ccc}$-YFP | Native *sasD* driven expression of YFP. Mutation in the putative WalR binding site. | |
| pIMC8: P$sle1_{ccc}$-YFP | Native *sle1* driven expression of YFP. Mutation in the putative WalR binding site. | This study |
| pIMC8: P$602_{ccc}$-YFP | Native *SAUSA300_0602* driven expression of YFP. Mutation in the putative WalR binding site. | This study |
| pIMC8: P$ltaS_{ccc}$-YFP | Native *ltaS* driven expression of YFP. Mutation in the putative WalR binding site. | This study |

| | | |
|---|---|---|
| pIMC8: P*ssaA1*$_{cccI}$-YFP | Native *ssaA1* driven expression of YFP. Mutation in the putative WalR binding site (cccI). | This study |
| pIMC8: P*ssaA1*$_{cccII}$-YFP | Native *ssaA1* driven expression of YFP. Mutation in the putative WalR binding site (cccII). | This study |
| pIMC8: P*isaA*$_{cccI}$-YFP | Native *isaA* driven expression of YFP. Mutation in the putative WalR binding site (cccI). | [3] |
| pIMC8: P*isaA*$_{cccII}$-YFP | Native *isaA* driven expression of YFP. Mutation in the putative WalR binding site (cccII). | [3] |
| **Bacterial LUX reporter plasmids for *S. aureus*.** | | |
| pIMK1-LUX: P*sasD* | Native *sasD* driven expression of LUX. | This study |
| pIMK1-LUX: P*sle1* | Native *sle1* driven expression of LUX. | This study |
| pIMK1-LUX: P*602* | Native *SAUSA300_0602* driven expression of LUX. | This study |
| pIMK1-LUX: P*ltaS* | Native *ltaS* driven expression of LUX. | This study |
| pIMK1-LUX: P*ssaA1* | Native *ssaA1* driven expression of LUX. | This study |
| pIMK1-LUX: P*isaA* | Native *isaA* driven expression of LUX. | This study |
| pIMK1-LUX: P*sasD*$_{ccc}$ | Native *sasD* driven expression of LUX. Mutation in the putative WalR binding site. | This study |
| pIMK1-LUX: P*sle1*$_{ccc}$ | Native *sle1* driven expression of LUX. Mutation in the putative WalR binding site. | This study |
| pIMK1-LUX: P*602*$_{ccc}$ | Native *SAUSA300_0602* driven expression of LUX. Mutation in the putative WalR binding site. | This study |
| pIMK1-LUX: P*ltaS*$_{ccc}$ | Native *ltaS* driven expression of LUX. Mutation in the putative WalR binding site. | This study |
| pIMK1-LUX: P*ssaA1*$_{cccII}$ | Native *ssaA1* driven expression of LUX. Mutation in the putative WalR binding site (cccII). | This study |
| pIMK1-LUX: P*isaA*$_{cccI}$ | Native *isaA* driven expression of LUX. Mutation in the putative WalR binding site (cccI). | This study |
| pIMK1-LUX: P*hup* | Native *hup* driven expression of LUX. | This study |
| pIMK1-LUX: P*spa* | Native *spa* driven expression of LUX. | This study |
| pIMK1-LUX: P*prs* | Native *prs* driven expression of LUX. | This study |
| pIMK1-LUX: P*rplK* | Native *rplK* driven expression of LUX. | This study |
| pIMK1-LUX: P*dnaA* | Native *dnaA* driven expression of LUX. | This study |
| pIMK1-LUX: P*dnaD* | Native *dnaD* driven expression of LUX. | This study |
| pIMK1-LUX: P*tarF* | Native *tarF* driven expression of LUX. | This study |
| pIMK1-LUX: P*tagG* | Native *tagG* driven expression of LUX. | This study |
| pIMK1-LUX: P*hup*$_{ccc}$ | Native *hup* driven expression of LUX. Mutation in the putative WalR binding site. | This study |
| pIMK1-LUX: P*spa*$_{ccc}$ | Native *spa* driven expression of LUX. Mutation in the putative WalR binding site. | This study |
| pIMK1-LUX: P*prs*$_{ccc}$ | Native *prs* driven expression of LUX. Mutation in the putative WalR binding site. | This study |

| | | |
|---|---|---|
| pIMK1-LUX: P*rplK*$_{ccc}$ | Native *rplK* driven expression of LUX. Mutation in the putative WalR binding site. | This study |
| pIMK1-LUX: P*dnaA*$_{cccI}$ | Native *dnaA* driven expression of LUX. Mutation in the putative WalR binding site closest to the start codon. | This study |
| pIMK1-LUX: P*dnaA*$_{cccII}$ | Native *dnaA* driven expression of LUX. Mutation in second putative WalR binding site. (See Fig 7C) | This study |
| pIMK1-LUX: P*dnaA*$_{cccIII}$ | Native *dnaA* driven expression of LUX. Mutation in third putative WalR binding site. (See Fig 7C) | This study |
| pIMK1-LUX: P*dnaD*$_{cccI}$ | Native *dnaD* driven expression of LUX. Mutation in the putative WalR binding site closest to the start codon.. | This study |
| pIMK1-LUX: P*dnaD*$_{cccII}$ | Native *dnaD* driven expression of LUX. Mutation in the second putative WalR binding site. | This study |
| pIMK1-LUX: P*tarF*$_{ccc}$ | Native *tarF* driven expression of LUX. Mutation in the putative WalR binding site. | This study |
| pIMK1-LUX: P*tagG*$_{ccc}$ | Native *tagG* driven expression of LUX. Mutation in the putative WalR binding site. | This study |
| **Protein overexpression in *E. coli.*** | | |
| pET28(a): *walR* | For production of recombinant WalR in *E. coli*. | This study |
| pET28(a): *vraR* | For production of recombinant VraR in *E. coli.* | This study |
| **CRISPRi plasmids for gene knockdown in *S. aureus*.** | | |
| pSD1:*walR* | CRISPRi knockdown of *walR* in *S. aureus*. | This study |
| pSD1:*hup* | CRISPRi knockdown of *hup* in *S. aureus.* | This study |
| pSD1:*hla* | CRISPRi knockdown of *hla* in *S. aureus.* | This study |
| **Split luciferase plasmids for protein:protein intereaction in *S. aureus.*** | | |
| pSmBIT*walR* | C-terminally tagged WalR with SmBIT luciferase fragment | This study |
| pSmBIT*walR*$_{D53A}$ | C-terminally tagged WalR$_{D53A}$ with SmBIT luciferase fragment | This study |
| pSmBIT*walR*$_{D53E}$ | C-terminally tagged WalR$_{D53E}$ with SmBIT luciferase fragment | This study |
| pSmBIT*walR*$_{T101A}$ | C-terminally tagged WalR$_{T101A}$ with SmBIT luciferase fragment | This study |
| pSmBIT*walR*$_{D53A/T101A}$ | C-terminally tagged WalR$_{D53A/101A}$ with SmBIT luciferase fragment | This study |
| pLgBIT*walK* | C-terminally tagged WalK with LgBIT luciferase fragment | This study |
| pLgBIT*walK*$_{Y32C}$ | C-terminally tagged WalK$_{Y32C}$ with LgBIT luciferase fragment | This study |
| pLgBIT*walK*$_{T389A}$ | C-terminally tagged WalK$_{T389A}$ with LgBIT luciferase fragment | This study |
| pLgBIT*walK*$_{G223D}$ | C-terminally tagged WalK$_{G223D}$ with LgBIT luciferase fragment | This study |
| pLgBIT*walK*$_{H385A}$ | C-terminally tagged WalK$_{H385A}$ with LgBIT luciferase fragment | This study |

### Supplementary Table 6: Primers

| | EMSA | |
|---|---|---|
| **Primer** | **Oliognucleotide 5'-3'** | **Name** |
| LS237 | /5Cy5/CTTTTGTAATCGTATGATAATGTAAATGTAATCAAATTGTAATATAAGGGGACAAGACAATGAAAAAATT | *sasD* EMSA cy5 fwd |
| LS238 | CTTTTGTAATCGTATGATAATGTAAATGTAATCAAATTGTAATATAAGGGGACAAGACAATGAAAAAATT | *sasD* EMSA fwd |
| LS239 | AATTTTTTCATTGTCTTGTCCCCTTATATTACAATTTGATTACATTTACATTATCATACGATTACAAAAG | *sasD* EMSA rev |
| LS234 | /5Cy5/TTTGATGATACAGTATATGATTTTTTTGTAATCATAATGTCATCAAACATCAACCTATTATACATAATAA | *sle1* EMSA cy5 fwd |
| LS235 | TTTGATGATACAGTATATGATTTTTTTGTAATCATAATGTCATCAAACATCAACCTATTATACATAATAA | *sle1* EMSA fwd |
| LS236 | TTATTATGTATAATAGGTTGATGTTTGATGACATTATGATTACAAAAAAATCATATACTGTATCATCAAA | *sle1* EMSA rev |
| LS243 | /5Cy5/GAAATATAAAAGTAGTAATTATTATTTGTAATATAATTGTAATATGACTGTTGTTTTAGAAATGATTGTT | *P602* EMSA fwd cy5 |
| LS244 | GAAATATAAAAGTAGTAATTATTATTTGTAATATAATTGTAATATGACTGTTGTTTTAGAAATGATTGTT | *P602* EMSA fwd |
| LS245 | AACAATCATTTCTAAAACAACAGTCATATTACAATTATATTACAAATAATAATTACTACTTTTATATTTC | *P602* EMSA rev |
| LS240 | /5Cy5/ATATTTATTAATTGAGCTATGCTTATTATTACAATTTGATTACAAATTTTAAATTTGTTAATTGAATGAT | *ltaS* EMSA fwd cy5 |
| LS241 | ATATTTATTAATTGAGCTATGCTTATTATTACAATTTGATTACAAATTTTAAATTTGTTAATTGAATGAT | *ltaS* EMSA fwd |
| LS242 | ATCATTCAATTAACAAATTTAAAATTTGTAATCAAATTGTAATAATAAGCATAGCTCAATTAATAAATAT | *ltaS* EMSA rev |
| LS246 | /5Cy5/AAAACTGAATGTTTCGAAAATAAGTCTGTTACAAATTTGTAATATTACTGAAAATTCTAAATGTATATTT | *ssaA* EMSA fwd Cy5 |
| LS247 | AAAACTGAATGTTTCGAAAATAAGTCTGTTACAAATTTGTAATATTACTGAAAATTCTAAATGTATATTT | *ssaA* EMSA fwd |
| LS248 | AAATATACATTTAGAATTTTCAGTAATATTACAAATTTGTAACAGACTTATTTTCGAAACATTCAGTTTT | *ssaA* EMSA rev |
| LS249 | /5Cy5/AAATAACACTTGATATTGTAATGTTTTGTAAAGAAAGTGTAATTTACTGGCTGGTTTTTTGTGATATAGT | *isaA* EMSA fwd Cy5 |
| LS250 | AAATAACACTTGATATTGTAATGTTTTGTAAAGAAAGTGTAATTTACTGGCTGGTTTTTTGTGATATAGT | *isaA* EMSA fwd |
| LS251 | ACTATATCACAAAAAACCAGCCAGTAAATTACACTTTCTTTACAAAACATTACAATATCAAGTGTTATTT | *isaA* EMSA rev |
| LS267 | /5Cy5/CATTAGGAGGAAATTATTTGCATCGGACTCGAGTATGAGCTACGTACCAAATATTATTTCATCATTTCTA | VraR EMSA *ltaS* fwd Cy5 |
| LS268 | CATTAGGAGGAAATTATTTGCATCGGACTCGAGTATGAGCTACGTACCAAATATTATTTCATCATTTCTA | VraR EMSA *ltaS* fwd |
| LS269 | TAGAAATGATGAAATAATATTTGGTACGTAGCTCATACTCGAGTCCGATGCAAATAATTTCCTCCTAATG | VraR EMSA *ltaS* rev |
| LS91 | ACTGTGACG*GGATCC*ATGGCTAGAAAAGTTGTTGTAGTTGATG | *walR* pET28 F |
| LS92 | CATCGACTTAG*AAGCTT*TACTCATGTTGTTGGAGGAAATATCCAAC | *walR* pET28 R |
| IM423 | TTTGTTTAACTTTAAGAAGGAGATATACCATGACGATTAAAGTATTGTTTGTGGATG | *vraR* pET21 F |

| | | |
|---|---|---|
| IM424 | GGCTTTGTTAGCAGCCGGATCCTAATGGTGATGATGGTGGTGATGGTGATGATGTTGAATTAAATTATGTTGGAATGC ATAG | *vraR* pET21 R |
| IM395 | GGTATATCTCCTTCTTAAAGTTAAACAAAATTATTTC | pET INV F |
| IM396 | GATCCGGCTGCTAACAAAGCC | pET INV R |

**_S. aureus_ mutants**

| Primer | Oliognucleotide 5'-3' | Name |
|---|---|---|
| IM31 | CCTCACTAAAGGGAACAAAAGCTGGGTACCCGTCCATTTTCTTTAAAATGTATGAACC | *walR*$^{T101A}$ AF |
| IM231 | aGcaACATAGTCATCTGCACCTAGTTC | *walR*$^{T101A}$ BR |
| IM232 | TTAGAACTAGGTGCAGATGACTATGTtgCtAAACCGTTTAGTACGCGTGAATTAATCG | *walR*$^{T101A}$ CF |
| IM233 | AGAACTAGGTGCAGATGACTATGTtg | *walR*$^{T101A}$ con F |
| IM181 | TTTTTCAAGGTTATTTGTAAAATATAACCC | *walR*$^{T101A}$ con R |
| IM107 | CCTCACTAAAGGGAACAAAAGCTGGGTACCATGGCTAGAAAAGTTGTTGTAG | *walK*$^{Y32C}$ AF |
| IM7 | CCTCACTAAAGGGAACAAAAGCTGGGTACCAGGTCGAAACGAATGAAGTGGCTAAAAC | *walK*$^{T389A}$ AF |
| IM121 | agcACGTAACTCATGTGATACATTGG | *walK*$^{T389A}$ BR |
| IM122 | CAATGTATCACATGAGTTACGTgctCCTTTAACTTCTATGAATAGTTACATTGAAGC | *walK*$^{T389A}$ CF |
| IM10 | CGACTCACTATAGGGCGAATTGGAGCTCCTCCTTATTATTCATCCCAATCACCGTC | *walR*$^{T101A}$/*walK*$^{Y32C}$/$^{T389A}$DR |
| IMT275 | CCTCACTAAAGGGAACAAAAGCTGGGTACCAATTCATATGGATGACGCGCAGC | Delta *spa* AF |
| IMT353 | CAAATTAATACCCCCTGTATGTATTTG | Delta *spa* BR |
| IMT354 | CAAATACATACAGGGGGTATTAATTTGTAAAAACAAACAATACACAACGATAGATATC | Delta *spa* CF |
| IMT278 | CGACTCACTATAGGGCGAATTGGAGCTCATTACTTGTGGCAGCTAACACTGC | Delta *spa* DR |
| IM1516 | GGTTATAGACTTTTTGAAGAAATTCTATAAAGGTCGAAACGAATGAAGTGGCTAAAAC | *walR*-SmBIT-*walK* CF |
| IM1517 | TTATAGAATTTCTTCAAAAAGTCTATAACCTG | *walR*-SmBIT-*walK* BR |
| IM1518 | GTTGTTTAGAGTAACTATAAACAGTTAAATGAATAATAAGGAGCATATTAAATCTGTC | *walK*-LgBIT-*yycH* CF |
| IM1519 | TTAACTGTTTATAGTTACTCTAAACAACATAGATCC | *walK*-LgBIT-*yycH* BR |
| IM44 | CGACTCACTATAGGGCGAATTGGAGCTCTTGTTAATTTTACGTAATCGTGGCGATC | *walK*-LgBIT-*yycHI* DR |
| IM1368 | AGCCCGATAATTTGCATACCAATG | *walK*-LgBIT con R |

**ChIP-seq**

| Primer | Oliognucleotide 5'-3' | Name |
|---|---|---|
| IM512 | ATAT*GGTACC*GATTACAAAGATGATGATGACAAATAAAATAGTCAAAAGCCTCCGGTCGGAGGCTTTTGACTGAATTCA CTGGCCGTCGTTTTAC | pRAB11 FLAG tonB (KpnI) F |
| IM513 | ATAT*GGTACC*TTTCCAATTCCTCCTCATCATACTCTATCAATGATAGAGAGC | pRAB11 RBS (KpnI) R |
| IM514 | GATTACAAAGATGATGATGACAAATAAAATAGTC | INV pRAB11 FLAG F |
| IM515 | TTTCCAATTCCTCCTCATCATACTCTATC | INV pRAB11 FLAG R |
| IM516 | CATTGATAGAGTATGATGAGGAGGAATTGGAAATGGCTAGAAAAGTTGTTGTAGTTG | WalR-FLAG F |
| IM517 | CTATTTTATTTGTCATCATCATCTTTGTAATCCTCATGTTGTTGGAGGAAATATCC | WalR-FLAG R |
| IM518 | CATTGATAGAGTATGATGAGGAGGAATTGGAAATGACCCACTTACTGATCGTGG | SaeR-FLAG F |
| IM519 | CTATTTTATTTGTCATCATCATCTTTGTAATCTCGGCTCCTTTCAAATTTATATCC | SaeR-FLAG R |
| IM520 | CATTGATAGAGTATGATGAGGAGGAATTGGAAATGACGATTAAAGTATTGTTTGTGGATG | VraR FLAG F |
| IM521 | CTATTTTATTTGTCATCATCATCTTTGTAATCTTGAATTAAATTATGTTGGAATGCATAG | VraR-FLAG R |
| IM522 | CATTGATAGAGTATGATGAGGAGGAATTGGAAATGTTTAAGGTAGTTATTTGTGATGATG | HptR-FLAG F |
| IM523 | CTATTTTATTTGTCATCATCATCTTTGTAATCTTTTGCTTGCTTACAATAATCACTTGG | HptR-FLAG R |

**YFP-LUX**

| Primer | Oliognucleotide 5'-3' | Name | Location |
|---|---|---|---|
| IM1 | GGTACCCAGCTTTTGTTCCCTTTAGTGAGG | pIMC8-YFP F | |
| IM385 | TGATTAACTTTATAAGGAGGAAAAACATATG | pIMC8-YFP-R | |
| IM1127 | CCTCACTAAAGGGAACAAAAGCTGGGTACCTATAACTTAATATAATTGAGGTGGAGCATC | *sasD* YFP F | |
| IM1107 | ATGTTTTTCCTCCTTATAAAGTTAATCATGTCTTGTCCCCTTATATTACAATTTG | *sasD* YFP R | |
| IM1108 | CCTCACTAAAGGGAACAAAAGCTGGGTACCTTAGAAAAATCAAATTCAGATGCAGTAAAG | *sle1* YFP F | |
| IM1109 | ATGTTTTTCCTCCTTATAAAGTTAATCATTTAAAATCCTCCTCTTGCTTAACTTTCC | *sle1* YFP R | |
| IM1129 | CCTCACTAAAGGGAACAAAAGCTGGGTACCCAAGAATCAAGTGCATTCCCATTTGTTG | *P602* YFP F | |
| IM1110 | ATGTTTTTCCTCCTTATAAAGTTAATCAAAGCACTCTCTCCTTTTATTTATATCG | *P602* YFP R | |
| IM1111 | CCTCACTAAAGGGAACAAAAGCTGGGTACCTAATATTTGGTACGTAGCTCATACTCG | *ltaS* YFP F | |
| IM1112 | ATGTTTTTCCTCCTTATAAAGTTAATCAGATTCTTTCCCCCGTTATTTAGATAATAAATC | *ltaS* YFP R | |
| LS371 | CCTCACTAAAGGGAACAAAAGCTGGGTACCCCAAATACCAAAGCTTTCATAATC | *ssaA* YFP F | |
| LS372 | ATGTTTTTCCTCCTTATAAAGTTAATCATTTAAAAATATCCTCCTAAAAATTTTAAATC | *ssaA* YFP R | |
| IM1128 | CCTCACTAAAGGGAACAAAAGCTGGGTACCTAACAGTATGTTTTTTGAAAATATGAGACC | *isaA* YFP F | |

| | | | |
|---|---|---|---|
| IM364 | ATGTTTTTCCTCCTTATAAAGTTAATCAAGTAAAAAATCCTCCAGTAATAATTG | *isaA* YFP R | |
| IMT300 | CCTCACTAAAGGGAACAAAAGCTGGGTACCATCCAGATTTAATAATAGGATGGTTAGG | *hpt* YFP F | |
| IMT301 | ATGTTTTTCCTCCTTATAAAGTTAATCACTCTGTCACCTCAATCATTTTCG | *hpt* YFP R | |
| IM1113 | ATGTTTTTCCTCCTTATAAAGTTAATCATGTCTTGTCCCCTTATATT**GGG**ATTTGATTAC | *sasD* ccc R | |
| IM1114 | GATGATACAGTATATGATTTTTT**CCC**AATCATAATGTCATCAAACATC | *sle1* ccc F | |
| IM1115 | **GGG**AAAAAATCATATACTGTATCATCAAAT | *sle1* ccc R | |
| IM1118 | GAAATATAAAAGTAGTAATTATTATT**CCC**AATATAATTGTAATATGACTGTTGTTTTAG | *P602* ccc F | |
| IM1119 | **GGG**AATAATAATTACTACTTTTATATTTCACTTATC | *P602* ccc R | |
| IM1116 | GAGCTATGCTTATTATTACAATTTGATT**GGG**AATTTTAAATTTGTTAATTGAATG | *ltaS* ccc F | |
| IM1117 | **CCC**AATCAAATTGTAATAATAAGCATAGCTC | *ltaS* ccc R | |
| LS375 | **GGG**AAGCACACAAGGACGCTAAT | *ssaA* cccI F | |
| LS376 | ATTAGCGTCCTTGTGTGCTT**CCC**TAACGTTTTGTAATTTTTTGCTAATATC | *ssaA* cccI R | closest to the *ssaA* start |
| IM1120 | CAGTAATATTACAAATTTGT**GGG**GACTTATTTTCGAAACATTCAG | *ssaA* cccII F | |
| IM1121 | **CCC**TACAAATTTGTAATATTACTGAAAATTCTAAATG | *ssaA* cccII R | furthest from the *ssaA* start |
| IM1122 | ACACTTGATATTGTAATGTTT**CCC**AAAGAAAGTGTAATTTACTGGCTGG | *isaA* cccI F | |
| IM1123 | **GGG**AAACATTACAATATCAAGTGTTATTTG | *isaA* cccI R | closest to the *isaA* start |
| IM1124 | CAGATATATTACAGCTATGTA**GGG**AAAATACAATCTGTAATATTACGAAAGC | *isaA* cccII F | |
| IM1125 | **CCC**TACATAGCTGTAATATATCTGACATGTAAC | *isaA* cccII R | furthest from the *isaA* start |

| **Primer** | **LUX Oliognucleotide 5'-3'** | **Name** | **Location** |
|---|---|---|---|
| IM1216 | ATAT*GTCGAC*TATAACTTAATATAATTGAGGTGGAGCATC | *sasD* LUX F | |
| IM248 | CATTGTCTTGTCCCCTTATATTACAATTTG | *sasD* LUX R | |
| IM1295 | ATAT*GTCGAC*TTAGAAAAATCAAATTCAGATGCAGTAAAG | *sle1* LUX F | |
| IM1296 | CACTTTAAAATCCTCCTCTTGCTTAACTTTCC | *sle1* LUX R | |
| IM1294 | ATAT*GTCGAC*CAAGAATCAAGTGCATTCCCATTTGTTG | 0602 LUX F | |

| | | |
|---|---|---|
| IM1062 | CATAAGCACTCTCTCCTTTTATTTATATCG | 0602 LUX R |
| IM1218 | ATAT*GTCGAC*TAATATTTGGTACGTAGCTCATACTCG | *ltaS* LUX F |
| IM1219 | CATGATTCTTTCCCCCGTTATTTAGATAATAAATC | *ltaS* LUX R |
| IM1297 | ATAT*GTCGAC*CTTACATCCTCACATATACAAATATATTG | *ssaA* LUX F |
| IM1298 | CATTTTAAAAATATCCTCCTAAAAATTTTAAATC | *ssaA* LUX R |
| IM1220 | ATAT*GTCGAC*TAACAGTATGTTTTTTGAAAATATGAGACC | *isaA* LUX F |
| IM1221 | CATAGTAAAAAATCCTCCAGTAATAATTG | *isaA* LUX R |
| IM1222 | ATAT*GTCGAC*TTCTGAGCAATGACGTGCAACTAG | *walR* LUX F |
| IM32 | CATTTGCATAAACCTCTTTTCTTAAATC | *walR* LUX R |
| IM1217 | CATTGTCTTGTCCCCTTATATT**GGG**ATTTGATTAC | *sasD* ccc LUX R |
| LS443 | TACGCGTCGACACACGATGCGAGCAATCAAAT | *dnaA* LUX F |
| IM1746 | CATAAATTTACTCCATCTTTCTTAAATTTTTAAG | *dnaA* LUX R |
| IM1285 | ATAT*GTCGAC*CAATTAGGTGCAGATGTTAAGGTTG | *prs* LUX F |
| IM1288 | CATTTATAGTCCTCCAATTATTTACTTACG | *prs* LUX R |
| LS460 | TACGC*GTCGAC*CATATGTATTAAACCGCTTTCATTATAAAAAATATCTCTATATTTTATCTG | *spa* LUX F |
| LS463 | CAAATTAATACCCCCTGTATGTATTTGTAAAGTCATCATAATATAACG | *spa* LUX R |
| IM1289 | ATAT*GTCGAC*TGCCGATCCTATGACATTTCTAGG | *hup* LUX F |
| IM1285 | CATTAGACATTCACCTCCTGAGG | *hup* LUX R |
| LS451 | TACGC*GTCGAC*GATGAGAAGCAGGACTATACAATGAG | *tarF* LUX F |
| LS452 | CATGTCGTACCTCCGACGTG | *tarF* LUX R |
| LS457 | TACGC*GTCGAC*GAAGATTTTGTTATTATCAGAGTGGG | *tagG* LUX F |
| LS458 | CATTCCATTAAACCACACTTTCAAATG | *tagG* LUX R |
| LS471 | TACGC*GTCGAC*GCTGGGCGATTTATTCTTGG | *dnaD* LUX F |
| LS472 | CATGTTCGTGCCCCCTTTTAA | *dnaD* LUX R |
| IM1745 | ATAT*GTCGAC*CTCACTAACAGATACTCTATAGAAGG | *dnaA* LUX F |
| IM1746 | CATAAATTTACTCCATCTTTCTTAAATTTTTAAG | *dnaA* LUX R |

| | | | |
|---|---|---|---|
| IM1734 | ATAT*GTCGAC*ATTAACAATTAAAGTTATTAAACTAACCAAAAG | *rplK* LUX F | |
| IM1735 | CACGATGTGCACCTCCTTGATATCG | *rplK* LUX R | |
| LS464 | CAAATTAATACCCCCTGTA**CCC**ATTTGTAAAGTCATCATAATATAACG | *spa* ccc LUX R | |
| IM1291 | CCACATAAATGATTGCGATTTA**CCC**AATGCTTGTGTTAAAGTTTTACC | *hup* ccc LUX CF | |
| IM1290 | **GGG**TAAATCGCAATCATTTATGTGGCAAAAACG | *hup* ccc LUX BR | |
| LS449 | GGATAACGATTTTT**CCC**TATGTTAAAGTGGTAC | *tarF* ccc LUX CF | |
| LS448 | **GGG**AAAAATCGTTATCCATTCATAACGTATG | *tarF* ccc LUX BR | |
| LS455 | GATAGCATCCA**CCC**ATAGTGATAGTATTTACAAC | *tagG* ccc LUX CF | |
| LS454 | **GGG**TGGATGCTATCAAAATTATTGTCTAGTTC | *tagG* ccc LUX BR | |
| LS467 | CCATAAAGTTCAT**CCC**TAAAATCTAGTGTTAAAAAATAC | *dnaD* cccI LUX CF | |
| LS466 | **GGG**ATGAACTTTATGGATAATCAGATGAACTA | *dnaD* cccI LUX BR | closest to the start |
| LS469 | GTTATTTAAAATT**CCC**TTGCTACTAGTTAAATATATTAAG | *dnaD* cccII LUX CF | |
| LS468 | **GGG**AATTTTAAATAACAGTATTTTTTAACACTAGATTTTAAC | *dnaD* cccII LUX BR | furthest from the start |
| IM1743 | **CCC**GAATATGTTGCTAAAAATAGTATAACTTTGTG | *dnaA* ccc LUX CF | |
| IM1744 | GTTATACTATTTTTAGCAACATATTC**GGG**GGTATTTGACATATAGAGAACTGAAAAAG | *dnaA* ccc LUX BR | |
| LS144 | GAGTATTGATTTTTTAATTAGAAAAG**CCC**TAAAATTATGTGGTCGCGC | *rplK* ccc LUX CF | |
| LS143 | **GGG**CTTTTCTAATTAAAAAATCAATACTCTTTTTTA | *rplK* ccc LUX BR | |
| LS140 | TGGCTAGGATAAAAGGATAATCCTA**CCC**AATATTAATGTAATCTTTATG | *prs* ccc LUX CF | |
| LS139 | **GGG**TAGGATTATCCTTTTATCCTAGCCATTTTAAATAC | *prs* ccc LUX BR | |
| LS441 | **CCC**TATTTTTGTGTATAACTTAAAAATTTAAGAAAGATG | *dnaA* ccc I LUX CF | closest to the start |
| LS440 | GTTATACACAAAAATA**GGG**GCCGTGGATAACTTC | *dnaA* ccc I LUX BR | |
| LS439 | **CCC**TATTAACTTGTGGATAATTATTAACATGGTG | *dnaA* ccc II LUX CF | |
| LS438 | CCACAAGTTAATA**GGG**ATCGGTGGATAAC | *dnaA* ccc II LUX BR | |
| IM1744 | GTTATACTATTTTTAGCAACATATTC**GGG**GGTATTTGACATATAGAGAACTGAAAAAG | *dnaA* ccc III LUX CF | |
| IM1743 | **CCC**GAATATGTTGCTAAAAATAGTATAACTTTGTG | *dnaA* ccc III LUX BR | furthest from the start |
| IM1241 | ATAT*AGATCT*ATATAGTTTTTGTATACGGTATTCATTCATG | pLOW rep F | |
| IM1242 | ATAT*GCATGC*TCAACTTTGCAACAGAACC | pLOW rep R | |

**RT-qPCR**

| Primer | Oliognucleotide 5'-3' | Name |
|---|---|---|
| IM1026 | GGCTCTATGAAAGCAGCAGATA | Hla RT-qPCR F |
| IM1027 | CTGTAGCGAAGTCTGGTGAAA | Hla RT-qPCR R |
| IM1020 | CGCAGGCGATTTTACCATTA | GyrB RT-qPCR F |
| IM1021 | ATGCTGGAACTTTACTTGCTGG | GyrB RT-qPCR R |
| IM1153 | GAACTAGGTGCAGATGACTATGT | WalR RT-qPCR F |
| IM1154 | CTTGTGCTGGTTGTGAGTAATG | WalR RT-qPCR R |
| IM1586 | GGTTTCGGTAACTTTGAGGTACG | Hu RT-qPCR F |
| IM1587 | ATGCTGGAACTTTACTTGCTGG | Hu RT-qPCR R |

**SmBIT/LgBIT**

| Primer | Oliognucleotide 5'-3' | Name |
|---|---|---|
| IM1290 | ATCATTAATTCCTCCTTTTTGTTGACAcTcTATCATTGATAGAGTTATTTGTCAAACTAG | Fix tetO1 F |
| IM1291 | gAgTGTCAACAAAAAGGAGGAATTAATGATG | Fix tetO1 R |
| IM1355 | ATAT*GGTACC*GTTAACAGATCTGAGCTCG | Introduce RBS F |
| IM513 | ATAT*GGTACC*TTTCCAATTCCTCCTCATCATACTCTATCAATGATAGAGAGC | Introduce RBS R |
| IM515 | TTTCCAATTCCTCCTCATCATACTCTATC | INV PCR to clone SmBIT/LgBIT R |
| IM1356 | GTTAACAGATCTGAGCTCGAATTCACTGG | INV PCR to clone SmBIT/LgBIT F |
| IM1360 | GGTTCTAGTGGTGGTGGTGGTTCTGG | INV PCR to clone genes F (IM515) |
| IM1363 | GATAGAGTATGATGAGGAGGAATTGGAAAATGGCTAGAAAAGTTGTTGTAGTTG | WalR pSmBIT F |
| IM1364 | GAACCACCACCACCACTAGAACCCTCATGTTGTTGGAGGAAATATCC | WalR pSmBIT R |
| IM1365 | GATAGAGTATGATGAGGAGGAATTGGAAAATGAAGTGGCTAAAACAACTACAATCC | WalK pLgBIT F |
| IM1366 | GAACCACCACCACCACTAGAACCTTCATCCCAATCACCGTCTTCAATGAC | WalK pLgBIT R |

**CRISPRi**

| Primer | Oliognucleotide 5'-3' | Name |
|---|---|---|

| | | |
|---|---|---|
| IM1559 | *CTA*GTTCATTAGACATTCACCTCCTG | *hup* CRISPRi F |
| IM1560 | *AAC*CAGGAGGTGAATGTCTAATGAAC | *hup* CRISPRi R |
| IM1180 | *CTA*TTAAATTCTAAAATATCAGCAAT | *walR* CRISPRi F |
| IM1181 | *AAC*ATTGCTGATATTTTAGAATTTAA | *walR* CRISPRi R |
| IM1182 | *CTA*CATTAGCGACAGGATTCATTAATA | *hla* CRISPRi F |
| IM1183 | *AAC*TATTAATGAATCCTGTCGCTAATG | *hla* CRISPRi R |

*Itallics*: restriction enzyme sites
Underlined: Tails complementary to plasmid for SLiCE cloning
lowercase: Mutations introduced
**Bold**: WalR binding site mutations
